# SorCS2 controls Wnt signaling homeostasis during neurodevelopment by regulating Ror2 activity and trafficking

**DOI:** 10.64898/2026.10.10.758224

**Authors:** Alena Salašová, Lucie Kuncová, Chika Yokota, Kasper Kjær-Sørensen, Anna Catharina Navis, Stella Nolte, Ondřej Bernatík, Enrique M. Toledo, Josef Lavický, Jan Krivanek, Manuel Varas-Godoy, Emma R. Andersson, Claus Oxvig, Vítězslav Bryja, Ernest Arenas, Anders Nykjær

**Affiliations:** Danish Research Institute of Translational Neuroscience DANDRITE, Nordic EMBL Partnership for Molecular Medicine, and Center of Excellence PROMEMO, 8000 Aarhus C, Denmark; Department of Biomedicine, Aarhus University, 8000 Aarhus C, Denmark; Division of Molecular Neurobiology, Department of Medical Biochemistry and Biophysics, Karolinska Institutet, 17177 Stockholm, Sweden; Department of Molecular Biology and Genetics, Aarhus University, 8000 Aarhus, Denmark; Department of Experimental Biology, Faculty of Science, Masaryk University, 62500 Brno, Czech Republic; Department of Histology and Embryology, Faculty of Medicine, Masaryk University, 62500 Brno, Czech Republic; Centro de Biología Celular y Biomedicina (CEBICEM), Facultad de Ciencias, Universidad San Sebastián, 8580702 Santiago, Chile; Centro Ciencia & Vida, Fundación Ciencia & Vida, 8580702 Santiago, Chile; Department of Cell and Molecular Biology, Karolinska Institute, 17177 Stockholm, Sweden; Department of Biosciences and Nutrition, Karolinska Institute, 14183 Huddinge, Sweden

**Keywords:** Wnt/β-catenin signaling, noncanonical Wnt signaling, Ror2, Wnt5a, Wnt1, SorCS2, Tcf4, dopaminergic circuits, Wnt homeostasis, intracellular sorting

## Abstract

Wnt signaling orchestrates key developmental processes, yet mechanisms restricting its spatial and temporal activity remain incompletely understood. Here we identify SorCS2, a VPS10p-domain sorting receptor, as a conserved regulator of Wnt signaling homeostasis. Integrating proteomics, cell biology, embryology, and vertebrate models, we show that SorCS2 interacts with the noncanonical Wnt receptor Ror2 and controls its trafficking, Wnt5a-induced internalization, and lysosomal degradation. In mouse dopaminergic progenitors, SorCS2 deficiency enhances responsiveness to canonical and noncanonical Wnt ligands, increases Ror2 phosphorylation, and amplifies downstream signaling of both Wnt signaling cascades. Loss of SorCS2 further disrupts the ligand-receptor feedback loop, leading to elevated *Ror2, Wnt5a,* and *Wnt1* expression, accelerated proliferation and neurogenesis, and increased expression of *Tcf4,* a gene implicated in neuropsychiatric disorders. Across vertebrate systems, SorCS2-dependent control of Ror2 signaling is required for anterior-posterior axis elongation, convergent extension, somitogenesis, and brain morphogenesis. These findings define a SorCS2-dependent sorting mechanism that fine-tunes morphogen receptor signaling during vertebrate neurodevelopment.

## Introduction

Wnts (Wingless/Integration)^1, 2^ are highly conserved, secreted lipid-modified glycoproteins essential for embryonic development, tissue homeostasis, and regeneration. Maintaining homeostatic Wnt activity is critical for signaling precision, as both loss and gain of function can have detrimental consequences. Over the past decade, Wnt signaling has become a promising therapeutic target due to its central role in developmental disorders, neurodegeneration, cancer, ciliopathies, or inflammatory diseases. However, the molecular mechanisms governing its activity and specificity remain poorly understood^3, 4, 5, 6, 7^.

Wnt signaling is categorized into two interconnected branches: the canonical Wnt/β-catenin pathway and the noncanonical Wnt signaling pathways. The Wnt/β-catenin pathway is activated by Wnt1/3a ligands binding to a Frizzled (Fzd) receptor and its co-receptor Low-density Lipoprotein Receptor-related Protein 5/6 (Lrp5/6) at the plasma membrane. This interaction leads to Lrp5/6 phosphorylation, recruitment and phosphorylation of Dishevelled (Dvl) proteins, and the subsequent inhibition of β-catenin destruction complex, resulting in the activation of β-catenin and transcription of TCF/LEF-dependent target genes that regulate e.g. proliferation, stemness, and differentiation^4, 5, 6, 7^.

Noncanonical Wnt signaling comprises several pathways with unresolved molecular mechanisms due to their complex interactomes and context-dependent effects. Noncanonical Wnt signaling is recognized for its role in convergent extension, body axis elongation, and neural tube morphogenesis^4, 7, 8, 9^. Wnt5a is the main noncanonical Wnt ligand, and its deficiency causes severe, often lethal developmental defects, affecting organogenesis, neural tube closure, and craniofacial and skeletal formation. The loss of its core receptor, single-pass transmembrane Receptor Tyrosine Kinase-like Orphan Receptor 2 (Ror2), largely phenocopies Wnt5a deficiency^4, 7, 10, 11, 12, 13, 14^.

Wnt5a binding to the extracellular Cysteine-Rich Domain (CRD) of Ror2 induces receptor dimerization (homo- or hetero-) and assembly of multimeric receptor complexes with Frizzled receptors (i.e. Fzd2)^15^ or co-receptors including Ror1, Lrp4, Inactive tyrosine-protein kinase 7 (Ptk7), or Vangl-like protein 2 (Vangl2)^4, 7, 11, 12, 13, 16, 17, 18^. Receptor engagement promotes recruitment of Dvl isoforms and associated kinases, such as Glycogen synthase kinase 3β (GSK3β) and Casein kinase 1 (CK1), resulting in Ror2 and Dvl phosphorylation – commonly used biochemical readouts of Wnt5a-dependent Ror2 signaling^7, 10, 13, 14, 18, 19, 20^.

Transduction of noncanonical Wnt signaling through Wnt5a-induced Ror2 phosphorylation activates downstream effectors including Protein Kinase C (PKC), Jun N-terminal Kinase (JNK), and small GTPases (i.e. Rac1, RhoA), thereby regulating e.g. cytoskeletal remodeling, cell polarity, differentiation, and migration^7, 14, 21^. Importantly, the intensity and duration of Wnt signaling activity are further controlled by endocytic trafficking and lysosomal sorting of Wnt5a-Ror2 complexes, although the mechanisms governing their internalization and subcellular distribution remain poorly understood^4, 7, 9, 14^.

Depending on ligand availability, receptor composition, and cellular context, Ror2 signaling can exert diverse biological outcomes, ranging from pro- and anti-tumorigenic functions to inhibition or potentiation of canonical Wnt/β-catenin signaling^7, 14, 21, 22, 23, 24, 25, 26^. Consequently, perturbation of Ror2 signaling can influence both the canonical and noncanonical Wnt pathways, likely through crosstalk within shared receptor complexes, including co-assembly of Ror2 with Lrp6^7, 14, 21, 27, 28^. In specific biological contexts, Wnt1 or Wnt3a-induced Ror2 activation enhances Lrp6 phosphorylation, β-catenin stabilization, and canonical Wnt target gene expression^23, 24, 25, 29^, illustrating that canonical and noncanonical Wnt signaling represent integrated rather than mutually exclusive signaling pathways. Despite its central role in Wnt signaling, the mechanisms regulating Ror2 activity, trafficking, and receptor homeostasis remain poorly understood.

Fine-tuned Wnt signaling is particularly important for development and function of dopaminergic neurons located in the ventral midbrain (VM)^30, 31, 32, 33^. The murine midbrain dopaminergic (mDA) lineage arises from a pool of proliferating neuroepithelial cells [Sox2-positive (+)] located in the ventricular zone of the VM floorplate. Between embryonic days E9 – E10.5, these primary progenitors specify into mDA progenitors [Lmx1a+, Sox2+], some committing to the multipotent radial glia fate [Lmx1a+, Sox2+, Glast+] between E10.5 - E11.5. The mDA progenitors undergo immense proliferation followed by neurogenesis that generates postmitotic mDA neuroblasts [Lmx1a+, Nurr1+]. Neurogenesis peaks around E11.5 to E12.5 and is finalized by E15. While differentiating, mDA neuroblasts migrate radially along the radial glia projections through the intermediate zone into the marginal zone of the VM floorplate where they become immature mDA neurons [Lmx1a+, Nurr1+, Tyrosine hydroxylase/TH+]. Thereafter, mDA neurons migrate tangentially to form the future mDA nuclei of substania nigra pars compacta and ventral tegmental area. By E12.5 to E13.5, the first mDA neurons are fully differentiated and begin their functional maturation and integration into the brain circuitry^32, 33^.

During mDA development, Wnt1 and Wnt5a are secreted from the VM floorplate and midbrain-hindbrain boundary, two major signaling centers that establish morphogen gradients essential for neural patterning^32, 33, 34, 35^. Wnt1 primarily regulates the early specification and proliferation of mDA progenitors, whereas Wnt5a promotes elongation of mDA domain, neuronal maturation, and innervation. Importantly, synergistic activity of both ligands is required for VM morphogenesis, mDA neurogenesis, and neuronal differentiation. Consistent with this, combined deletion of Wnt1 and Wnt5a results in substantially more severe developmental defects than loss of either ligand alone, demonstrating that both signaling cascades cooperate in spatiotemporal manner to drive normal mDA development^30, 32, 33, 36, 37, 38, 39^.

Despite the established roles of Wnt1 and Wnt5a ligands, the receptor signaling involved in mDA development remains understudied. To narrow this gap, we performed an unbiased immunoprecipitation-coupled tandem mass spectrometry (IP-MS/MS) analysis of Ror2 binding partners in an mDA cell line. We discovered that Ror2 interacts with the Sortilin-related VPS10p Domain-containing receptor 2 (SorCS2) in mDA progenitors, a finding validated *in vitro* and *in vivo*. SorCS2 is a single-pass transmembrane receptor involved in neuronal survival, dopaminergic innervation, and synaptic plasticity^40, 41, 42^. It mostly localizes to intracellular vesicles where it facilitates protein sorting. However, SorCS2 also participates in signal transduction at the plasma membrane by binding various ligands and co-receptors to regulate neurotrophic signaling and Rac1-dependent growth cone retraction^40, 41, 42, 43, 44, 45^. Notably, mice harboring a disruptive insertion in *Sorcs2* exhibit shorter and disorganized cochlear stereocilia^46^, a hallmark of noncanonical Wnt signaling dysfunction, suggesting SorCS2’s association with the Wnt cascades. Recently, *SORCS2* has emerged as a risk gene with common and rare damaging variants to attention-deficit hyperactivity disorder (ADHD), schizophrenia, and bipolar disorder, neuropsychiatric conditions associated with dysfunctional mDA circuits^47, 48, 49, 50, 51, 52^.

This study demonstrates that SorCS2 is essential for regulating Ror2 titration, transport, Wnt5a-dependent activation, and subsequent internalization. SorCS2 loss enhances both Wnt1- and Wnt5a-dependent signaling, resulting in increased proliferation and accelerated neurogenesis of mDA progenitors. Importantly, SorCS2-dependent regulation of Wnt signaling extends beyond the mDA lineage to control anterior-posterior body axis formation, convergent extension, and brain morphogenesis. Together, our findings identify SorCS2 as a conserved, sorting-based regulator of Wnt signaling homeostasis across vertebrates, providing new insight into the spatiotemporal control of Wnt signaling during development and disease.

## Results

### Unbiased identification and validation of Ror2 binding partners in mDA progenitors

To explore the Ror2 interactome, we conducted immunoprecipitation (IP) for Ror2 using the mouse mDA progenitor like cell line SN4741^53^ (**Suppl. Fig. S1a**) and an anti-Ror2 antibody recognizing the C-terminus (Ror2_Sigma) (**Fig. 1f**), followed by tandem mass spectrometry (MS/MS). The pulldown quality was evaluated by immunoblotting and silver staining (**Fig. 1a-b**). We performed four independent experiments, identifying 239-519 specific hits per experiment after subtracting IgG backgrounds (**Fig. 1c, Suppl. Table S1-2**). In two experiments, cells were treated with recombinant Wnt5a for two hours to assess Wnt5a-responsive proteins. This strategy yielded 625 candidate hits detected in two or more replicates (**Fig. 1c, Suppl. Table S3**), and included established Ror2 binding partners like Ror1, Ptk7, CK1, PKC, Calcium/calmodulin-dependent protein kinase II (CAMKII), E3 ubiquitin-protein ligase NEDD4, Filamin-A or 14-3-3, endorsing the data reproducibility (**Suppl. Table S4**)^7, 14^.

**Figure 1:**
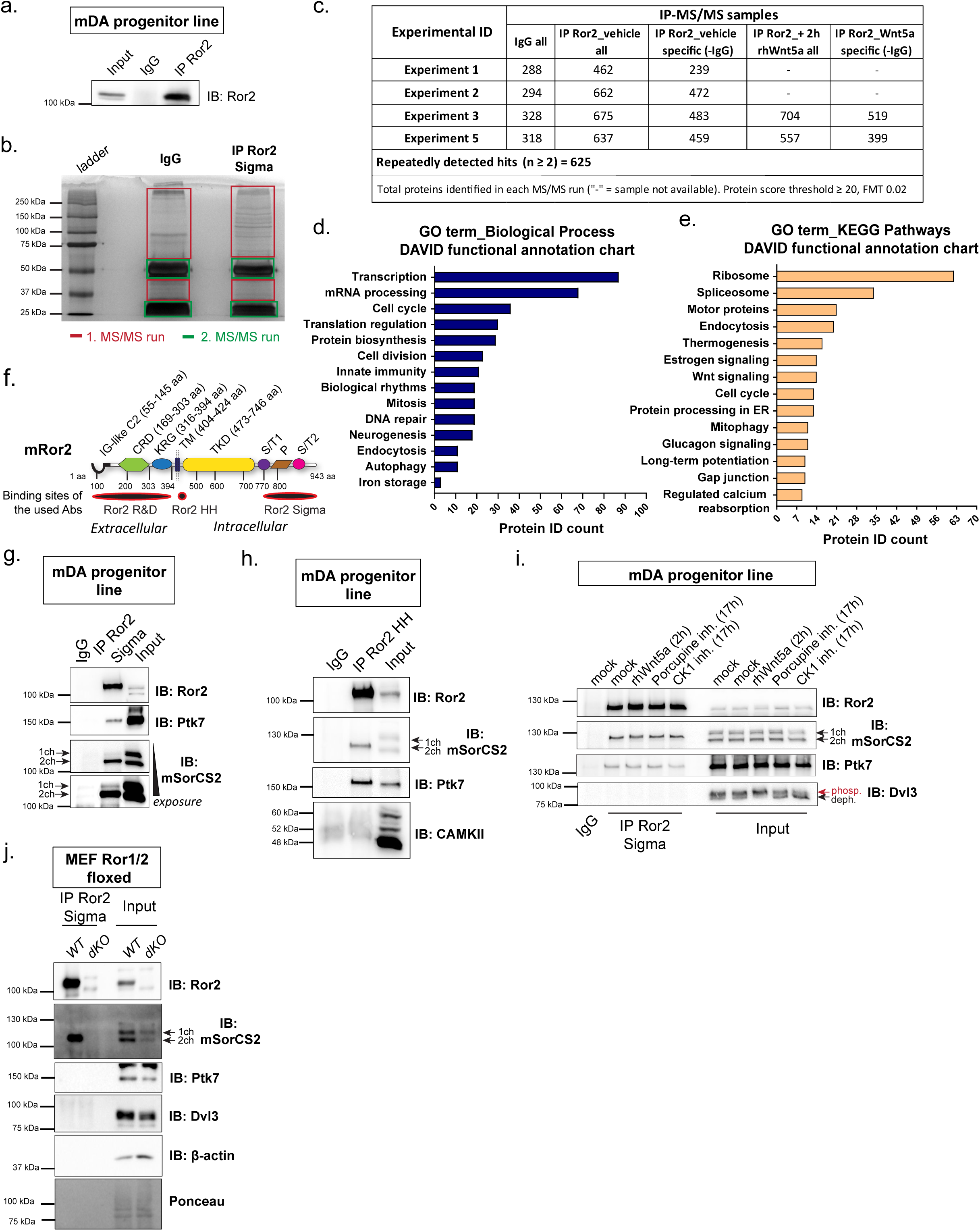
Unbiased identification and validation of Ror2 binding partners in mDA progenitors. **(a)** Immunoblotting (IB) validation of Ror2 immunoprecipitation (IP) from mDA progenitor cell line SN4741 and anti-Ror2_Sigma antibody. IgG served as a negative control. **(b)** Representative silver-stained gel of IgG and enriched IP_Ror2 samples subjected to the IP-MS/MS in two MS/MS runs based on in-gel protein saturation. **(c)** Overview of proteins identified in each IP-MS/MS experiment. Specific Ror2 binding partners were determined by comparison to IgG controls (“-IgG”), with fragment mass tolerance (FMT) of 0.02 and protein score ≥ 20 as thresholds. A total of 625 recurrent Ror2 binding partners were identified and analyzed via DAVID functional annotation for gene ontology (GO) “Biological Process” **(d)** and “KEGG pathway analysis” **(e)**, with 58.7% of proteins annotated in DAVID. Only GO terms with P value < 0.05 are shown. **(f)** Schematic of mouse Ror2 protein structure, highlighting binding sites of two anti-Ror2 antibodies (Abs) used for IP. **(g)** IPRor2**_**Sigma followed by IB validated two Ror2 binding partners, Ptk7 and SorCS2. SorCS2 two-chain (2ch) isoform showed stronger binding to Ror2 than the single-chain (1ch). **(h)** IPRor2**_**HH confirmed the interaction of Ror2 with Ptk7 and SorCS2 two-chain. **(i)** IPRor2_Sigma followed by IB showed that Ror2 interaction with Ptk7 and SorCS2 is unaffected by Wnt5a stimulation or Wnt signaling inhibition (Porcupine inhibitor and casein kinase 1δ/ε inhibitor). Phosphorylated (phosp.) and dephosphorylated (deph.) Dvl3 forms indicate Wnt signaling activation and inhibition, respectively. **(j)** IPRor2_Sigma using Ror1/2-floxed mouse embryonic fibroblasts (MEFs), with Ror1/2 double knockout (dKO) cells serving as a negative control. Ror2 interacts with SorCS2 two-chain but not with Ptk7. All experiments were performed with independent biological and technical replicates, n ≥ 3. Source data are provided as a Source Data file.

Gene ontology (GO) analysis revealed that 77% of the 625 hits comprise phosphoproteins, with many involved in cell cycle, mitosis, neurogenesis, Wnt signaling, endocytosis, and autophagy (**Fig. 1d-e, Suppl. Fig. S1e, Suppl. Tables S5-S6**). KEGG pathway analysis further emphasized associations with neurodegenerative diseases, addiction, and cancer (**Suppl**. **Fig. S1f**). **Suppl. Table S4** highlights several novel candidates, especially those associated with Wnt signaling, receptor biology, cell cycle, and protein trafficking. SorCS2, a transmembrane receptor consistently detected across all replicates, was identified as a top candidate since it is co-expressed with *Ror2, Wnt1,* and *Wnt5a* in the VM floorplate during early embryogenesis^54, 55^ (**Suppl. Table S4, Suppl. Fig. S2a-d**).

Using two independent anti-Ror2 antibodies for pulldown (**Fig. 1f**), we confirmed that Ror2 binds SorCS2 and Ptk7 in mDA progenitor line (**Fig. 1g-i, Suppl. Fig. S1b**). Only the Sigma antibody pulled-down CAMKII, while Dvl3 did not bind Ror2 (**Fig. 1g-i, Suppl. Fig. 1b**), confirming our MS/MS data (**Suppl. Table S4**). Additionally, a pulldown assay performed with *WT* and *Ror1^-/-^; Ror2^-/-^* mouse embryonic fibroblasts (MEF) lines^10^ demonstrated that Ror2 specifically interacts with SorCS2 but not with Ptk7 nor Dvl3 in this cell line (**Fig. 1j**).

SorCS2 exists in two cleavage-specific variants called single-chain (1ch, 122 kDa) and two-chain (2ch, 104 kDa)^42^. The single-chain isoform remains as a full-length protein after synthesis and subsequent propeptide removal. The two-chain isoform arises from additional proteolytic cleavage of SorCS2 single-chain during anterograde trafficking or at the cell surface. This cleavage generates two non-covalently associated fragments (**Suppl. Fig. S1c-d**)^42, 56^. The relative expression, signaling and sorting properties of the single-chain and two-chain isoforms differ between cell types^42^. Notably, endogenous Ror2 preferentially interacted with the SorCS2 two-chain receptor in both cell types (**Fig. 1g-j**).

To evaluate the modulation of Wnt activity, we treated mDA progenitors with recombinant Wnt5a for 2 hours, with porcupine inhibitor (to block Wnt signaling) or with CK1δ/ε inhibitor for 17 hours. Ror2 interactions with SorCS2 and Ptk7 remained consistent across conditions (**Fig. 1i**), indicating that these receptor complexes form independently of Wnt activation status.

### SorCS2 binds Ror2 in *cis* and reduces its protein levels upon overexpression

To characterize the interaction between Ror2 and SorCS2, we overexpressed SorCS2^42^ and Ror2^57^ separately or together in human embryonic kidney 293T cell line (HEK293T). HEK293T cells do not express endogenous SorCS2^58^. As described previously, when expressed alone, full-length Ror2 (Ror2-Flag) localized to the plasma membrane and induced filopodia formation^59^, while SorCS2 full-length (FL) localized to cytosolic vesicles the plasma membrane^40, 42^ (**Fig. 2a, Suppl. Fig. S3a-b**). However, when co-expressed, Ror2 and SorCS2 changed their localization pattern and co-localized at the plasma membrane, in cytosolic vesicles, and in aggregate-like structures (**Fig. 2a, Suppl. Fig. S3a-b**). Co-immunoprecipitation (co-IP; **Fig. 2b**) and proximity ligation assay (PLA; **Fig. 2d**) demonstrated that Ror2 and SorCS2 interact only when co-expressed in the same cell (*cis*-interaction), and not in co-cultures of cells expressing each receptor separately (*trans*-interaction). Contrasting the endogenous data from mDA progenitors (**Fig. 1g-i**), overexpressed Ror2 interacted with SorCS2 single-chain (**Fig. 2b**). This discrepancy may be attributed to differences in cell type, protein levels, and/or the ratio between single and two-chain variants.

**Figure 2:**
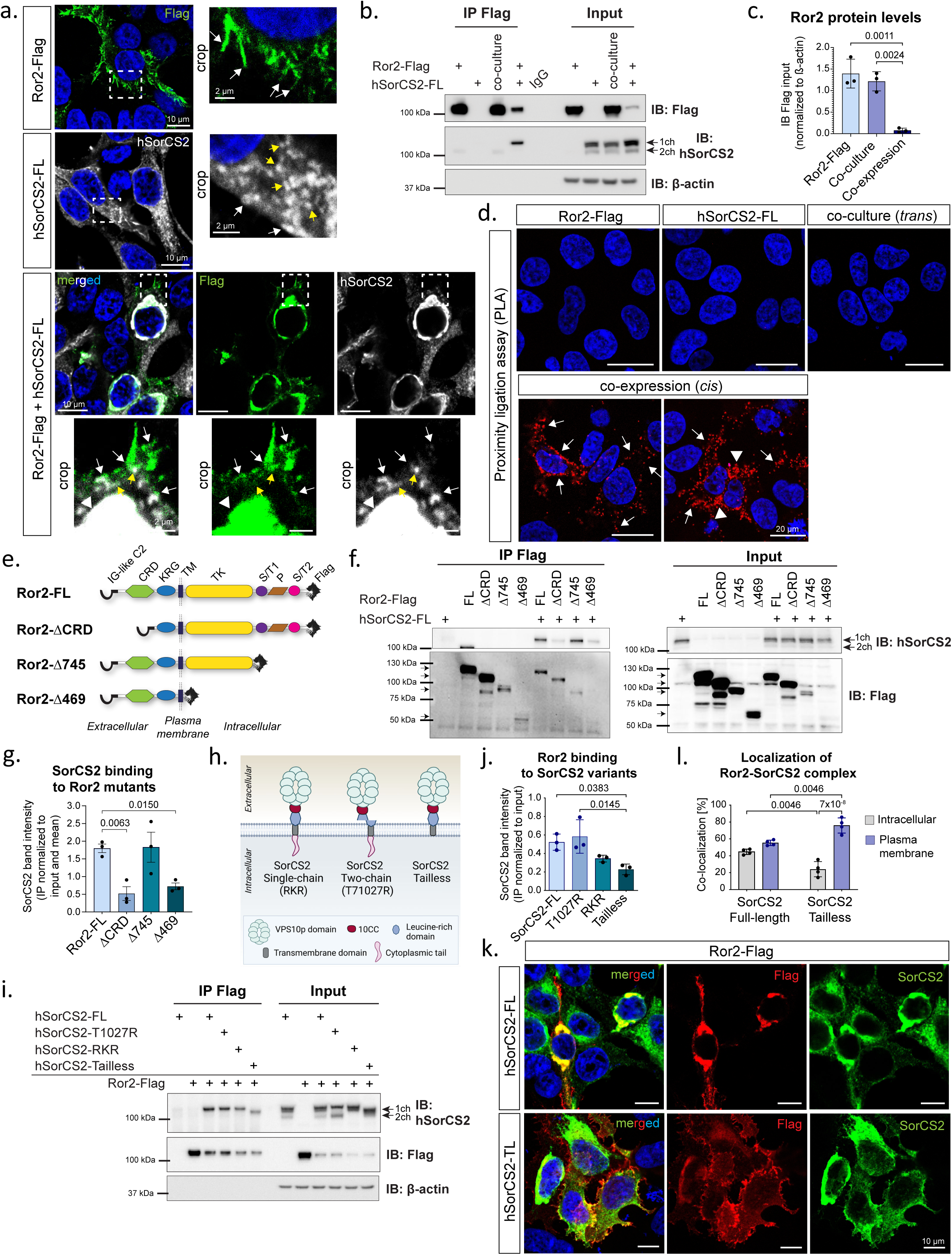
Domain mapping of the Ror2-SorCS2 interaction. Overexpression studies using HEK293T cells. **(a)** *Upper panels:* Immunofluorescence followed by confocal scanning. Ror2 full-length (Ror2-Flag; green) localized mostly to the plasma membrane where it induced filopodia (white arrows), while SorCS2 full-length (SorCS2-FL; grey) was primarily vesicular (yellow arrows) or plasma membrane-associated (white arrows). *Lower panels*: Co-expression of the constructs led to reduced filopodia formation (also **Suppl. Fig. S3a**). Ror2-SorCS2 co-localized at the membrane (white arrows), intracellular sorting vesicles (yellow arrows) and aggregate-like structures (arrowheads). Scale bar = 10 µm/2 µm in crop. Hoechst in blue. **(b)** Co-immunoprecipitation (IP) of Ror2 shows binding to SorCS2 single-chain only when co-expressed (*cis-*interaction) but not when co-cultured (*trans-*interaction). SorCS2 overexpression downregulates Ror2 protein levels, quantified in **(c)**, one-way ANOVA (P = 0.0009) with Tukey’s multiple comparison test. **(d)** Proximity ligation assay demonstrates a specific *cis*-interaction of Ror2 with SorCS2 at the plasma membrane (arrows), and in intracellular vesicles and aggregates (arrowheads). Scale bar = 20 µm. Hoechst in blue. **(e)** Schematic of C-terminally Flag-tagged Ror2 constructs. **(f)** Co-IP of Ror2 constructs with SorCS2-FL reveals that Ror2-SorCS2 interaction is dependent on the CRD domain, quantified in **(g)** using one-way ANOVA (P = 0.0088) and Fisher’s LSD test. **(h)** Schematic of SorCS2 structure and constructs used in the study. Created with Biorender. **(i)** Ror2 pulldown with SorCS2 constructs. SorCS2 variants show differential binding to Ror2, data quantified in **(j)** using one-way ANOVA (P = 0.0122) and Tukey’s multiple comparison test. **(k)** Immunofluorescence of cells co-expressing Ror2-Flag (red) with SorCS2-FL or SorCS2-Tailless (green). While SorCS2-FL co-localized with Ror2 at the intracellular structures and plasma membrane comparably, the deletion of SorCS2 tail retained the receptors mostly at the plasma membrane. Scale bar = 10 µm. Hoechst in blue. These data are quantified in **(l)** using one-way ANOVA (P = 0.000001) with Tukey’s multiple comparison test. Graphs represent mean ± S.E.M, with individual data points representing independent biological experiments; n = 3 **(b-c, f-g, i-j)** and n =4 **(k-l)**. Source data are provided as a Source Data file.

Notably, SorCS2 overexpression led to significant Ror2 downregulation (**Fig. 2b-c**) and reduced filopodia formation (**Fig. 2a, Suppl. Fig. S3a-b**), suggesting that SorCS2 might regulate Ror2 protein levels. Control experiments confirmed the selectivity of this phenotype, as Ror2 downregulation was not observed when co-expressed with Lrp4 or the SorCS2’s paralog SorCS1 (**Suppl. Fig. S3c-d**). The specificity of the Ror2-SorCS2 complex was further confirmed by immunofluorescence, which showed minimal to no co-localization between co-expressed Ror2 and SorCS1, or SorCS2 and Dvl2 (**Suppl. Fig. S3e-f**). These results highlight a selective, *cis*-specific regulation of Ror2 by SorCS2 when overexpressed.

In line with this, SorCS2 overexpression also reduced the levels of key Ror2-associated binding partners, i.e. Ror1, Ptk7, Vangl2, and Wnt5a (**Suppl. Fig. S4a-d**). Co-immunoprecipitation further revealed a physical interaction between SorCS2 and intracellular Wnt5a (**Suppl. Fig. S4e**). Together, these data indicate that SorCS2 might associate with multiple noncanonical Wnt components upon co-expression, including Wnt ligands.

### Domain mapping of the Ror2-SorCS2 interaction

To identify the Ror2 domain responsible for SorCS2 binding, we co-expressed SorCS2-FL with Ror2-FL and three truncated mutants: Ror2-ΔCRD (extracellular CRD domain deletion), Ror2-Δ745 (Serine/Threonine and Proline-rich domains deletion), or Ror2-Δ469 (entire intracellular terminus deletion) (**Fig. 2e**). Co-IP and co-localization analyses demonstrated that Ror2-SorCS2 interaction is reliant on the CRD domain (**Fig. 2f-g, Suppl. Fig. S5a-i**). The apparent reduction in binding of the Ror2-Δ469 mutant should be interpreted cautiously due to its low protein expression levels. Despite clear co-localization with SorCS2 (**Suppl. Fig. S5i**), this construct was excluded from quantitative analysis to avoid bias arising from unequal protein abundance (**Fig. 2g**).

Next, we examined whether SorCS2 variants differ in the interaction with Ror2. Full-length Ror2 was co-expressed with SorCS2-FL, truncated SorCS2-Tailless (cytoplasmic tail deletion), and two mutants with disrupted cleavage sites representing isoform imbalances: SorCS2-RKR (producing only the single-chain) and SorCS2-T1027S (favoring the production of the two-chain)^42^ (**Fig. 2h-j**). Co-IP showed that Ror2 interacted with the SorCS2 single-chain under all conditions. The interaction was comparable when co-expressed with SorCS2-FL and SorCS2-T1027S, whereas SorCS2-RKR and SorCS2-Tailless variants showed reduced binding to Ror2 by 30% and 50%, respectively (**Fig. 2i-j**). Given that no significant difference was observed between single- and two-chain SorCS2, the reduced co-precipitation of the tailless construct likely reflects altered subcellular distribution rather than impaired binding per se.

To further assess the role of cytoplasmic tail, we examined how its deletion affects Ror2-SorCS2 subcellular localization. In cells expressing SorCS2-FL, approximately 58% of the Ror2-SorCS2 complex localized to the plasma membrane, with the remaining 42% found intracellularly. Conversely, SorCS2-Tailless increased the plasma membrane localization of Ror2 to about 79%, with 21% being in intracellular aggregates (**Fig. 2k-l**). These findings indicate that the cytoplasmic tail of SorCS2 plays a crucial role in regulating Ror2 trafficking and subcellular distribution.

### SorCS2 regulates Ror2 trafficking through endosomal sorting and ER-associated lysosomal degradation of excess Ror2

Next, we investigated the SorCS2-dependent trafficking and degradation of Ror2. IP-MS/MS analysis of Ror2 binding partners revealed proteins involved in endocytosis and autophagy (**Fig. 1d-e**), prompting us to examine associated pathways. SorCS2-FL and Ror2-FL constructs were overexpressed individually or co-expressed in HEK293T cells, which was followed by a 24-hour treatment with the lysosome inhibitor chloroquine and the dynamin-dependent endocytosis inhibitor dynasore (**Fig. 3a-c, Suppl. Fig. S6a**). Both treatments significantly increased Ror2 levels in the co-expression compared to mock and single Ror2 expression (**Fig. 3a-c**). These data suggest that SorCS2 traffics Ror2 via endosomal and lysosomal pathways. To assess the involvement of proteasome-mediated degradation, we performed a ubiquitination assay with additional co-expression of His-tagged ubiquitin, followed by His pulldown and immunoblotting^60^. SorCS2 expression did not increase the protein smear in the anti-Flag immunoblot, indicating that Ror2 is not targeted for proteasome-mediated degradation (**Suppl. Fig. S6b**).

**Figure 3:**
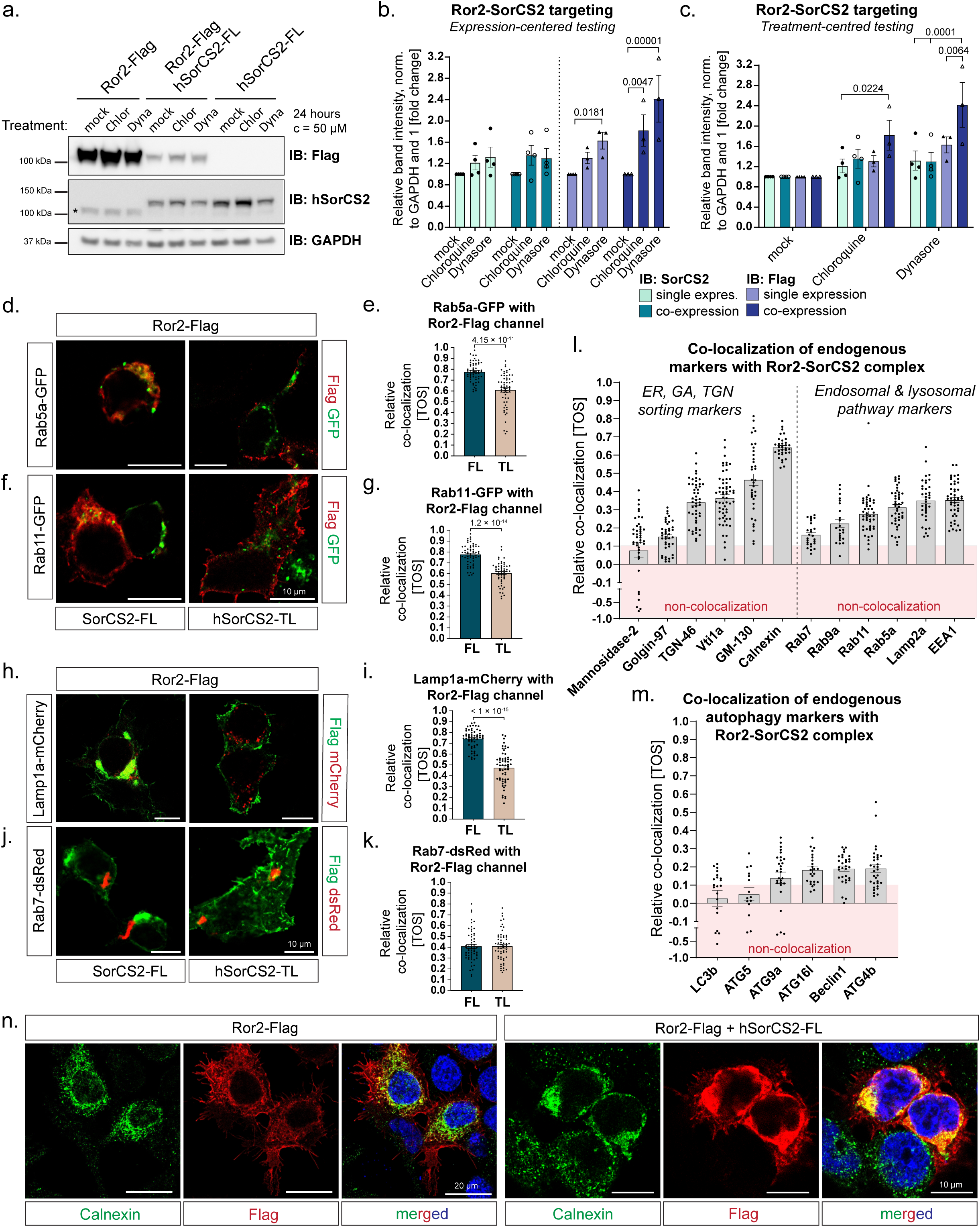
SorCS2 regulates Ror2 trafficking and mediates ER-to-lysosome associated degradation of excess Ror2. **(a)** Immunoblotting of HEK293T cells co-expressing Ror2-Flag and SorCS2-FL were treated with the lysosome inhibitor chloroquine or the dynamin-dependent endocytosis inhibitor dynasore for 24 hours. Star = unspecific band. **(b-c)** Quantification of the immunoblots from four independent biological experiments, two-way ANOVA (P_Treatment_ = 0.00003, P_Expression_ = 0.0022) with Fisher’s LSD test. Relative band intensity was first normalized to the loading control GAPDH then to 1 for mock, displaying fold change. Both treatments, particularly dynasore, significantly rescued Ror2 levels in co-expression compared to mock and single expression conditions. Multiple-comparisons analysis was performed with comparisons either centered on the expression profile **(b)** or treatment **(c)**. **(d-j)** Immunofluorescence (IF), confocal scanning and co-localization analysis of Ror2-Flag, SorCS2-FL, or SorCS2-Tailless in HEK293T cells co-expressed with constructs encoding fluorescently labelled sorting proteins: Rab5a-GFP **(c-d)**, Rab11-GFP **(e-f)**, Lamp1a-mCherry **(g-h)**, and Rab7-dsRed **(i-j)**. Statistical significance was assessed using a two-tailed Mann–Whitney test; n = 60 cells per group from three independent biological experiments. **(c-h)** Rab5a, Rab11, and Lamp1a co-localized with the Ror2-SorCS2 complex, which was dependent on SorCS2’s tail. **(i-j)** Rab7 exhibited low co-localization with the Ror2-complex and it was not affected by the SorCS2 tail deletion. Representative IF images are shown in **(Suppl. Fig. S5b-f)**. Scale bar = 10 µm. Hoechst in blue. **(k-l)** Co-localization analysis of the Ror2-SorCS2 complex with endogenous cellular compartment markers in HEK293T cells, based on confocal imaging data **(Suppl. Figs. S6-S7)**. Three independent biological experiments, n = 16-60 cells/group. **(k)** Co-localization analysis with markers for Endoplasmic Reticulum (ER), Golgi Apparatus (GA) and *Trans-*Golgi Network (TGN), together with markers for endosomal and lysosomal pathways. **(l)** Co-localization analysis of endogenous markers for autophagy. Representative IF images are shown in **(Suppl. Figs. S8). (m)** Representative confocal images of the ER marker Calnexin in Ror2-Flag single expression or co-expression with SorCS2-FL using HEK293T cells. Ror2-SorCS2 intracellular staining strongly co-localized with the ER marker. Hoechst in blue, scale bars = 20/10 µm. The graphs represent mean ± S.E.M. Source data are provided as a Source Data file.

To assess the localization of the Ror2-SorCS2 complex within sorting pathways, we next co-expressed the two full-length receptors with constructs encoding fluorescently labelled markers for sorting vesicles. We found that the Ror2-SorCS2 complex co-localized with early endosomes (Rab5a) (**Fig. 3d-e, Suppl. Fig. S6c**), early/recycling endosomes (Rab4) (**Suppl. Fig. S6d**), recycling endosomes (Rab11) (**Fig. 3f-g, Suppl. Fig. S6e**), and lysosomes (Lamp1a) (**Fig. 3h-i, Suppl. Fig. S6g**), but less so in late-endosomes-to-lysosomes (Rab7) (**Fig. 3j-k, Suppl. Fig. S6f**). Notably, except for Rab7, these co-localizations were reduced in the presence of SorCS2-Tailless, demonstrating the requirement of the cytosolic terminus of SorCS2 for its capacity to mediate endosomal and lysosomal targeting of Ror2 (**Fig. 3d-k**).

To further characterize these findings, we employed a panel of antibodies labeling endogenous organelle and sorting proteins. Indeed, the co-expressed Ror2-SorCS2 co-localized with early endosomes (EEA1, Rab5a), recycling endosomes (Rab11), late-endosome-to-TGN vesicles (Rab9a), and lysosomes (Lamp2a), with minimal co-localization in late-endosomes-to-lysosomes (Rab7) (**Fig. 3l, Suppl. Fig. S7**). When evaluating cellular compartments involved in sorting, Ror2-SorCS2 complex co-localized most prominently with the endoplasmic reticulum (Calnexin) (**Fig. 3l, n**), and with proteins involved in ER-Golgi vesicle trafficking (GM130) and trans-Golgi sorting and recycling within secretory and endosomal systems (Vti1a, TGN46) (**Fig. 3l, Suppl. Fig. S8a-c**). Minimal to non-colocalization was detected with *medial*-Golgi apparatus marker (Mannosidase-2), Golgi-specific vesicle tethering marker (Golgin-97) (**Fig. 3l, Suppl. Fig. S8d-e**), and autophagy markers (**Fig. 3m, Suppl. Fig. S9**).

These findings suggest that the SorCS2 has a potential to retain excess Ror2 in the ER, likely targeting it for the ER-mediated lysosomal degradation to prevent pathway hyperactivation^61, 62^. SorCS2 may then facilitate the transport of residual Ror2 to the plasma membrane, ensuring its availability for signaling, and the subsequent internalization and sorting through recycling or endosomal pathways.

### SorCS2 regulates noncanonical Wnt signaling and mitigates hyperactivated Ror2 activity in gain-of-function assays using *Xenopus laevis*

Convergent extension (CE) is a fundamental process in embryogenesis that drives tissue elongation and narrowing. Disruptions in noncanonical Wnt signaling homeostasis during CE - whether due to excessive, deficient, or mislocalized protein activity - result in severe phenotypes, including body axis truncation, aberrant somite formation, and neural tube defects. These phenotypic outcomes serve as key readouts of Ror2 signaling activity *in vivo*, particularly in *Xenopus laevis (X. laevis)* and zebrafish embryos^7, 8, 9, 63^.

To evaluate the functional interaction between Ror2 and SorCS2 *in vivo*, we first conducted gain-of-function assays in *X. laevis*. Marginal-to-dorsal injections of *Ror2* mRNA at the 4-8 cell stage embryos produced dose-dependent CE defects, including body axis truncation and distortion (**Fig. 4a-c**). CE defects were categorized into four classes: normal, mild, moderate, and severe (**Fig. 4b-c**). Injection of *SorCS2* mRNA resulted in mild CE defects and reduced head and eye development (arrows) (**Fig. 4a, c**). Remarkably, co-expression of *SorCS2* with *Ror2* rescued the moderate and severe CE phenotypes induced by excess *Ror2* (**Fig. 4a, c**). Similar results were obtained from animal cap assay, a method that takes advantage of Activin-A treatment that induces CE movements in cultured animal cap explants^64, 65^ (**Fig. 4d**). While the control animal caps elongated by ∼60% compared to untreated tissue, the explants injected with *Ror2* or *SorCS2* failed to elongate, indicating dysregulated noncanonical Wnt signaling. Strikingly, co-injection of *SorCS2* with *Ror2* partially rescued the elongation defects (by ∼30%) caused by excess *Ror2* (**Fig. 4d-e**). These data demonstrate that SorCS2 regulates noncanonical Wnt signaling and mitigates hyperactivated Ror2 signaling *in vivo*.

**Figure 4:**
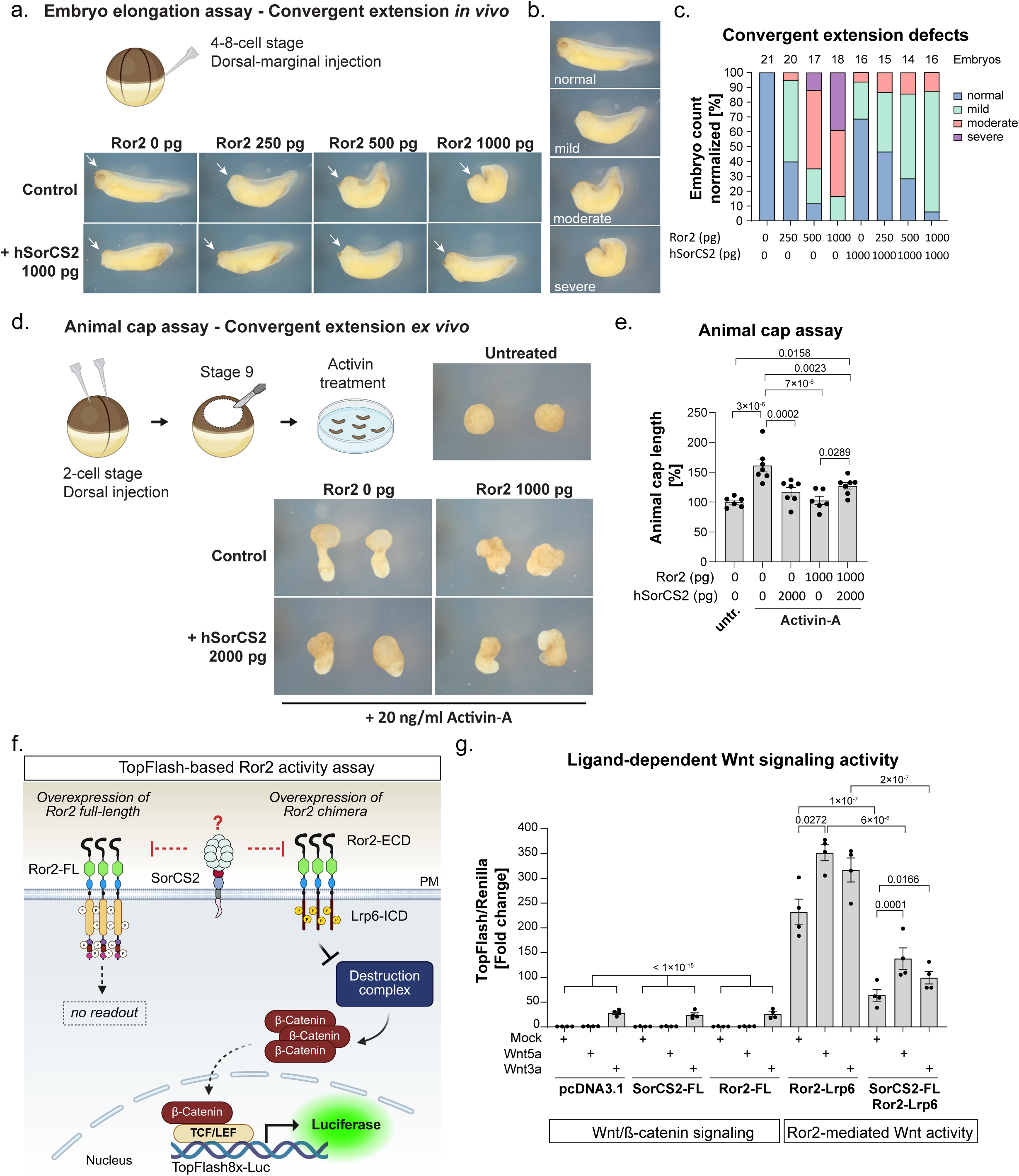
SorCS2 overexpression disrupts noncanonical Wnt signaling and attenuates Ror2 hyperactivation in *Xenopus laevis*. **(a–c)** Embryo elongation assays assessing noncanonical Wnt signaling activity during *X. laevis* convergent extension (CE), using individual embryos collected across three independent biological experiments. **(a)** Ror2 RNA injection into the dorsal-marginal region of 4- to 8-cell stage embryos caused abnormal noncanonical Wnt signaling, resulting in truncated, distorted body axis, and reduced head development (arrows) by stage 32. Higher Ror2 concentrations aggravated the phenotype. Injection of SorCS2 RNA alone induced mild CE defects, including reduced head (arrows). Co-injection of Ror2 with SorCS2 RNA partially rescued the Ror2-induced CE defects. **(b)** Categorized CE defects reflecting the activity of noncanonical Wnt signaling. Quantification of the CE defects in individual embryos using the embryo elongation assay **(c). (d)** Animal cap elongation assay performed on embryos injected with Ror2 and/or SorCS2 RNA into the animal pole at the 2-cell stage. Animal caps were dissected at stage 9, treated with recombinant Activin-A for one hour to induce CE movements, and cultured until sibling embryos reached stage 17. **(e)** Quantification of animal cap elongation of individual embryos shows that while individual injections of Ror2 or SorCS2 caused anterior-posterior elongation defects, co-injection partially rescued the phenotype. One-way ANOVA (P = 0.000015) with Fisher’s LSD test. **(f)** Schematic of the TopFlash-based Ror2 activity assay. A chimeric Ror2-Lrp6 receptor [Ror2 extracellular domain (ECD) fused to the Lrp6 intracellular domain (ICD)] was used to convert Ror2 activation into Lrp6-ICD phosphorylation, promoting nuclear β-catenin accumulation and the activation of the TopFlash reporter (8×TCF/LEF sites upstream of luciferase). PM = plasma membrane. **(g)** Dual-luciferase assay in HEK293T cells co-transfected with SuperTopFlash8X, Renilla (internal control of translation), Ror2-Flag (FL), SorCS2-FL, and Ror2-Lrp6 constructs. The cells were treated with PBS-0.1%BSA (mock), and recombinant Wnt3a and Wnt5a (100 ng/ml) for 16h. Firefly luciferase activity was normalized to Renilla and to 1. pcDNA3.1 served as basal Wnt activity control. Four independent biological experiments, One-way ANOVA (P < 1×10⁻¹⁵) with Fisher’s LSD test. Graphs represent mean ± S.E.M. Source data are provided as a Source Data file. Schematics created with BioRender.

Due to the lack of direct *in vitro* readouts for Ror2 activity, we investigated the impact of SorCS2 on Ror2 hyperactivation using a chimeric Ror2(ECD)-Lrp6(ICD) reporter, in which the extracellular domain of Ror2 is fused to the intracellular domain of Lrp6, thereby converting Ror2 activation into a measurable TopFlash luciferase readout (**Fig. 4f**)^28, 66^. A dual-luciferase reporter assay was performed in HEK293T cells co-expressing SuperTopFlash8x-Luc^63^, Renilla-tkLuc (internal normalization control), full-length Ror2-Flag, SorCS2-FL, and the Ror2-Lrp6 construct, in the absence or presence of Wnt5a or Wnt3a stimulation. Both Wnt5a and Wnt3a ligands are known to increase Ror2-Lrp6 reporter activity, although only Wnt5a induces Ror2 phosphorylation^28^. Consistent with previous reports^28, 66^, Ror2-Lrp6 overexpression markedly hyperactivated Ror2-mediated Wnt activity, which was further potentiated by Wnt5a and Wnt3a treatment (**Fig. 4g**). Notably, co-expression of SorCS2-FL significantly reduced Ror2 hyperactivation under basal conditions as well as upon Wnt5a and Wnt3a stimulation. Importantly, despite reducing Ror2 hyperactivation, SorCS2 preserved responsiveness to Wnt ligands, indicating that SorCS2 limits excessive Ror2 signaling without abolishing Wnt signal transduction (**Fig. 4g**).

Together, these findings demonstrate that SorCS2 is part of a regulatory system that limits excessive Ror2 signaling to maintain Wnt signaling homeostasis. Furthermore, because elevated SorCS2 levels alone can disrupt noncanonical Wnt signaling, our findings underscore the importance of balanced SorCS2 expression during embryogenesis.

### Endogenous Ror2-SorCS2 signaling controls convergent extension in zebrafish

To assess SorCS2 regulation of Ror2-dependent noncanonical Wnt signaling across species, we used *Danio rerio* (zebrafish) as an *in vivo* model^8^. Unlike *X. laevis*, CE defects in zebrafish have not been consistently observed upon Ror2 overexpression. In line with this, overexpression of mouse Ror2 mRNA did not induce CE defects (**Suppl. Fig. S10a**), demonstrating that Ror2 overexpression is not a suitable gain-of-function readout of noncanonical Wnt signaling activity in zebrafish. Instead, CE defects emerge only after *ror2* knockdown or in *ror1;ror2* double-knockout genetic lines^67, 68, 69, 70^, the latter precluding ror2-specific analysis. We therefore focused on targeted endogenous loss-of-function approaches during the first 24 hours post-fertilization (hpf) (**Fig. 5a**). Besides *ror2* and *sorcs2*, we also examined *wnt5b*, a fish orthologue of mouse/human Wnt5a.

**Figure 5:**
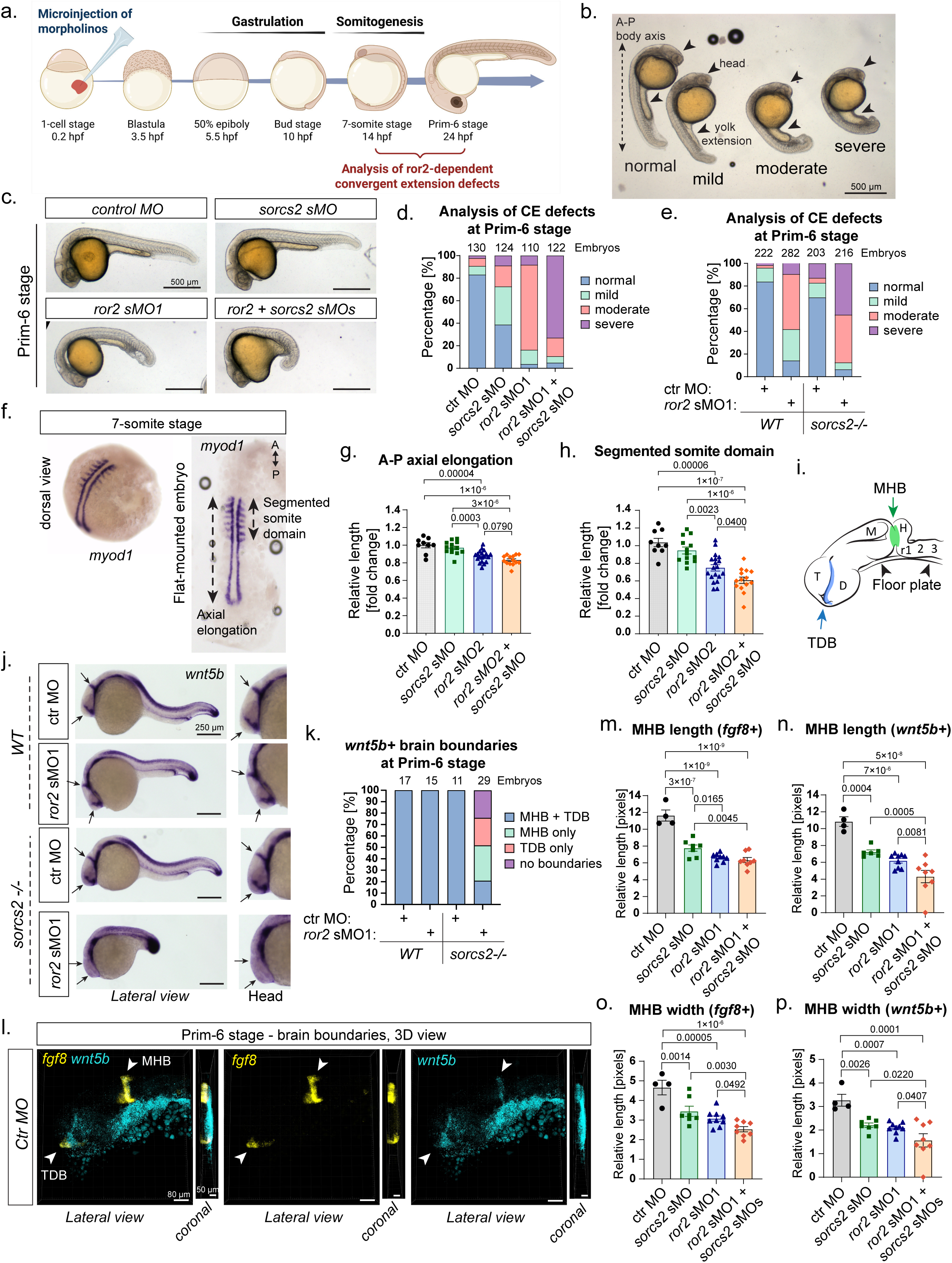
The Ror2-SorCS2 complex regulates convergent extension and brain morphogenesis in zebrafish. **(a)** Strategy to assess endogenous noncanonical Wnt signaling. Hpf = hours post fertilization. Created with BioRender. **(b)** Classification of CE defects: anterior–posterior (A-P) axis truncation, shortened/widened yolk extension, and reduced head size. *ror2* splice morpholinos (sMOs) were titrated to induce a moderate phenotype. **(c-d)** Embryos injected with control morpholino (ctr MO), *sorcs2*-specific splice-site morpholino (*sorcs2* sMO), *ror2* sMO1, and a combination of *sorcs2* and *ror2* sMOs. Co-injection of *ror2* with *sorcs2* sMOs exacerbated *ror2*-dependent body axis and head defects. **(e)** Quantification of *ror2* sMO1 injection in *WT* and *sorcs2^−/−^* embryos showing aggravated CE defects in mutants. Three **(d)/**five **(e)** independent microinjections. **(f-h)** Alkaline phosphatase WISH labelling *myod1* transcripts at the 7-somite stage for quantification of A-P axis **(g)** and segmented somite length **(h)**. Three independent experiments (embryos n = 9 *ctr*, n = 13 *sorcs2* MO, n = 20 *ror2* MO, and n = 14 *sorcs2+ror2* MOs). One-way ANOVA with Fisher’s LSD test [P = 1.8×10⁻⁷, **(g)**, P = 4.6×10⁻⁸ **(h)**]. **(i)** Schematic of Prim-6 brain anatomy. T = telencephalon, D = diencephalon, M = midbrain, H = hindbrain, segmented into rhombomeres (r1-r7). TDB = telencephalon-diencephalon boundary (blue), MHB = midbrain-hindbrain boundary (green). **(j-k)** Alkaline-phosphatase WISH for *wnt5b* in *WT* and *sorcs2^−/−^* embryos injected with *contro*l MO or *ror2* sMO1. *ror2* knockdown in *sorcs2^−/−^*embryos led to loss of *wnt5b* expression at one or both brain boundaries (arrows). Embryos quantified from two independent experiments. **(l)** HCRv3.0 detecting *fgf8* and *wnt5b* transcripts for 3D analysis of MHB morphology. Prim-6 confocal Z-stacks. Scale bars: 80 µm (lateral views), 50 µm (coronal views). **(m-p)** Quantification of MHB morphogenesis based on *fgf8* **(m, o)** and *wnt5b* **(n, p)** domains (embryos n = 4 *ctr*, n = 7 *sorcs2* MO, n = 9 *ror2* MO, and n = 8 *sorcs2+ror2* MOs). One-way ANOVA [**(m)** P = 3.1×10⁻⁹, **(n)** P = 6.38×10⁻⁷, **(o)** P = 1.28×10⁻⁵, **(p)** P = 0.00016] with Fisher’s LSD test. Graphs represent mean ± S.E.M. Source data are provided as a Source Data file.

Using whole-mount RNA *in situ* hybridization methods (alkaline phosphatase and fluorescent multi-probe HCR3.0^71^), we detected *sorcs2* transcripts from the blastula stage onward (**Suppl. Fig. S11a-e, Suppl. Fig. S12a-c**). *sorcs2* was co-expressed with *ror2* at all analyzed stages, particularly in the developing brain (**Suppl. Fig. S11b-e, S12a-c**). *ror2, sorcs2,* and *wnt5b* were co-expressed in the forebrain and midbrain around the telencephalon-diencephalon (TDB) and midbrain-hindbrain boundaries (MHB), structures critical for neural patterning (**Suppl. Fig. S12a-d**). Consistent with the zebrafish data, *Sorcs2* and *Wnt5a* were co-expressed at the MHB in mouse embryos (**Suppl. Fig. S2b-c).**

The expression profiles suggested that SorCS2 may functionally interact with Ror2 during CE and brain morphogenesis. To test this hypothesis, we performed knockdowns (KD) using validated morpholino oligonucleotides (MO) targeting splice-sites of *sorcs2* (*sorcs2* sMO)^58^ and *ror2* (*ror2* sMO1 and sMO2)^67^ at 1-4-cell stage (**Fig. 5a**). We assessed the established Ror2-dependent CE defects^67^ in live control and *ror2* KD embryos at the Prim-6 stage. These included anterior-posterior body axis truncation (CE), shortened and widened yolk extension (CE), and reduced head development. Phenotypes were classified according to severity as normal, mild, moderate, or severe (**Fig. 5b**). Using two independent *ror2* morpholinos (sMO1, sMO2), we titrated *ror2* KD to achieve a moderate phenotype. *Sorcs2* KD caused a mild CE defects indicating that SorCS2 deficiency may affect noncanonical Wnt signaling. Strikingly, combined *sorcs2* and *ror2* KDs significantly exacerbated the CE defects with either *ror2* sMO, compared to single KD alone and irrespective of genetic background (**Fig. 5c-d, Suppl. Fig. S10b-c**). These data suggested that SorCS2 and Ror2 functionally interact to regulate noncanonical Wnt signaling during CE and brain morphogenesis.

To rule out off-target effects, we performed microinjections in *WT* and *sorcs2^-/-^* mutant embryos^58^. While *sorcs2^-/-^* embryos exhibited mild CE defects, simultaneous *ror2* KD resulted in severe defects compared to *ror2* KD in *WT* embryos (**Fig. 5e, Suppl. Fig. S10d**). Finally, injecting translation-inhibiting *wnt5b* tMO or *sorcs2* sMO into *sorcs2^-/-^* embryos did not aggravate the CE defects (**Suppl. Fig. S10e-f**).

Finally, to assess noncanonical Wnt signaling activity at an earlier stage of convergent extension, we performed *myod1* RNA *in situ* hybridization at the 7-somite stage to visualize and quantify relative anterior-posterior axial elongation and the length of the segmented somite domain (**Fig. 5f**). All embryos were stage-matched based on the somite number. Both *ror2* KD and combined *ror2; sorcs2* KDs resulted in reduced axial elongation (**Fig. 5g**). However, the length of segmented somite domain was significantly reduced in double *ror2; sorcs2* KD embryos compared to *ror2* KD alone (**Fig. 5h**). These findings demonstrate that SorCS2 functionally interacts with Ror2 at endogenous levels, and that disruption of Ror2-SorCS2 signaling impairs noncanonical Wnt activity during convergent extension and somitogenesis.

### Ror2-SorCS2 signaling controls brain morphogenesis in zebrafish

Since Wnt5a is the main ligand for Ror2, we investigated whether Ror2-SorCS2 signaling affects its expression. During somitogenesis, the *Wnt5a* ortholog *wnt5b* was highly expressed in the tail and partially in the somites (**Suppl. Fig. S11c-d**). At the Prim-6 stage, *wnt5b* was highly expressed in the floorplate, telencephalon-diencephalon boundary, and midbrain-hindbrain boundary (**Fig. 5j, Suppl. Fig. S12a-d**). While deficiency of either *sorcs2* or *ror2* alone did not prominently alter *wnt5b* levels, the simultaneous loss of both *ror2* and *sorcs2* drastically reduced *wnt5b* transcripts at the brain boundaries (**Fig. 5j-k, Suppl. Fig. S12e-f**). These results uncover the additional role of the Ror2-SorCS2 signaling in neural patterning.

To determine whether Ror2-SorCS2 loss-of-function affects overall brain morphogenesis or specifically *wnt5b* expression, we used 3D confocal imaging to visualize *wnt5b* transcripts at the midbrain-hindbrain boundary together with its established marker *fgf8*^34, 35^ (**Fig. 5i, Suppl. Fig. S12g**). Lateral and coronal measurements revealed a reduction in midbrain-hindbrain boundary length and width following *sorcs2* KD or *ror2* KD compared to control. Notably, double *ror2; sorcs2* KD further exacerbated these defects (**Fig. 5m-p**). These findings underscore a synergistic role of Ror2-SorCS2 signaling in shaping *wnt5b*-expressing brain boundaries during morphogenesis.

### SorCS2 loss causes transient defects in anterior-posterior axis elongation at E12.5

To assess whether Ror2-SorCS2 signaling contributes to tissue patterning and body plan organization in mice, we analyzed Wnt5a/Ror2-related macro-anatomical morphometric features, such as anterior-posterior body axis formation, craniofacial defects, and limb development^7, 14^. Adult *Sorcs2^-/-^* mice display a truncated body axis with reduced trunk and tail length, as well as decreased body weight (**Suppl. Fig. S13a-c**), each independent of sex. Whereas *Sorcs2^-/-^*mice are viable^42^, *Ror2^-/-^* pups fail to survive postnatally, owing to severe developmental defects^72^. We therefore generated *Ror2^+/-^; Sorcs2^+/-^* mice by crossing *Ror2^+/-^* ^72^ and *Sorcs2^+/-^* ^42^ lines, enabling littermate comparisons at E12.5 and E14.5 (**Suppl. Fig. S13d, S14a**).

At E14.5, *Ror2^+/+^; Sorcs2^-/-^* embryos were indistinguishable from controls, whereas *Ror2^-/-^; Sorcs2^+/+^* and *Ror2^-/-^; Sorcs2^-/-^* embryos exhibited severe truncation of the anterior-posterior (A-P) body axis, including reduced brain proportions (**Suppl. Fig. S13b-e**), craniofacial defects (**Suppl. Fig. S13g-h**), and shortened limbs (**Suppl. Fig. S13i-j**). These features are characteristic of *Ror2* loss in mice and consistent with established impairments of noncanonical Wnt signaling activity^10, 12, 72^. Forebrain-midbrain thickness and eye diameter remained unchanged across genotypes (**Suppl. Fig. S13f, k**).

At E12.5, craniofacial defects were not yet apparent (**Fig. 6a-b, Suppl. Fig. S13d**). However, *Ror2^-/-^; Sorcs2^+/+^* embryos already showed truncation of the A-P axis driven by a shortened trunk (**Fig. 6c-d**), while brain proportions were unchanged (**Fig. 6e-f**). Strikingly, *Ror2^+/+^; Sorcs2^-/-^* embryos exhibited an elongated A-P axis by approximately ∼900 µm relative to *Ror2^+/+^; Sorcs2^+/-^* controls, reflecting increased trunk length (∼500 µm) and enlarged brain dimensions (∼250 - 400 µm) (**Fig. 6c-f**). Forebrain-midbrain thickness was further enhanced in *Ror2^-/-^; Sorcs2^-/-^* embryos, despite trunk truncation comparable to *Ror2^-/-^; Sorcs2^+/+^* embryos (**Fig. 6d-f**).

**Figure 6:**
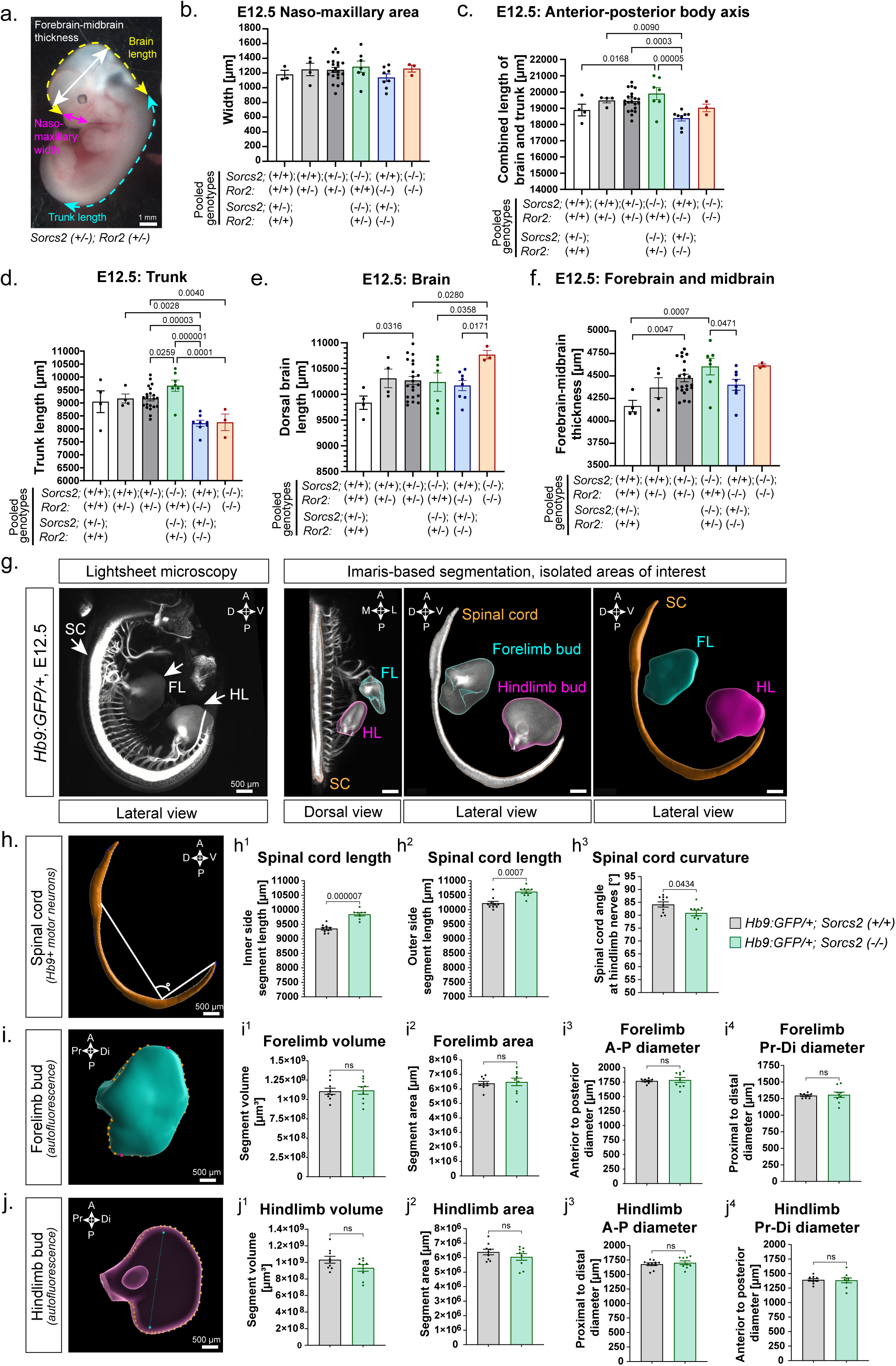
SorCS2 loss causes transient defects in the anterior–posterior axis elongation and brain expansion at E12.5. **(a-f)** Macro-anatomy quantification of live E12.5 embryos from double-heterozygote breeding, total n = 47 littermates from 6 breeding females. **(a)** A brightfield embryo image with highlighted measurements (see also **Suppl. Fig. S13d**). SC = spinal cord, FL = forelimb, HL = hindlimb, A = anterior, P = posterior, D = dorsal, V = ventral, M = medial, L = lateral. Some genotypes were pooled to secure sufficient biological replicates for statistical assessment: *Sorcs2(+/+);Ror2(+/+)* & *Sorcs2(+/-);Ror2(+/+)* [n = 4; 4 breeding females (f)]. *Sorcs2(+/+); Ror2(+/-)* [n = 4; 3f]. *Sorcs2(+/-);Ror2(+/-)* [n = 21; 6f], *Sorcs2(-/-);Ror2(+/+)* & *Sorcs2(-/-);Ror2(+/-)* [n = 7; 3f]. *Sorcs2(+/+); Ror2(-/-)* & *Sorcs2(+/-); Ror2(-/-)* [n = 8; 5f]. *Sorcs2(-/-); Ror2(-/-)* [n = 3; 3f]. **(b)** Naso-maxillary area representing craniofacial features. Ordinary one-way ANOVA (P = 0.5432) with Fisher’s LSD test. **(c-e)** Anterior-posterior proportions. **(c)** Body axis length, ordinary one-way ANOVA (P = 0.0011) with Fisher’s LSD test. **(d)** Trunk length, ordinary one-way ANOVA (P = 0.00001) with Fisher’s LSD test. **(e)** Brain length measured along the dorsal side of the brain, ordinary one-way ANOVA (P = 0.0479) with Fisher’s LSD test. **(f)** Forebrain-midbrain thickness, ordinary one-way ANOVA (P = 0.0107) with Fisher’s LSD test. **(g)** Representative image of a 3D-reconstructed E12.5 *Hb9:GFP/+* embryo (see **Suppl. video SV1)**, quantifications in **(h-j)**. *Hb9:GFP/+; Sorcs2(+/+)* [n = 10 embryos; 3 breeding females]; *Hb9:GFP/+;Sorcs2(-/-)* [n = 9 embryos; 2 breeding females]. **(g)** *Left panel:* Light-sheet microscopy image of E12.5 *Hb9:GFP/+;Sorcs2(+/+)* embryo. *Right panel:* Imaris-based segmentation of isolated areas of interest: spinal cord (Hb9+) and limb buds (autofluorescence in forelimb and hindlimb). **(h)** Quantifications of spinal cord length **(h1-2)** and curvature **(h3)**. Two-tailed Mann-Whitney test **(h2)**, two-tailed unpaired t-test **(h1, h3)**. **(i)** Quantifications of forelimb bud proportions. Pr = Proximal, Di = distal. Two-tailed Mann-Whitney test **(i3)**, two-tailed unpaired t-test **(i1-2, i4)**. **(j)** Quantifications of hindlimb bud proportions. Two-tailed Mann-Whitney test **(j1-2)**, two-tailed unpaired t-test **(j3-4)**. All graphs represent mean ± S.E.M. Source data are provided as a Source Data file.

SorCS2-dependent neural tube elongation at E12.5 was further examined by whole-mount light-sheet microscopy of transgenic *Hb9:GFP/+* embryos^58^, labeling all motor neurons. As both SorCS2^58^ and Ror2 are co-expressed in the embryonic spinal cord (**Suppl. Fig. S2c-d**), Hb9-based segmentation was used to quantify spinal cord morphogenesis, alongside autofluorescence-based segmentation of limb buds (**Fig. 6g, Suppl. Video SV1**). *Hb9:GFP/+; Sorcs2^-/-^* embryos exhibited an ∼500 µm increase in spinal cord length and abnormal curvature (**Fig. 6h**), while limb bud development remained unaffected (**Fig. 6i-j**), indicating neural specificity of SorCS2 function.

Together, our data demonstrates that SorCS2 loss induces a transient elongation of the anterior-posterior axis at E12.5, characterized by neural tube expansion and increased spinal cord length. Notably, similar phenotypes have been reported in embryos with ectopic Wnt1 expression^73^ or constitutive β-catenin activation^74^, both associated with hyperactivated canonical Wnt signaling. These observations suggest that the transient elongation observed in *Sorcs2^−/−^* embryos may reflect elevated Wnt/β-catenin activity at this developmental stage. This temporal phenotype resolves by E14.5 yet might contribute to the A-P axis truncation in adult *Sorcs2^−/−^* mice.

### Ror2 and SorCS2 form a receptor complex in floorplate mDA radial glia at E11.5

To investigate the role of Ror2-SorCS2 signaling in mDA lineage development, we first performed TaqMan qPCR of the ventral midbrain at E11.5, E12.5, and E14.5. While *Ror2* expression temporarily decreased by 50% between E11.5 and E12.5, *Sorcs2* expression doubled at E12.5 (**Fig. 7a-b**). These findings suggest dynamic regulation of *Ror2* and *Sorcs2* expression between E11.5 and E12.5, supporting a potential role for Ror2-SorCS2 signaling during this developmental window.

**Fig. 7:**
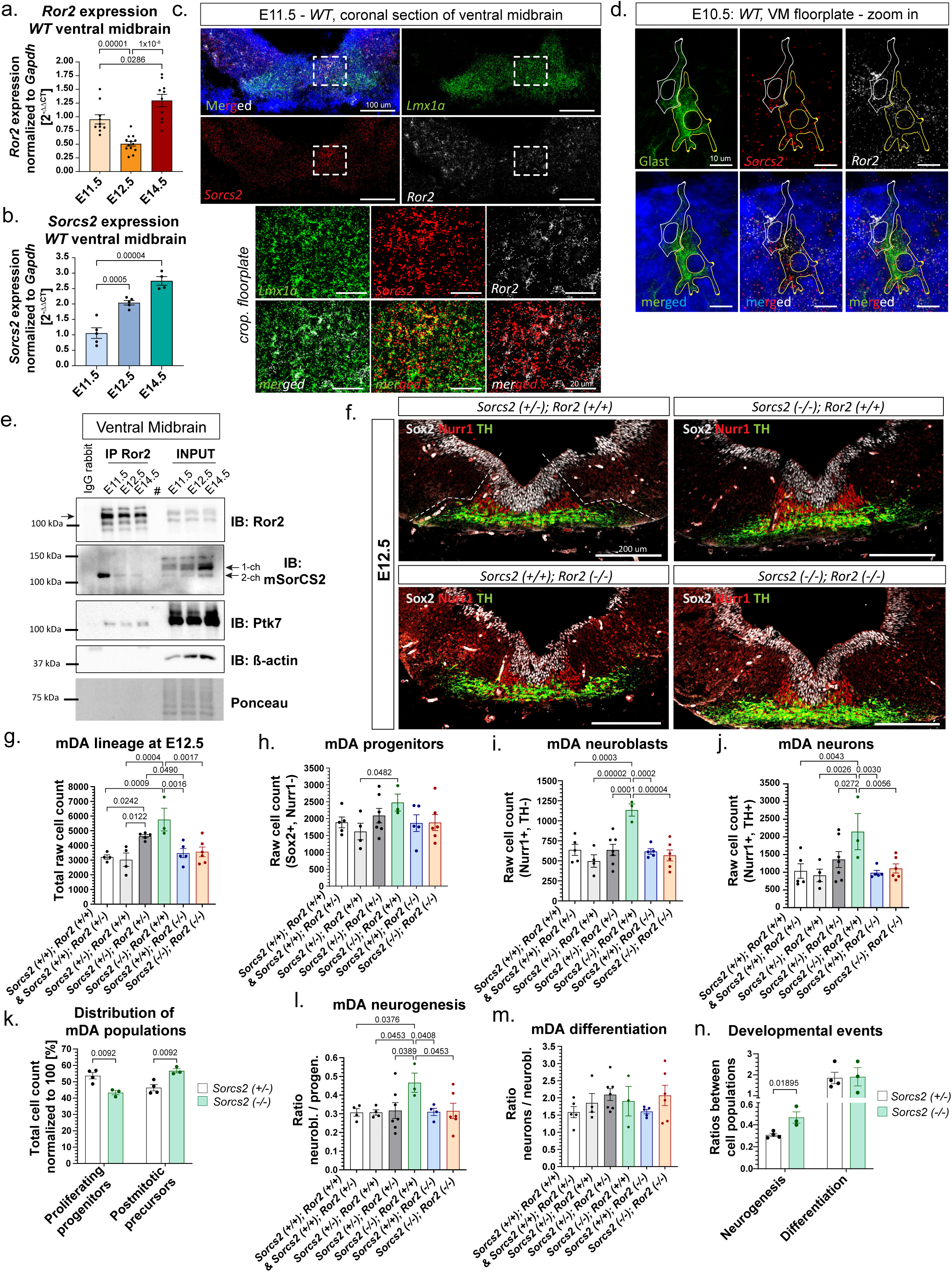
SorCS2 interacts with Ror2 in murine mDA progenitors to regulate their proliferation and neurogenesis. **(a-b)** TaqMan qPCR of *Ror2* **(a)** and *Sorcs2* **(b)** expression in mouse VM tissue, E11.5 to E14.5 littermates. One-way ANOVA [P = 4x10^-8^ **(a)**, P = 0.0001 **(b)**] with Fisher’s LSD test. **(a)** E11.5 (n = 11; 2 breeding females, f). E12.5 (n = 14; 3f). E14.5 (n = 10, 2f). **(b)** E11.5 (n = 5; 1f), E12.5 (n = 5; 1f), E14.5 (n = 5; 1f). **(c-d)** Representative fluorescent RNA-ISH in VM coronal sections, three biological replicates, three independent experiments. Hoechst (blue). **(c)** *Ror2* and *Sorcs2* were co-expressed in *Lmx1a+* mDA precursors at E11.5. Scale bars: 100 µm/20 µm (cropped). **(d)** *Ror2* and *Sorcs2* RNA-ISH combined with IF labeling of VM mDA radial glia [Glast+] at E10.5. Scale bar: 10 µm. **(e)** Representative IP-Ror2 and immunoblotting of E11.5–E14.5 VM tissue from three independent experiments. **(f-n)** mDA lineage cell-fate analysis in double-heterozygote breeding littermates at E12.5. Total n = 35 littermate embryos from 10 breeding females (f): *Sorcs2(+/+);Ror2(+/+)/Ror2(+/-)* [n = 5; 3f]. *Sorcs2(+/-);Ror2(+/+)* [n = 4; 4f]. *Sorcs2(+/-);Ror2(+/-)* [n = 7; 7f]. *Sorcs2(+/-);Ror2(-/-)* [n = 5; 5f]. *Sorcs2(-/-);Ror2(+/+)* [n = 3; 3f]. *Sorcs2(+/+);Ror2(-/-)* [n = 5; 3f]. *Sorcs2(-/-);Ror2(-/-)* [n = 6; 5f]. **(g-j, l-m)** Multiple-comparison Fisher’s LSD test. **(f)** Representative confocal images, mDA progenitors (white), postmitotic mDA neuroblasts (red), immature mDA neurons (green). Dashed borderline marks quantified mDA domain. **(g)** mDA lineage cell count, one-way ANOVA (P = 0.0047). **(h)** mDA progenitors, one-way ANOVA (P = 0.5510). **(i)** mDA neuroblasts, one-way ANOVA (P = 0.0010). **(j)** mDA neurons. One-way ANOVA (P = 0.0348). **(k)** Normalized distribution of mDA lineage in *Sorcs2(+/-);Ror2(+/+)* and *Sorcs2(-/-);Ror2(+/+)* showed a developmental shift between proliferative and postmitotic populations. Two-tailed multiple unpaired t-tests. **(l)** Neurogenesis index = ratio of mDA neuroblasts to progenitors. Brown-Forsythe ANOVA (P = 0.2288). **(m)** Differentiation index = ratio of mDA neurons to neuroblasts. One-way ANOVA (P = 0.2581). **(n)** Developmental events in *Sorcs2(+/-);Ror2(+/+)* and *Sorcs2(-/-);Ror2(+/+)* embryos. Two-tailed multiple unpaired t-tests. Graphs represent mean ± S.E.M. Source data are provided as a Source Data file.

Analysis of single-cell RNA sequencing data set from mouse ventral midbrain^75^ revealed that *Ror2* is expressed in all midbrain cell types, including mDA progenitors (i.e. radial glia), mDA neuroblasts, and mDA neurons (**Suppl. Fig. S15a-b**). *Sorcs2* expression was significantly enriched in radial glia, with fewer transcripts detected in mDA neuroblasts and mDA neurons (**Suppl. Fig. S15c**). While *Wnt5a* was significantly enriched in radial glia, mDA neuroblasts, and newborn mDA neurons (**Suppl. Fig. S14e**), *Ror1* transcripts were not detected (**Suppl. Fig. S15f**). Fluorescent multiplex ISH confirmed *Ror2* and *Sorcs2* co-expression in the mDA domain [*Lmx1a+*] at E10.5 and E11.5 (**Fig. 7c, Suppl. Fig. S16a**). Sequential IF labeling with the floorplate radial glia marker Glutamate Aspartate Transporter (Glast/EAAT1; gene *Slc1a3*, expression profile in **Suppl. Fig. S15d**) revealed *Ror2* and *Sorcs2* co-expression in newborn radial glia at E10.5, with increased cell number at E11.5 (**Fig. 7d, Suppl. Fig. S16b-c**).

Strikingly, Ror2 pulldown experiments using ventral midbrain tissue from E11.5, E12.5 and E14.5 *WT* embryos demonstrated a strong interaction between Ror2 and SorCS2 exclusively at E11.5 (**Fig. 7e**). These results indicate that the Ror2-SorCS2 complex forms in floorplate radial glia specifically at E11.5, potentially regulating their proliferation or neurogenesis.

### SorCS2 deficiency accelerates proliferation and neurogenesis of mDA progenitors

Adult *Sorcs2^-/-^* mice exhibit abnormal dopamine transmission and hyperactive-like behavior in adulthood^40, 41, 42^. To investigate the role of Ror2-SorCS2 signaling in mDA lineage development, we analyzed embryos at E12.5, a stage immediately following peak receptor interaction at E11.5, allowing assessment of early downstream developmental consequences. Coronal ventral midbrain sections were collected to quantify mDA progenitors [Sox2+, Nurr1-], mDA neuroblasts [Nurr1+, TH-], and immature mDA neurons [Nurr1+, TH+] by immunofluorescence (**Fig. 7f**).

While Ror2 loss alone did not affect mDA lineage development (**Fig. 7g-j, l-m**), total cell numbers were increased in *Sorcs2^+/-^; Ror2^+/-^* and *SorCS2^-/-^; Ror2^+/+^* embryos (**Fig. 7g**)*. Sorcs2^-/-^; Ror2^+/+^* embryos further showed an expanded pool of proliferating mDA progenitors (**Fig. 7h**), and a two-fold increase in postmitotic mDA neuroblasts and neurons (**Fig. 7i-j**). These effects were abolished in *Sorcs2^-/-^; Ror2^-/-^* embryos, indicating that SorCS2 restrains Ror2-dependent signaling during mDA proliferation and neurogenesis. Together, these findings indicate that SorCS2 deficiency enhances Ror2-dependent signaling, whereas Ror2 loss abrogates this effect.

Comparative population analysis revealed a reduction in proliferating mDA progenitors with a concomitant increase in postmitotic mDA precursors in *Sorcs2^-/-^; Ror2^+/+^* embryos compared to *Sorcs2^+/-^; Ror2^+/+^*littermates (**Fig. 7k**), indicating accelerated neurogenesis upon SorCS2 loss. Consistently, *Sorcs2^-/-^; Ror2^+/+^* embryos exhibited an elevated neurogenesis index (neuroblasts-to-progenitors ratio) that was rescued in *Sorcs2^-/-^; Ror2^-/-^* embryos (**Fig. 7l, n**), whereas the differentiation index (neurons-to-neuroblasts ratio) remained unchanged (**Fig. 7m-n**), supporting the selective role of SorCS2 in regulating Ror2-dependent cell cycle exit and mDA neurogenesis.

These findings suggest that SorCS2 deficiency increases Wnt signaling activity and that Ror2-SorCS2 signaling axis is essential for mDA progenitor proliferation and neurogenesis. The absence of phenotypes in *Ror2^-/-^; Sorcs2^+/+^* and *Ror2^-/-^; Sorcs2^-/-^* embryos indicates that concurrent loss of *Ror2* abolishes the SorCS2-dependent phenotype, demonstrating that SorCS2 functions to limit Ror2 activity in a context-dependent, spatiotemporally controlled manner during mDA lineage development.

### SorCS2 loss alters Wnt component levels, distribution, and activity, sensitizing mDA progenitors to Wnt exposure *in vitro*

While increased proliferation of *Sorcs2^-/-^*mDA progenitors is indicative of increased Wnt/β-catenin signaling^73^, accelerated neurogenesis suggests a simultaneous overactivation of both Wnt1- and Wnt5a-dependent signaling^30, 32, 33^ (**Fig. 10a**). Overexpression of SorCS2-FL in HEK293T cells increased basal Wnt/β-catenin activity by ∼60%, an effect reduced to ∼28% in presence of SorCS2-Tailless (**Fig. 8a**), indicating that SorCS2 also modulates Wnt/β-catenin signaling. To examine how impaired Ror2-SorCS2 signaling affects these two Wnt signaling cascades in mDA progenitors, we generated monoclonal *Ror2* and *Sorcs2* single-knockout (KO) and double-knockout (dKO) SN4741 cell lines using CRISPR/Cas9. Gene disruption was confirmed by immunoblotting (**Suppl. Fig. S18a**).

**Figure 8:**
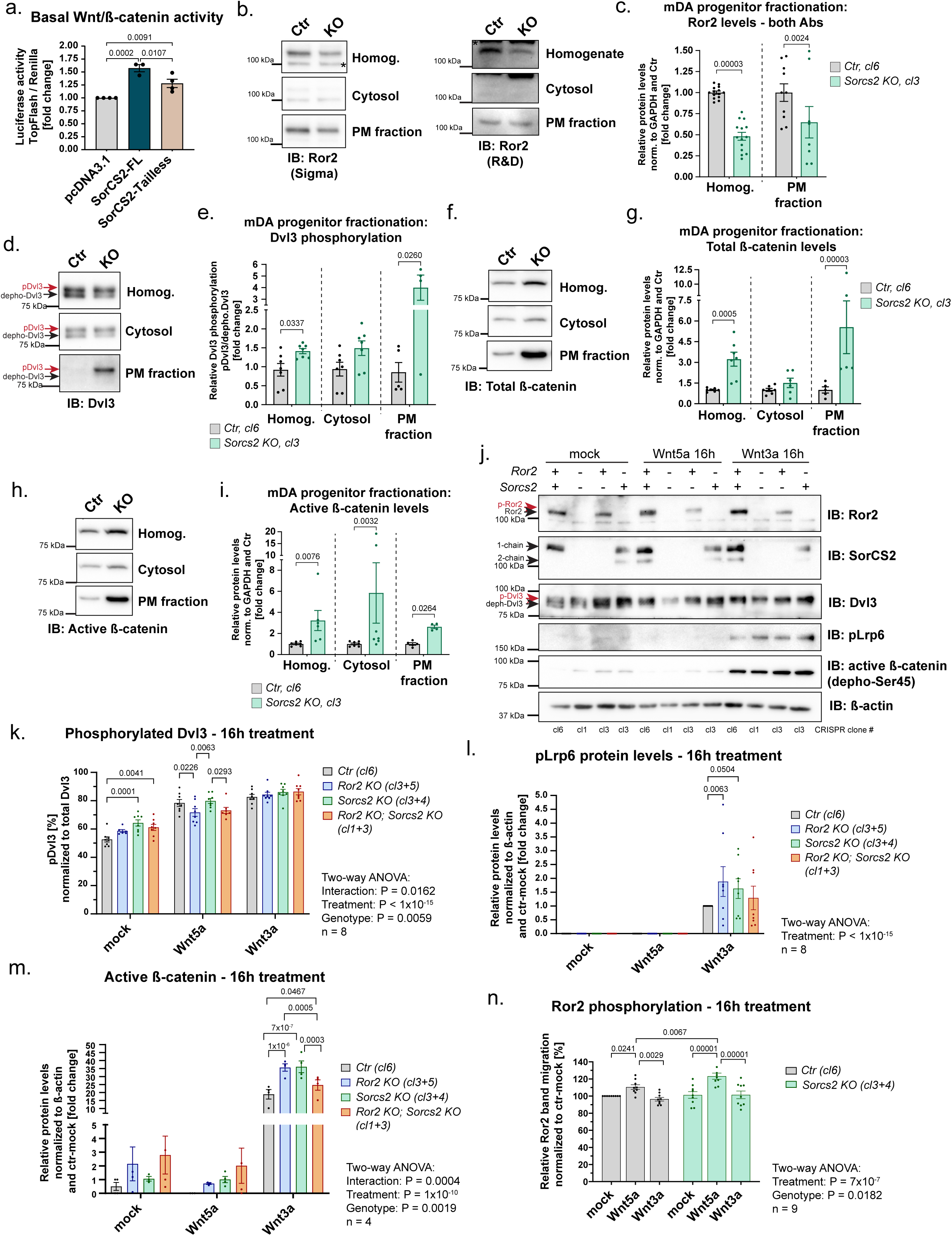
SorCS2 loss alters Wnt component levels, distribution, and activity, sensitizing mDA progenitors to Wnt exposure. **(a)** TopFlash dual luciferase assay in transfected HEK293T cells. Three independent biological experiments (n = 3); one-way ANOVA (P = 0.0006) with Fisher’s LSD test. **(b-i)** Subcellular fractionation and immunoblotting of control (*Ctr*) and *Sorcs2 KO* monoclonal mDA progenitor lines. Homog. = Homogenate, PM = Plasma Membrane. Five independent experiments with 1-2 biological replicate/group. Multiple two-tailed unpaired t-tests. Fractions were normalized to GAPDH and *Ctr* samples (**Suppl. Fig. S16a-b**). **(b-c)** Ror2 protein levels (*Ctr* n = 14; *Sorcs2 KO* n = 10). Results from two Ror2 antibodies **(b)** were combined for statistical analysis **(c). (d-e)** Dvl3 phosphorylation (*Ctr* n = 6; *Sorcs2 KO* n = 7, PM n = 4). **(f-g)** Total β-catenin (*ctr* n = 7, PM n = 5; *Sorcs2 KO* n = 7, PM n = 5). **(h-i)** Active β-catenin (*ctr* n = 7, PM n = 5; *Sorcs2 KO* n = 7, PM n = 5). **(j-n)** Immunoblotting of *Sorcs2* and *Ror2* KO or double-KO (dKO) lines after 16h Wnt exposure. Mock = PBS-0.1% BSA, recombinant human Wnt5a (100 ng/ml), or Wnt3a (100 ng/ml). Two independent CRISPR clones per genotype were pooled from 3-5 independent experiments; two-way ANOVA with Fisher’s LSD test. **(j-m)** Wnt3a amplified canonical Wnt/β-catenin signaling, seen as increased Dvl3 phosphorylation (n = 8) **(k)** and Lrp6 phosphorylation (n = 8) **(l)**, and accumulated active β-catenin (n = 4) **(m)**. Wnt5a stimulation amplified noncanonical Wnt signaling seen as enhanced Dvl3 phosphorylation **(k)** and Ror2 phosphorylation (n = 9) **(n)** (see also **Suppl. Fig. S18h)**. *Ror2 KO* and *Sorcs2 KO* lines exhibited increased Wnt/β-catenin signaling upon Wnt3a, which was reduced in *Ror2; Sorcs2 dKO* line **(k, m)**. **(k)** *Sorcs2 KO* and *Ror2; Sorcs2 dKO* lines exhibited increased Dvl3 phosphorylation in mock, indicating enhanced basal Wnt signaling. *Ror2 KO* and *Ror2; Sorcs2 dKO* lines showed reduced Dvl3 phosphorylation upon Wnt5a. *Sorcs2 KO* line exhibited Wnt5a-induced Ror2 hyperphosphorylation **(n)**. Graphs represent mean ± S.E.M. Source data are provided as a Source Data file.

Because specific antibodies suitable for immunofluorescent detection of endogenous Ror2 are unavailable, we performed subcellular fractionation of control (*Ctr)* and *Sorcs2 KO* lines to quantify Ror2 and other Wnt components in homogenate, cytosol, and plasma membrane (PM) fractions using immunoblotting. *Sorcs2 KO* mDA progenitors showed a ∼50 % reduction in Ror2 protein levels in both homogenate and PM fraction (**Fig. 8b-c, Suppl. Fig. S17a-b**). Although total Dvl3 levels were unchanged, Dvl3 phosphorylation was increased in both homogenate and PM fractions in *Sorcs2 KO* mDA progenitors (**Fig. 8d-e, Suppl. Fig. S17c**), indicating enhanced Wnt signal transduction at the plasma membrane. *Sorcs2 KO* cells also displayed elevated total Lrp6 levels in homogenate, with no detectable changes in PM localization or phosphorylation (**Suppl. Fig. S17d-g**).

Additionally, SorCS2 loss resulted in a ∼200% increase in total β-catenin in both homogenate and PM fractions (**Fig. 8f-g**), accompanied by up to threefold increase in activated β-catenin across all fractions (**Fig. 8h-i**). Together, these data indicate that SorCS2 deficiency elevates basal Wnt signaling activity in mDA progenitors, likely through impaired sorting of Wnt pathway components, including Ror2.

To assess how the *Ror2* and *SorCS2* mutant lines respond to distinct Wnt stimuli, mDA progenitors were treated with mock (PBS + 0.1% BSA), recombinant human Wnt5a (inducing Ror2 and Dvl3 phosphorylation), or Wnt3a (inducing Lrp6 and Dvl3 phosphorylation, and β-catenin activation) for 2 hours (acute stimulation) (**Suppl. Fig. S18a-e, i**) or 16 hours (chronic stimulation) (**Fig. 8j-n, Suppl. Fig. S18f-h**).

Under mock conditions, *Sorcs2 KO* and *Ror2; Sorcs2 dKO* lines displayed elevated Dvl3 phosphorylation, confirming increased basal Wnt activity upon SorCS2 loss (**Fig. 8k, Suppl. Fig. S18d**). Acute Wnt3a treatment produced no significant effects (**Suppl. Fig. S18b-e**). In contrast, chronic Wnt3a stimulation enhanced Wnt/β-catenin signaling in *Ror2 KO* and *Sorcs2 KO* lines, respectively, as shown by increased levels of phosphorylated Lrp6 and active β-catenin (**Fig. 8l-m**). These results demonstrate that disruption of Ror2-dependent signaling influences both noncanonical and canonical Wnt pathways, and that SorCS2 deficiency sensitizes mDA progenitors to Wnt3a stimulation.

Conversely, chronic Wnt5a stimulation reduced Dvl3 phosphorylation in *Ror2 KO* and *Ror2; Sorcs2 dKO* lines (**Fig. 8k**), consistent with impaired noncanonical Wnt signaling in the absence of Ror2^10, 12, 36^. Notably, *Sorcs2 KO* cells displayed Wnt5a-dependent Ror2 hyperphosphorylation following both acute and chronic stimulation, assessed by increased Ror2 electrophoretic mobility^76, 77^ (**Fig. 8n, Suppl. Fig. S18h-i**), despite reduced Ror2 levels at the plasma membrane (**Fig. 8c**). These findings suggest that SorCS2 promotes internalization of phosphorylated Ror2 upon Wnt5a binding and that SorCS2 deficiency sensitizes mDA progenitors to Wnt5a stimulation.

Together with the mDA cell fate data (**Fig. 7**), our findings show that SorCS2 deficiency in mDA progenitors likely impairs Ror2 internalization and sorting, resulting in increased noncanonical Wnt signaling, as exemplified by elevated Wnt5a-dependent Ror2 and Dvl3 phosphorylation. In parallel, SorCS2 loss increases Lrp6 and β-catenin levels and enhances their responsiveness to prolonged Wnt3a stimulation, indicating elevated canonical Wnt/β-catenin activity. Thus, SorCS2 functions as a regulator of Wnt receptor sorting that limits excessive Ror2 signaling and maintains balanced noncanonical and canonical Wnt pathway activity.

### SorCS2 loss does not cause global early endosome defects in the mDA lineage *in vivo*

Next, we asked whether SorCS2 loss globally affects endocytic internalization within the mDA lineage. We examined SorCS2 co-localization with early endosomes [EEA1+] in the VM floorplate at E12.5. SorCS2 was enriched in mDA radial glia [Glast+], highly polarized ventricular zone progenitors, with apically positioned somata facing the neural tube cavity and elongated processes extending through the intermediate to marginal zones toward the meninges^32^. Early endosomes were predominantly localized apically within these cells (**Fig. 9a-c**). SorCS2 co-localized with Glast at the plasma membrane (arrowheads) and with intracellular sorting vesicles, including EEA1+ early endosomes (arrows) (**Fig. 9b**).

**Figure 9:**
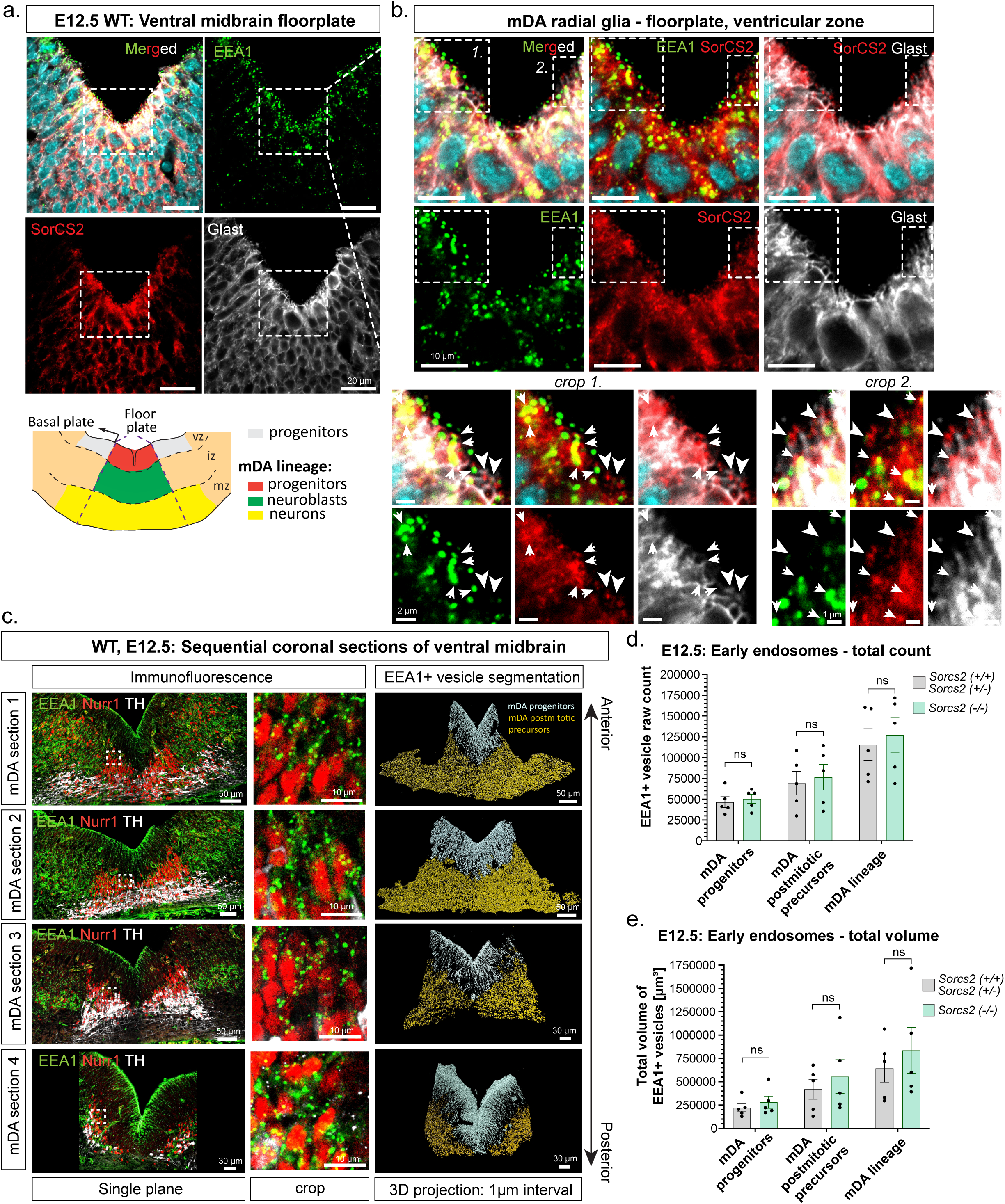
SorCS2 loss does not cause early endosome defects in the mDA lineage *in vivo.* **(a-b)** Representative confocal images of SorCS2 immunofluorescent labeling with radial glia marker Glast and early endosome marker EEA1 in coronal sections of wild-type E12.5 VM floorplate. Hoechst (cyan), three independent experiments and three biological replicates. SorCS2 is expressed in Glast+ mDA progenitors of the ventricular zone (vz) of the VM floorplate, where EEA1+ endosomes are enriched apically towards the neural tube cavity **(a)**. Scale bar: 20 µm. **(b)** SorCS2 co-localizes with Glast at the plasma membrane (arrowheads) and with EEA1+ endosomes near the plasma membrane (arrows). Scale bars: 10 µm/2 µm (crop). **(c-e)** 3D reconstruction and quantification of EEA1+ endosomes in the E12.5 mDA domain. Embryos derived from *Sorcs2+/-* breeding: *Sorcs2+/+* & *Sorcs2+/-* (n = 5 embryos; 3 breeding females); *Sorcs2-/-* (n = 5 embryos; 3 breeding females). **(c**) Representative 1-µm-interval confocal Z-stacks and Imaris-based 3D reconstructions of EEA1+ endosomes. Vesicles were reconstructed along the anterior–posterior VM axis from four sequential, 14-µm thick coronal sections. Nurr1 and TH labeling defined the mDA domain (**c**; *right panel* - segmentation), including postmitotic precursors [Nurr1+] and progenitors [Nurr1-]. EEA1+ structures associated with blood vessels and meninges were excluded based on autofluorescence. Scale bars: 50/30 µm; crops 10 µm. **(d-e)** Total EEA1+ vesicle number **(d)** and volume **(e)** across all sections did not differ between genotypes. Two-tailed multiple unpaired t-tests. Graphs represent mean ± S.E.M. Source data are provided as a Source Data file.

Using 3D confocal microscopy combined with Imaris-based segmentation, we assessed whether *Sorcs2^-/-^* embryos exhibit impaired endocytic vesicle formation compared to control littermates. EEA1+ early endosomes were 3D-reconstructed within the mDA lineage, and vesicle number and volume were quantified in proliferating mDA progenitors [Nurr1-, TH-] and postmitotic mDA precursors [Nurr1+; TH+] (**Fig. 9c, Suppl. Video SV2**), analyzed either separately or combined (**Fig. 9d-e**). Vesicles originating from blood vessels or meninges were excluded based on autofluorescence. *Sorcs2^-/-^* embryos showed no differences in early endosome number or volume, indicating that SorCS2 loss does not globally impair endocytic vesicle formation. These data support a selective SorCS2 effect on Ror2 internalization following Wnt5a-dependent phosphorylation.

### *Sorcs2^-/-^* embryos display elevated *Wnt1*, *Wnt5a,* and *Tcf4* expression, and hyperactivated Ror2-dependent Wnt signaling

Precise spatiotemporal control of Wnt signaling activity in ventral midbrain, exemplified by dynamic *Wnt1* and *Wnt5a* expression, shapes mDA progenitor pools and neuronal diversity essential for correct dopaminergic innervation. Between E11.5 and E14.5, Wnt1 promotes mDA progenitor proliferation, whereas Wnt5a provides inhibitory cues that, together with Wnt1, synergistically promote mDA neurogenesis (**Fig. 10a**)^30, 32, 36, 39, 78^.

**Figure 10:**
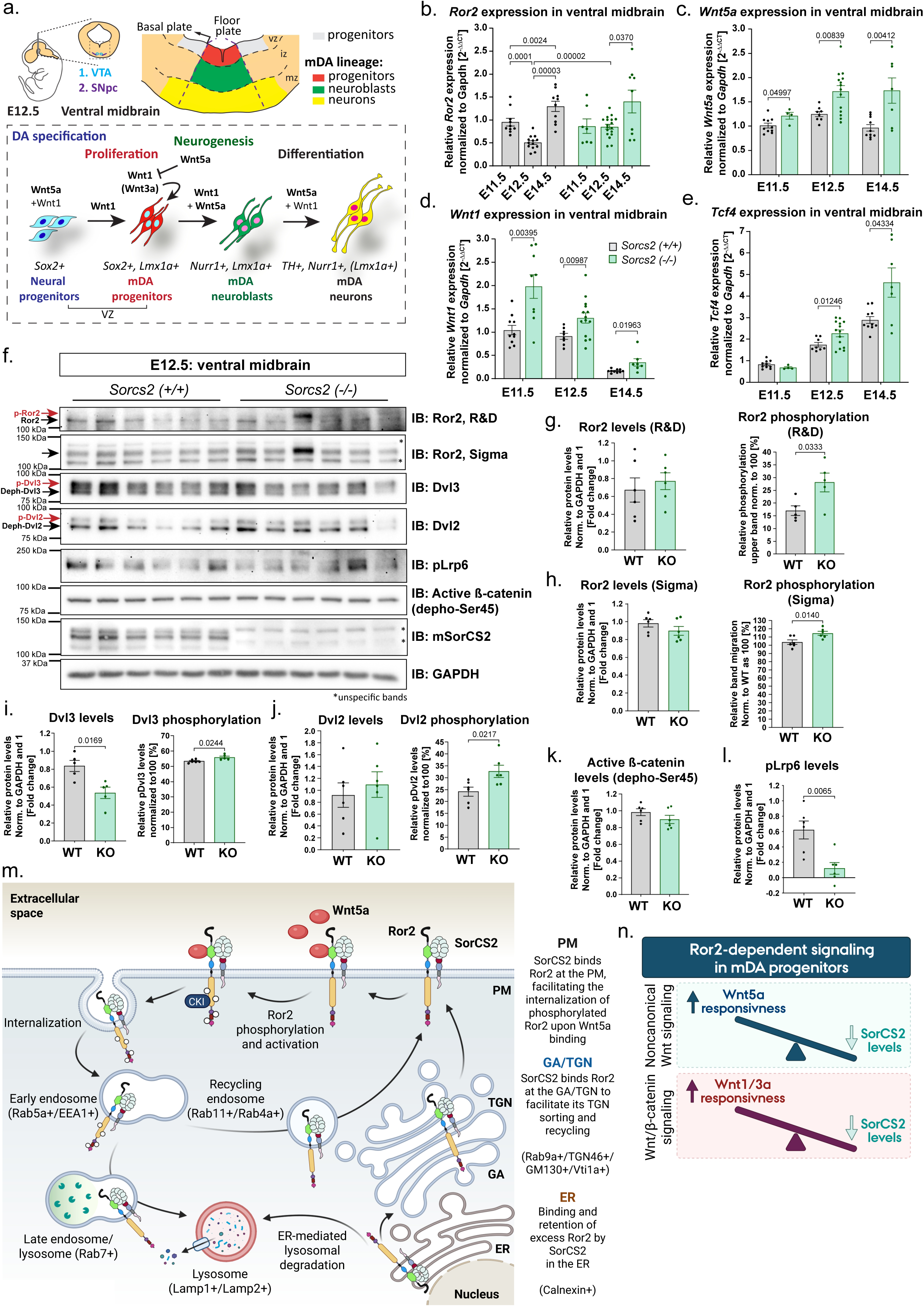
S*o*rcs2*^-/-^* embryos display altered transcription of Wnt components and hyperactivated Ror2-dependent Wnt signaling. **(a)** Schematic of Wnt1- and Wnt5a-dependent signaling during mDA lineage development (E10.5 - E14.5)^33^. **(b-e)** TaqMan qPCR analysis of VM tissue. *WT* and *Sorcs2 KO* littermate embryos. Normalized to *Gapdh* and to mean *Sorcs2(+/+)* E11.5. **(b)** *Ror2* expression, mixed-effects analysis [P_age_ = 0.000002, P_genotype_ = 0.2919, P_interaction_ = 0.0936] with Fisher’s LSD test. E11.5: WT (n = 11; 2 breeding females, f), KO (n = 8; 2f). E12.5: WT (n = 14; 3f), KO (n = 19; 3f). E14.5: WT (n = 10; 2f), KO (n = 9; 1f). **(c)** *Wnt1* expression. E11.5: WT (n = 10; 2f), KO (n = 9; 3f). E12.5: WT (n = 8; 2f), KO (n = 14; 3f). E14.5: WT (n = 10; 2f), KO (n = 9; 1f). **(d)** *Wnt5a* expression. E11.5: WT (n = 10; 2f), KO (n = 4; 2f). E12.5: WT (n = 8; 2f), KO (n = 14; 3f). E14.5: WT (n = 10; 2f), KO (n = 9; 1f). **(e)** *Tcf4* expression. E11.5: WT (n = 10; 2f), KO (n = 4; 2f). E12.5: WT (n = 8; 2f), KO (n = 14; 3f). E14.5: WT (n = 10; 2f), KO (n = 9; 1f). **(c-e)** Two-tailed multiple unpaired t-tests. **(f-l)** Wnt components immunoblotting analysis, VM tissue, E12.5 littermate embryos. Six independent biological replicates per genotype [*Sorcs2(+/+)* n = 6 embryos; 1 breeding female. *Sorcs2(-/-)* n = 6; 1f], analyzed in parallel and quantified in **(g-l)**, two-tailed lognormal Welch’s t-test. **(g-h)** Ror2 protein and phosphorylation levels. Antibodies recognized Ror2 N-terminus **(g)** and C-terminus **(h)**. Quantification of Dvl3 **(i)** and Dvl2 **(j)** protein and phosphorylation levels. **(k)** Quantification of active β-catenin. **(l)** Quantification of pLrp6. Graphs represent mean ± S.E.M. Source data are provided as a Source Data file. **(m)** Model of SorCS2-dependent regulation of Ror2. SorCS2 facilitates endosomal trafficking, Wnt-dependent internalization, and lysosomal degradation of Ror2, thereby establishing a negative feedback loop that prevents Wnt signaling hyperactivation. **(n)** Schematic of SorCS2 deficiency effects on Wnt signaling in mDA progenitors. **(m-n)** Created with BioRender.

Since Ror2 signaling has been associated with positive feedback regulation of Ror2 and Wnt ligand expression^79^, we first examined whether SorCS2 deficiency alters Ror2 expression dynamics by TaqMan qPCR. Ror2 levels were comparable between genotypes at E11.5 and E14.5. However, while *Sorcs2^+/+^* embryos exhibited an approximately ∼50% reduction in Ror2 expression at E12.5, *Sorcs2^−/−^* embryos failed to undergo this temporal downregulation (**Fig. 10b**). Given that SorCS2 and Ror2 interact at E11.5, these transcriptional changes at E12.5 likely occur downstream of their disrupted Ror2-SorCS2 interaction and may reflect enhanced signaling feedback.

Consistent with altered *Ror2* transcript regulation, SorCS2 deficiency also resulted in elevated *Wnt5a* and *Wnt1* expression in VM tissue at E11.5, E12.5, and E14.5 (**Fig. 10c-d**). Together, these findings indicate that SorCS2 deficiency alters both Wnt receptor and ligand expression dynamics within the ventral midbrain, potentially amplifying Wnt signaling output during critical windows of mDA progenitor expansion and neurogenesis. Whether the increased expression reflects compensatory feedback or direct transcriptional regulation by SorCS2 remains unresolved.

To assess downstream transcriptional consequences of altered SorCS2-dependent mDA development, we analyzed expression of the basic helix–loop–helix transcription factor 4 (*Tcf4*)^80, 81^. *Tcf4* is enriched in the mDA lineage, particularly in progenitors and neuroblasts (**Suppl. Fig. S15g–h**). SorCS2 loss resulted in elevated *Tcf4* expression at E12.5 and E14.5 (**Fig. 10e**), supporting transcriptional deregulation downstream of SorCS2-modulated Wnt signaling. Given that *TCF4* is a major susceptibility gene for neurodevelopmental psychiatric disorders and has been associated with impaired neurogenesis and Wnt signaling^82, 83, 84^, these findings provide a disease-relevant molecular context, likely downstream of altered Ror2-SorCS2 signaling.

Finally, we analyzed Wnt signaling activity in *Sorcs2^+/+^*and *Sorcs2^-/-^* VM tissue at E12.5 by immunoblotting. To quantify Wnt5a-dependent Ror2 phosphorylation, we included an additional antibody (“R&D”) in which the higher, upper band corresponds to p-Ror2^85^ (**Fig. 10f**). While total Ror2 protein levels were unchanged, both Ror2 antibodies revealed increased Ror2 phosphorylation in *Sorcs2^-/-^* embryos (**Fig. 10g-h**). Moreover, despite reduced total Dvl3 levels, *Sorcs2^-/-^* embryos exhibited increased phosphorylation of Dvl3 and Dvl2 (**Fig. 10i-j**).

In parallel, active β-catenin levels were modestly reduced (**Fig. 10k**), although this did not reach statistical significance, probably reflecting tissue heterogeneity and/or a temporal delay in β-catenin accumulation relative to upstream signaling changes. Notably, *Sorcs2^-/-^* embryos showed an approximately 75-90% reduction in Lrp6 phosphorylation (**Fig. 10l**), consistent with enhanced Wnt5a/Ror2-dependent signaling and increased inhibition of Lrp6 activity, leading to accelerated neurogenesis of mDA progenitors at E12.5 (**Fig. 7k-l, 10a**).

These data are consistent with our *in vitro* observations and demonstrate that SorCS2 loss sensitizes mDA progenitors to Wnt1/3a and Wnt5a stimulation, resulting in enhanced Ror2-dependent Wnt signaling during mDA progenitor proliferation and neurogenesis (schematized in **Fig. 10n**).

## Discussion

This study identifies SorCS2 as a sorting-based regulator of Wnt signaling homeostasis that controls the localization, trafficking, activity, and function of the core Wnt receptor Ror2. Rather than acting as a classical co-receptor, SorCS2 limits excessive Ror2 activity by regulating receptor residence across subcellular compartments, including ER retention and endosomal and lysosomal trafficking. This level of control constrains Ror2 accumulation at the plasma membrane and restricts signal duration and amplitude *in vitro* and *in vivo.* By titrating ligand-induced Ror2 activation, SorCS2 enables tight control of Ror2-dependent signaling, thereby modulating and constraining both noncanonical and canonical Wnt pathway activity.

In the **Fig. 10m**, we propose a model of SorCS2-dependent regulation of Ror2 activity, trafficking and degradation in vertebrates. SorCS2 facilitates Ror2 endosomal trafficking, internalization, and lysosomal degradation, establishing a negative feedback loop that prevents Wnt signaling hyperactivation. SorCS2 retains excess Ror2 in the endoplasmic reticulum (ER) and targets it for lysosomal degradation. Physiological levels of the Ror2-SorCS2 receptor complex are trafficked through the Golgi apparatus (GA) and *trans*-Golgi network (TGN) to the plasma membrane (PM), where Ror2 becomes available for signaling. Upon Wnt5a binding, Ror2 undergoes sequential phosphorylation, initiating signal transduction. To limit Wnt5a-Ror2 activity, SorCS2 subsequently mediates internalization of phosphorylated Ror2 for endosomal recycling or lysosomal degradation, thereby maintaining homeostatic activity of Ror2-dependent Wnt signaling.

The Ror2-SorCS2 interaction depends on the extracellular CRD domain. Deletion of intracellular regions of either Ror2 (Δ745, Δ469) or SorCS2 (SorCS2-Tailless) did not abolish their physical interaction or co-localization, indicating that binding is mediated through extracellular domains of both receptors. The SorCS2 cytoplasmic tail appears to modulate this interaction indirectly by controlling receptor localization and trafficking rather than serving as a direct binding interface. The sorting motifs within the SorCS2 C-terminal tail, which are required for efficient receptor targeting^45^, may therefore modify Ror2 subcellular distribution and interaction dynamics, potentially through recruitment of additional regulatory factors.

SorCS2 predominantly localizes to sorting vesicles, where it regulates intracellular trafficking and targeting of its cargo proteins, including several tyrosine kinase receptors^45^. Like Ror2^7, 14, 59^, SorCS2 also localizes to actin-rich membrane protrusions that function as signaling hubs, including filopodia, axon growth cones, dendritic spines, and postsynaptic densities^40, 42, 45, 86^. Within these microdomains, SorCS2 has been linked to Rac1- and PKC-dependent signaling^44^, known modulators of both canonical and noncanonical Wnt signaling pathways^87^. This spatial organization suggests that SorCS2 not only regulates Ror2 trafficking but may also influence receptor availability or polarity within specialized signaling compartments, potentially extending its regulatory scope to additional Wnt signaling components, for example Wnt5a, Ror1, Ptk7, Vangl2, Lrp6, or β-catenin. However, the extent and specificity of SorCS2-dependent regulation of other Wnt signaling components remains to be determined.

In addition to regulating receptor trafficking, SorCS2 loss is associated with increased expression of *Wnt1, Wnt5a, Ror2*, and the Wnt-responsive transcription factor *Tcf4*, indicating broader effects on Wnt signaling output. Ror2 expression and signaling is tightly associated with Wnt5a ligand expression, which is regulated in a positive feedback loop^79^. Given that SorCS2 and Ror2 interact at E11.5, the transcriptional changes at E12.5 likely arise downstream of Ror2-SorCS2 interaction, possibly reflecting altered signaling feedback.

Elevated ligand expression, together with increased responsiveness of *Sorcs2^-/-^* mDA progenitors to Wnt3a and Wnt5a exposure, suggests that SorCS2-dependent sorting influences both ligand availability and downstream Wnt signaling activity. This is further supported by increased β-catenin and Lrp6 levels in *Sorcs2^-/-^* progenitors, consistent with enhanced Wnt signaling activity rather than a switch between the canonical and noncanonical Wnt pathways.

These molecular alterations may contribute to the transient anterior-posterior axis elongation observed in the brain and spinal cord of *Sorcs2^-/-^* embryos at E12.5. In developmental contexts like the ventral midbrain, where Wnt signals must be precisely titrated, even moderate dysregulation may promote excessive Wnt1-driven proliferation, premature cell-cycle exit driven by Wnt5a-mediated inhibition of Wnt/β-catenin signaling, and accelerated neurogenesis resulting from higher activity of both Wnt1 and Wnt5a^30, 36, 73, 78^. Nevertheless, whether the altered expression profiles arise from compensatory feedback mechanisms or direct transcriptional regulation by SorCS2 remains unresolved.

In mouse ventral midbrain, SorCS2 deficiency accelerates mDA proliferation and neurogenesis through enhanced responsiveness to Wnt1 and Wnt5a, whereas concurrent loss of Ror2 abolishes this effect. However, despite the rescue of mDA proliferation and neurogenesis, *Ror2-/-; Sorcs2-/-* embryos still display expansion of the brain at E12.5, suggesting that additional aspects of brain morphogenesis remain altered and likely involve signaling mechanisms beyond those directly responsible for the rescued phenotypes. This indicates that the proliferation and neurogenesis phenotypes in *Sorcs2^-/-^* embryos are mediated predominantly through Ror2-dependent signaling, supporting a model in which SorCS2 regulates Ror2-dependent Wnt signaling during neurodevelopment.

In mDA progenitors, Ror2 impairment is associated with increased Lrp6 phosphorylation and β-catenin activity upon chronic Wnt3a exposure, suggesting that elevated Wnt/β-catenin signaling in *Sorcs2^-/-^*progenitors may arise because of altered Ror2 signaling. Consistent with our observations, several studies have reported that Wnt3a-induced Ror2 activation enhances Lrp6 phosphorylation, β-catenin stabilization, and canonical Wnt target gene expression^23, 24, 25, 29^, while subsequent Wnt5a-mediated inhibition of canonical Wnt/β-catenin signaling does not reduce the pool of stabilized β-catenin^22^. Together, these findings demonstrate that disruption of Ror2 signaling can affect both noncanonical and canonical Wnt/β-catenin pathways^7, 14, 21, 22,23, 24, 25, 26^, likely through crosstalk at shared receptor complexes, including Ror2-Lrp6 co-assembly within signalosomes^14, 21, 27, 28^. Indeed, our data do not indicate that Ror2 directly activates canonical Wnt/β-catenin signaling in mDA progenitors. Rather, our findings demonstrate that disruption of the Ror2-SorCS2 axis alters cellular responsiveness to Wnt1, Wnt3a, and Wnt5a, thereby modulating both canonical and noncanonical Wnt signaling. Nevertheless, a potential direct contribution of SorCS2 to canonical Wnt/β-catenin signaling beyond Ror2, and in other contexts, remains to be determined.

Functional assays in mice, zebrafish, and *X. laevis* indicate that SorCS2-dependent receptor sorting regulates Ror2 signaling during convergent extension, somitogenesis, and brain morphogenesis, supporting an evolutionarily conserved role in vertebrate development. Differences in SorCS2-dependent phenotypic outcomes, such as axial truncation in zebrafish embryos versus transient elongation in mouse embryos, likely reflect species-, tissue-, and stage-specific contexts. Nevertheless, our loss- and gain-of-function analyses indicate that Wnt signaling outputs depend on homeostatic Ror2-SorCS2 interactions rather than absolute receptor levels. This is consistent with the well-established context dependence of Wnt signaling, where perturbations in core components, including Ror2, β-catenin, Dvl, and Lrp6, produce diverse phenotypes across models^6, 14^. This diversity underlies Wnt signaling complexity and reflects differences in tissue mechanics, cellular context, composition of ligand-receptor interactome, spatiotemporal expression patterns, concentration-dependent Wnt activity, and species-specific gene repertoires (e.g., 19 Wnt genes in mammals versus ∼25 in zebrafish)^88^.

Importantly, our data consistently supports the role for SorCS2 in regulating Ror2 signaling dynamics in a spatiotemporal manner. In Xenopus, SorCS2 attenuates Ror2 overactivation; in mouse, SorCS2 deficiency leads to Ror2 hyperactivation; and in zebrafish, physiological Ror2-SorCS2 signaling is required for proper convergent extension and brain morphogenesis. Although Ror2 activity cannot be directly measured in zebrafish, these findings are consistent with a conserved, context-dependent role of SorCS2 in fine-tuning Ror2 signaling during vertebrate development.

Notably, neither *Ror2^-/-^* nor *Ror2^-/-^; Sorcs2^-/-^* mice exhibit impaired dopaminergic fate, suggesting that constitutive Ror2 loss engages compensatory mechanisms during dopaminergic lineage development. Consistent with this interpretation, combined loss of *wnt5b* and *sorcs2* in zebrafish embryos does not exacerbate CE defects, indicating that in certain tissues and developmental contexts, noncanonical Wnt signaling may proceed through parallel, SorCS2-independent pathways, potentially involving alternative Wnt5a receptors such as Frizzled, Ryk, or Celsr family members^4, 7, 14^ or other members of the VPS10p-domain receptor family.

Indeed, all VPS10p-domain receptors are dynamically expressed during vertebrate development, regulating proliferation, neuronal outgrowth, survival, and cell death across healthy and diseased nervous systems^45, 58^. This suggests that the regulation of Wnt signaling may extend beyond SorCS2 to other VPS10p-domain receptors. Because the expression patterns of SorCS2, Ror2, and Wnt5a do not always overlap, future studies should systematically define whether and how individual VPS10p-domain receptors modulate Wnt signaling in a context- and concentration-dependent manner and whether disease-associated mutations disrupt such regulatory interplays^52, 89, 90^. Elucidating these mechanisms may provide insight into disease pathogenesis and inspire new therapeutic opportunities.

Disrupted Wnt signaling has been implicated in neuropsychiatric disorders associated with dysfunctional dopaminergic circuits, including autism spectrum disorder, schizophrenia, ADHD, and bipolar disorder^91, 92, 93^. Consistent with this, *SORCS2* has emerged as a risk gene for several of these conditions^40, 42, 47, 48, 49, 50, 51, 52^. We previously demonstrated that SorCS2 deficiency alters mDA neuron innervation and firing in adult mice, leading to hyperactivity-like, risk-taking, and stimulus-seeking behaviors^41, 42^. Although *Sorcs2^-/-^* embryos exhibit an expanded pool of mDA precursors at E12.5, adult mice display a normal mDA neuron number, indicating elimination of excess cells during circuit maturation^32, 42^. Notably, Wnt5a exerts stage-dependent effects on dopaminergic axons, initially promoting outgrowth before acting as a chemorepellent that induces growth cone retraction^31^, a switch reminiscent of SorCS2-dependent regulation^42, 44^. Together with evidence that SorCS2 promotes axon outgrowth in mDA neurons and motor neurons^42, 58^, our data suggest that the Wnt5a-Ror2-SorCS2 axis contributes to the establishment and targeting of mesocortical, mesostriatal, and mesolimbic circuits, although whether these effects arise directly from altered cell fate decisions or from independent roles in axon guidance remains to be determined.

We further identify increased expression of the Wnt-responsive transcription factor *Tcf4* downstream of SorCS2-dependent Wnt hyperactivation, providing a mechanistic link between altered Wnt signaling dynamics and transcriptional programs implicated in neurodevelopmental disease. *TCF4* is a major susceptibility gene for schizophrenia, Pitt–Hopkins syndrome, autism spectrum disorder, and bipolar disorder, and its dysregulation has been associated with abnormal neurogenesis, dopaminergic subtype specification, and altered Wnt signaling^80, 81, 84^. Together, these findings suggest that SorCS2-dependent control of Wnt signaling during embryonic development intersects with transcriptional programs linked to neuropsychiatric risk, potentially influencing long-term maturation and functional organization of dopaminergic circuits.

Our overexpression analyses revealed that ectopic Ror2 preferentially interacts with the SorCS2 single-chain, whereas endogenous Ror2 associates with the two-chain isoform, likely reflecting differences in cell type, protein abundance, or SorCS2 isoform stoichiometry. Although overexpression can perturb protein homeostasis, this approach allowed examination of how altered receptor levels affect Wnt signaling dynamics. Using this system, we show that SorCS2 can counteract excessive Ror2 activity by promoting its lysosomal degradation through both anterograde trafficking from the ER and endocytic sorting at the plasma membrane. This mode of regulation is particularly relevant in pathological contexts, as Ror2 hyperactivation has been linked to increased proliferation and invasiveness in several cancers while exhibiting tumor-suppressive functions in others^14, 21, 79^. Consistent with this model, SorCS2 has recently been identified as a tumor suppressor in ovarian cancer, where reduced expression correlates with abnormal Wnt signaling, increased tumor growth, and poor clinical outcome^94^. Together, these findings support a broader role for SorCS2 in constraining pathological Wnt signaling states and suggest that modulating SorCS2-dependent receptor sorting may represent a novel strategy to normalize deregulated Wnt signaling.

In conclusion, this study identifies a SorCS2-dependent mechanism that spatially and temporally restricts Wnt signaling activity. By controlling Ror2 trafficking, internalization, and degradation, SorCS2 regulates noncanonical and canonical Wnt signaling and prevents excessive pathway activation within critical developmental windows. These findings highlight receptor sorting pathways as promising therapeutic targets in disorders associated with dysregulated Wnt signaling and SorCS2 dysfunction.

## Methods

### Mouse lines genetics and husbandry

All breeding and experiments were approved by the Swedish (Stockholm’s Norra Djurförsöketiska Nämnd N158/15) and Danish Animal Experiments Inspectorate (permits 2016-15-0202-00051, 2017-15-0201-01192, 2021-15-0201-00850) and adhered to national and European guidelines (Directive 86/609/EEC, FELASA). Mice were bred and group-housed under 12 h light/dark cycles with *ad libitum* access to standard chow and water. Genotypes were confirmed by genomic PCR. The following lines were used: C57BL/6j BomTac (*WT*, Taconic), *Sorcs2^-/-^* (*Sorcs2 KO*)^42^, *Ror2^+/-^* (*Ror2 Htz*)^72^, *Hb9:GFP/^+^; Sorcs2^+/+^*^58^, and *Hb9:GFP/^+^, Sorcs2^-/-^*^58^, all backcrossed over nine generations to the C57BL/6j BomTac background. *Ror2^+/-^; Sorcs2^+/-^* double heterozygotes were generated by crossing *Sorcs2^+/-^* with *Ror2^+/-^* line.

Embryos were obtained from time-mated (6pm – 6am), plug-positive breeding females euthanized by cervical dislocation. The abdominal cavity was opened by midline incision, and embryos were collected by opening the uterine sacs and removing extra-embryonic membranes. Embryos older than E11.5 were euthanized by decapitation. Embryos were washed in ice-cold 1x PBS-pH7.4 for 2 min prior to tissue collection. Embryos were analyzed at developmental stages E10.5–E15.5. Embryonic sex was not determined, and the analyses were performed irrespective of sex. Sex was determined for 9–10-week-old mice and considered in the analyses (**Suppl. Fig. S13b-c**). Sex-disaggregated data are provided in the Source Data file.

### *Xenopus laevis* husbandry and microinjections

Animal care and research protocols followed institutional guidelines (Etiska Nämnden, permit N241/14). *Xenopus laevis* eggs were obtained by injecting frogs with 700 units of human chorionic gonadotropin (Pregnyl, Merck Sharp & Dohme). Fertilization was achieved using a sperm suspension, and embryos were dejellied with 1% thioglycolic acid at the two-cell stage. Embryos were cultured in 0.2× Marc’s Modified Ringer’s (MMR) solution at 18–21°C and staged as described^95^.

#### Cloning and mRNA synthesis

Human *SORCS2*^42^ was amplified with Phusion® DNA Polymerase (NEB), and cloned into the EcoRI/XbaI sites of the pCS2-super vector^63^ using forward primer AGCTGAATTCAAGCTTCTGCCTGCCGCCTG, and reverse primer AGCTTCTAGATCAGCTCACCAGGTAGCTGT. Mouse *Ror2*^96^ was cloned into the XhoI site of pCS2-super vector, and orientation was confirmed via restriction digestion. Constructs were linearized with NotI, purified (Qiagen PCR purification kit), and transcribed into capped mRNAs using mMessenger mMachine sp6 kit (Ambion).

#### Microinjections

For embryo elongation assay, 250-1000 pg of *Ror2* mRNA, with or without 1000 pg of *SORCS2* mRNA, was injected into two dorsal marginal cells at the 4-8-cell stage embryos. For the animal cap assay, 1000 pg of *Ror2* mRNA, with or without 2000 pg of *SORCS2* mRNA, was injected into the animal pole at the 2-cell stage embryos. The embryos’ phenotypes were analyzed at stage 32. Animal caps (ectodermal region) were dissected at stage 9, treated *ex vivo* with 20 ng/ml of recombinant human Activin-A (R&D Systems) in 0.1% BSA-0.4x MMR (Marc’s Modified Ringer’s solution) for 1 hr at 20-22°C, washed in 0.2x MMR, and cultured in 0.1% BSA-0.2x MMR until sibling embryos reached stage 17. We analyzed 14-21 embryos/group in the embryo elongation assay and 6-7 embryos/group in the animal cap assay.

### Zebrafish husbandry and microinjections

Zebrafish were maintained at 28.5 °C under a 14 h light/10 h dark cycle and fed four times daily, following national guidelines (permits 2017-15-0202-00098, 2022-15-0202-00169). Embryos were obtained from natural crosses of pathogen-free adults, monitored with the Zebrafish Comprehensive PCR Panel (Idexx GmbH). The *sorcs2^sa116^*^42^ mutant allele, a nonsense mutation in exon 10 of the *sorcs2* gene, was sourced from the Zebrafish Mutation Project^97^ and used as the *sorcs2 KO* line as described previously^58^, with genotypes confirmed by DNA sequencing. The AB strain (European Zebrafish Resource Center) provided inbred *WT* embryos, while ABxCasper strain served as an outbred *WT* line. Embryos were rinsed in E3 buffer (5 mM NaCl, 0.17 mM KCl, 0.33 mM CaCl_2_, 0.33 mM MgSO_4_, 2 mM Hepes-pH7.3, 10^-5^ % methylene blue) and kept in 10 cm petri dishes at 28°C incubator. Following developmental stages were analyzed: 3.5 hpf, 50% epiboly, bud stage, 7-somite stage, 22-somite stage, and Prim-6.

#### Microinjections

Morpholinos (MOs) were obtained from GeneTools, LLC, and were previously validated^58, 67, 98, 99^. These MOs were used:

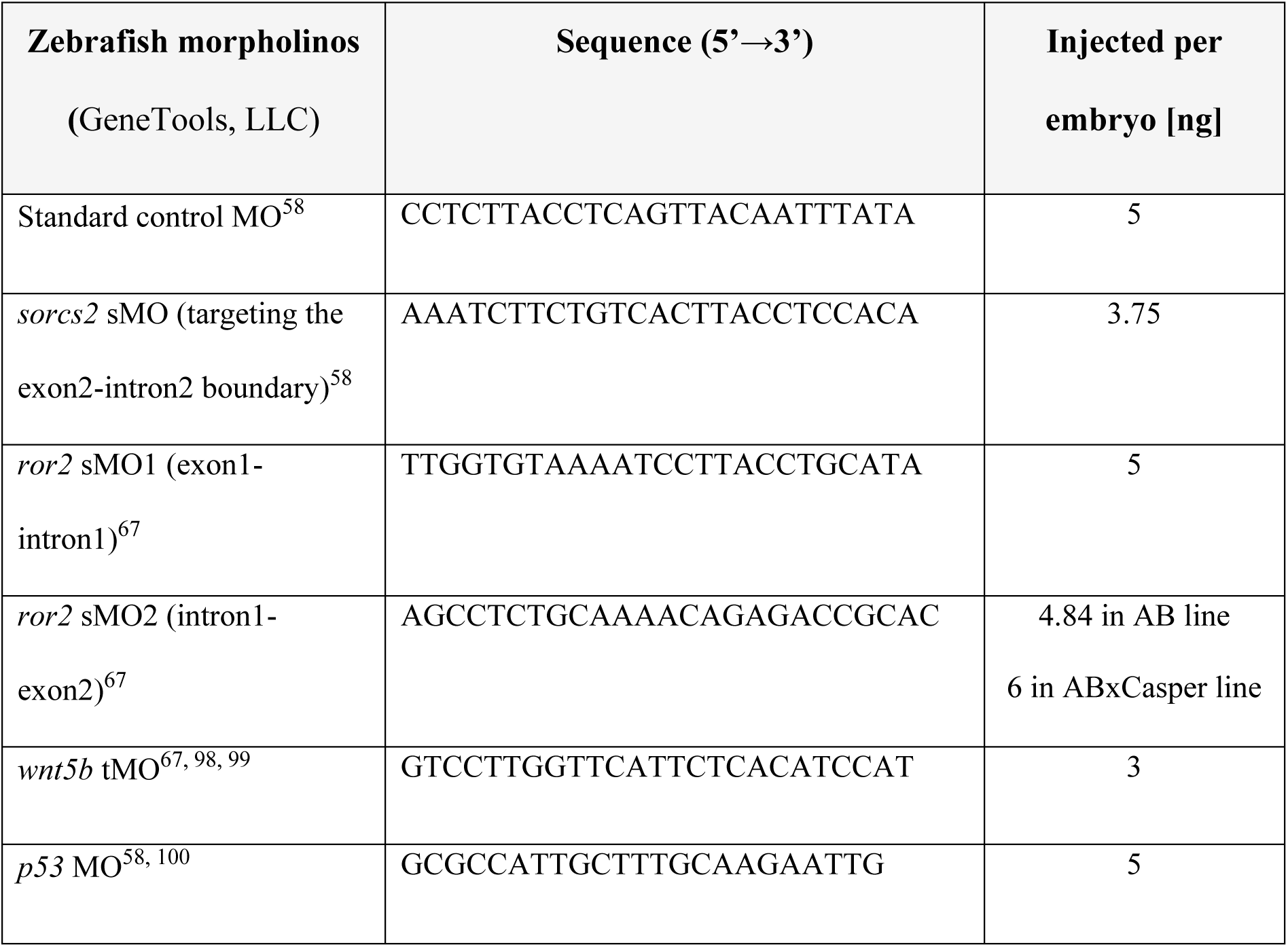

Material for mRNA injections was prepared as described above for the *Xenopus laevis* model. Injection mixes contained 0.2% phenol red in 1 mM Hepes-pH7.5, and 5 nl was injected per embryo at the 1–4-cell stage using a pico-injector. Embryos were sorted 3 h post-injection and staged as described^101^. Embryos were anesthetized in Tricaine-E3 buffer and mounted in 3% methylcellulose for imaging. Bright-field images were captured with an Olympus IX71 microscope and DP71 camera. Ror2-dependent CE defects were assessed as anterior-posterior body axis truncation, shortened and widened yolk extension, and reduced head/brain development based on published literature^67^, classified according to observed severity. The number of analyzed embryos and independent microinjections is specified in the figures and figure legends.

### Cell culture and transient transfection

All cell lines were maintained at 37°C with 95% H_2_O and 5% O_2_. Cells were passaged with 0.05% Trypsin-EDTA (Thermo Fisher Scientific). Human embryonic kidneys 293T cells (HEK293T; ATCC, #CRL-3216) and mouse embryonic fibroblasts (MEFs) were cultured in DMEM supplemented with 10% FBS, 2 mM L-glutamine, 50 U/ml penicillin-streptomycin (Thermo Fisher Scientific). Mouse substantia nigra dopaminergic progenitor cells (SN4741; gift from Jin H. Son^53^) was cultured in the same medium with 0.6% glucose. To generate double-knockout *Ror1/2* cell line, MEFs with double-floxed *Ror1; Ror2* (gift from Henry Ho^10^) were transfected with Cre-recombinase and treated first with 0.25 μM of 4-hydroxytamoxifen (Sigma Aldrich), and then 0.1 μM of 4-hydroxytamoxifen on subsequent days^10^. The number of independent biological experiments is specified in the figures and figure legends.

#### Treatments

Cells were treated in complete media at least 24 h post-seeding. Mock controls were treated with DMSO or PBS-0.1% BSA. The following treatments were used: chloroquine (50 µM, 10 h post-transfection for 24 hours; Sigma Aldrich), dynasore (50 µm, 10 h post-transfection for 24 hours; Sigma Aldrich), recombinant Wnt3a (100 ng/ml, 2-17 h; R&D Systems, #1324-WN/CF), recombinant Wnt5a (100 ng/ml, 2-17 h; R&D Systems, #645-WN/CF), CK1ɛ/δ inhibitor PF670462 (5 µM, 17 h; Sigma Aldrich), and porcupine inhibitor LGK-974 (1 µM, 17 h; Sigma Aldrich).

#### Transient transfection

HEK293T cells were seeded in 40% confluence in complete medium 24 h prior to transfection. For co-immunoprecipitation, cells were seeded on 10 cm plates. For other applications, cells were seeded into 24-well plates at 40,000 cells/well with or without 13 mm glass coverslips coated with 0.1% gelatin or 1 mg/ml Poly-D-lysine (Sigma Aldrich). Transfections were performed with Lipofectamine-2000 (Thermo Fisher Scientific) or FuGENE 6 (Promega) following the manufacturers’ instructions, as 1 μg DNA + 3 μl transfection reagent, mixed in OptiMEM medium. DNA amounts used: 5 µg total for 10 cm plates (2.5 µg/plasmid) and 0.7 µg total for 24-well plates (0.2–0.3 µg/plasmid or 50 ng Renilla). Co-cultures were seeded on 10 cm plates, transfected, and re-seeded after 24 h into co-culture or individual conditions for imaging or functional assays. Cells were harvested 48–72 h post-transfection.

#### Plasmids

pcDNA3.1 empty vector, pcDNA3.1-Ror2-FL-Flag, pcDNA3.1-Ror2-ΔCRD-Flag, pcDNA3.1-Ror2-Δ745-Flag, pcDNA3.1-Ror2-Δ469-Flag (gifts from Sigmar Stricker^57^). Lrp4-GFP (gift from Ruth Herbst). pcDNA3.1-hSorCS2-FL, pcDNA3.1-hSorCS2-RKR (single-chain), pcDNA3.1- hSorCS2-T71027R (two-chain), pcDNA3.1- hSorCS2-Tailless plasmids were described previously^42^. Dvl2-GFP (gift from Robert Lefkowitz), Chimeric Ror2-ECD-Lrp6-ICD (gift from Akira Kikuchi^66^), pDS-Rab7-WT (Addgene), EGFP-C1-Rab5a (gift from Philip D. Stahl), GFP-Rab11-WT (Addgene), GFP-Rab4 WT (gift from Casper Hoogenraad), pLAMP1-mCherry (Addgene), pcDNA3.1-Vangl2-HA (gift from Thomas Ringstedt), pCS2-hPtk7-FL-Myc (gift from Annette Borchers), pcDNA3.2-Wnt5a-V5 (Wnt Open Source Kit, Addgene #1000000022) pcDNA3.1-Ror1-V5-His^102^, Super8X-TOPFlash_Luc (gift from Randy Moon), pRenilla_tkLuc (Promega).

### CRISPR/Cas9-derived knockout cell lines using lentivirus delivery

#### sgRNA design and cloning

Guide sequences were designed for a control line (irrelevant gene *EGFP*^63^, 5’-GAAGTTCGAGGGCGACACCC-3’), *Sorcs2 KO* (5’-GTAGGTTGTCCCAAAATCTG-3’) targeting exon 3, and *Ror2 KO* (5’-GTACACCAGCACATTGCGTG-3’) targeting exon 9. Guide sequences were inserted into lentiCRISPRv2_puro and/or lentiCRISPRv2_blast vectors by Golden-Gate cloning using BsmBII^103^ and verified via Sanger sequencing using a U6 primer (5’-ACTATCATATGCTTACCGTAAC-3’). LentiCRISPRv2_puro and lentiCRISPRv2_blast were a gift from Brett Stringer (Addgene plasmids #98290, http://n2t.net/addgene:98290, RRID:Addgene_98290; and #98293, http://n2t.net/addgene:98293, RRID:Addgene_98293). *EGFP-* and *Sorcs2*-sgRNAs were cloned into lentiCRISPRv2_puro, while the *EGFP-* and *Ror2* sgRNAs were cloned into lentiCRISPRv2_blast. All plasmids were purified using an endotoxin-free Zymopure Maxiprep kit (Zymo research).

#### Lentiviral production

Replication-deficient lentiviruses were generated using 3^rd^-generation packaging vectors (pMD2.G, psPAX/2) in HEK293FT cells, following local ethical guidelines (Arbetsmiljöverket, permit Dnr 2-5164/2016). HEK293FT cells (80% confluence) were transfected with 10 µg lentiCRISPRv2, 5 µg pMD2.G, and 7.5 µg psPAX/2 using Lipofectamine-3000 (1 µg DNA : 2 µl) in OptiMEM (Invitrogene). After 5 h, the transfection medium was replaced with DMEM containing 10% FBS and 1% Pen/Strep. Viral supernatants were collected at 48 and 72 h, pooled, centrifuged (200 g, 5 min, RT), filtered (0.45 µm), and ultracentrifuged (60.000 g, 2 h, 4 °C). Viral pellets were re-suspended in 200 µl PBS, incubated overnight at 4 °C, aliquoted, and stored at -80 °C.

#### Generation of monoclonal CRISPR/Cas9 cell lines

SN4741 cells were seeded at 40% confluence in 6-wells plates 24 h before transduction. Cells were transduced overnight in 1.25 ml fresh complete medium with 8 μg/ml of polybrene (Millipore) and 25 μl of lentiviruses per well. Mock wells (no virus) served as controls for antibiotic selection. We first generated *WT* lines and *Sorcs2 KO* lines, transducing *sgEGFP-* lentiCRISPRv2_puro or *sgSorcs2*-lentiCRISPRv2_puro, respectively. Medium was exchanged 24 h post-transduction, puromycin selection was initiated at 48 h post-transduction (4 μg/ml; ThemoFisher Scientific) and continued for 9 days. Mock controls died within 48 h. Positive clones were isolated via serial dilution and confirmed by immunoblotting. To generate *Ror2 KO* and *Ror2; Sorcs2 double KO* lines, *sgEGFP*- lentiCRISPRv2_blast or *sgRor2-*lentiCRISPRv2_blast were transduced into *WT* or *Sorcs2 KO* lines as described above, followed by blasticidin selection 48 h post-transduction (20 µg/ml, increased to 30 µg/ml next day; InvivoGen). Cells were maintained under selection until mock cells died. Positive clones were isolated, expanded, and confirmed for protein absence by immunoblotting.

### Co-immunoprecipitation and immunoblotting

#### Co-immunoprecipitation

For mass spectrometry (MS/MS) analysis, endogenous Ror2 was pulled down from SN4741 cells grown to 90% confluency on 15 cm plates. For small-scale co-IP, SN4741, MEFs, or transiently transfected HEK293T cells were grown on 10 cm plates. Cells were harvested on ice, washed twice with ice-cold PBS (pH 7.4), and lysed in a mild lysis buffer (150 mM NaCl, 20 mM Hepes pH 7.4, 0.08 mM EDTA, 1% Triton X-100, 0.1 mM DTT, protease inhibitor cocktail [Roche], and phosphatase inhibitors [CalBiochem]). Lysates were prepared as described previously^63^ and incubated overnight at +4°C with following antibodies: rabbit Ror2 (HPA021868, Sigma Aldrich), rabbit Ror2 (gift from Henry Ho^10^), FlagM2 mouse (Sigma Aldrich), goat GFP (Abcam), and mouse V5 (Invitrogene). Normal rabbit IgG (Santa Cruz) served as a negative control (**Suppl. Table S7**). For MS/MS, 5-10 µg/condition of antibody was used; for small-scale co-IP, 1 µg/condition of antibody was used. Protein-G magnetic beads (Invitrogen) were added to the lysate the next day for 2-4 hours. Beads were washed five times with lysis buffer (without DTT). Protein complexes were eluted with 1× Laemmli buffer, denatured at 95 °C for 5 min, and loaded onto 8-10% polyacrylamide SDS-PAGE gels.

#### Immunoblotting

Immunoblotting was performed as described previously^58, 63^. After the SDS-page, proteins were transferred onto PVDF membrane (GE-Healthcare) using wet-transfer on ice. Membranes were blocked with 5% milk for 45 minutes at RT and incubated with primary antibodies overnight at 4 °C (**Suppl. Table S7**). Loading controls were always analyzed on the same membranes as the corresponding proteins of interest. Where applicable, membranes were cut at the appropriate molecular-weight positions, as determined using the protein ladder, to allow incubation of individual membrane sections with different primary antibodies. In instances where the amount of protein loaded per well was low, membranes were not stripped between antibody incubations but were sequentially probed with additional primary antibodies. Sequential probing was performed only when the respective target proteins could be clearly distinguished based on their molecular weights and when antibody detection yielded clean background signal, allowing unambiguous identification of the individual target proteins. Consequently, bands corresponding to previously detected proteins may remain visible on some of the original uncropped immunoblots. After washing, membranes were incubated with HRP-conjugated secondary antibodies (1:5000, Sigma-Aldrich or 1:2000, DAKO) at room temperature for 1 h. The immunoblots were developed with Amersham ECL Prime system (GE Healthcare) or Immobilon Forte Western HRP substrate (Sigma Aldrich) and imaged on charge-coupled device camera (Bio-Rad), LAS4000 Imager (FujiFilm), and iBright Imager (Thermo Fisher Scientific). Unsaturated immunoblots were analyzed for band intensity using peak area (square pixel = 1) or mean gray value with subtracted background and inverted pixel density (final score = net value) in Fiji software^104^. The number of independent biological experiments is specified in the figures and figure legends. Original uncropped immunoblots corresponding to the main figures are provided in the **Source Data file**, while original uncropped immunoblots corresponding to the Supplementary Figures are provided in **Supplementary Figs. S19-S24**.

### Subcellular fractionation of cytosolic and plasma membrane proteins

#### Samples

SN4741-GFP/Blasticidin control cells (clone 6) and SN4741-SorCS2 KO cells (clone 3) were seeded 48 h prior to lysis and harvested at near confluence. For each biological replicate, cells from 3-4 pooled 10 cm dishes were used.

#### Subcellular fractionation

Plasma membrane and cytosolic protein fractions were isolated using the Plasma Membrane Protein Extraction Kit (Abcam, #ab65400) according to the manufacturer’s recommendations. All steps were performed on ice or at 4 °C. Cells were washed twice with ice-cold PBS, scraped, and pelleted by centrifugation at 3,000 rpm (700g) for 5 min at room temperature. Pellets were resuspended in 2 mL of ice-cold homogenization buffer supplemented with protease inhibitors and lysed by mechanical homogenization on ice using a precooled hand-held homogenizer (20 x 2s strokes, 5 min incubation on ice, repeated 5 times). Homogenates were clarified by two sequential centrifugations at 700 × g for 10 min at 4 °C. Protein concentration was determined using a BCA assay and evened for all samples. Total protein input ranged from 2000 µg to 11000 µg. Cytosolic and membrane fractions were separated by two sequential centrifugations at 10,000 × g for 30 min at 4 °C, yielding a cytosolic supernatant and a membrane-enriched pellet. The pellet was subjected to aqueous two-phase separation using the upper and lower phase solutions supplied with the kit. Phase separation was performed by centrifugation at 1,000 × g for 5 min at 4 °C, and the plasma membrane–enriched upper phase was recovered according to the protocol. Plasma membrane proteins were diluted in ice-cold MQ water, incubated for 5 min on ice, and pelleted by two sequential high-speed centrifugations at 21,320 × g for 30 min at 4 °C. The final plasma membrane pellet was resuspended in 50µL RIPA buffer and prepared for immunoblotting by addition of 4x loading buffer (Invitrogene) supplemented with final 20 mM DTT, similarly to 50 µL aliquots from homogenate and cytosolic fractions. Samples were vortexed, briefly centrifuged, and stored at -70 °C until SDS-page electrophoresis and immunoblotting. Analyzed data comprises five independent experiments, with n = 4-14 biological replicates/group. Each fraction was normalized to GAPDH and *Ctr* samples

### Analysis of Ror2 and Dvl phosphorylation by immunoblotting

#### Dvl phosphorylation

Phosphorylated and dephosphorylated Dvl2/3 bands were quantified in Fiji using a fixed rectangular region of interest (ROI) and mean gray value measurement. The same ROI was applied across all samples to measure pixel intensity of phosphorylated Dvl (p-Dvl2/3), dephosphorylated Dvl (depho-Dvl2/3), and background signal immediately below each band. GAPDH was quantified from the corresponding blots as loading control. Measured pixel intensities were inverted using the formula “255 – X”, where 255 represents the maximum pixel intensity of an 8-bit image. Inverted background values were subtracted from the inverted band values to obtain net (NET) intensities. Dvl band intensities were normalized to GAPDH NET intensity. Total Dvl2/3 was calculated as the sum of NET intensities of p-Dvl and deph-Dvl normalized to GAPDH. Dvl phosphorylation levels were expressed as the percentage of phosphorylated Dvl relative to total Dvl (total Dvl = 100%). Values of dephosphorylated Dvl were not shown and correspond to the remaining percentage (100% - pDvl), except in **Fig. 8e**, where values are presented as the ratio pDvl3/deph-Dvl3, as the plasma membrane fraction did not contain absolute Dvl levels. The number of independent biological experiments is specified in the figures and figure legends.

#### Ror2 phosphorylation

Ror2 phosphorylation was analyzed in Fiji using two independent antibodies for immunoblotting: “Ror2 Sigma” (#HPA021868) and “Ror2 R&D” (#AF20264SP). The Ror2 Sigma antibody detects Wnt5a-dependent Ror2 phosphorylation as a shift in electrophoretic mobility^76, 77^ (see also **Suppl. Fig. S18h**), quantified as distance of the Ror2 band from a reference line drawn at 75 kDa. Relative band migration was normalized to 100% in *WT/control* samples. In contrast, the Ror2 R&D antibody detects Wnt5a-dependent Ror2 phosphorylation as a distinct upper band^85^. For this antibody, phosphorylated (p-Ror2) and dephosphorylated (deph-Ror2) bands were quantified as described above for Dvl analysis, including background subtraction and normalization. The number of independent biological experiments is specified in the figures and figure legends.

### Immunoprecipitation-coupled liquid chromatography tandem mass spectrometry

#### Experimental design and strategy

To characterize the Ror2 interactome, we performed Ror2 immunoprecipitation (IP) followed by tandem mass spectrometry (MS/MS) using mouse dopaminergic progenitor-like cell line SN4741 and an antibody recognizing the Ror2 C-terminus (Ror2_Sigma). We performed four independent IP-MS/MS experiments comprising in total 10 samples: IgG control (n = 4), IPRor2_vehicle (n = 4), and IPRor2_+2h Wnt5a (n = 2), with each sample analyzed in two separate MS/MS runs based on the in-gel saturation; run 1 = unsaturated proteins, run 2 = saturated proteins (visualized in **Fig. 1b**). The results from both runs were combined. Normal IgG pulldown served as a negative control. In two experiments, cells were treated with recombinant Wnt5a for 2 h to identify Wnt5a-responsive interactors. The raw data are available via the ProteomeXchange Consortium under accession PXD085067 and can be accessed through the PRIDE repository: https://www.ebi.ac.uk/pride/archive/projects/PXD085067. Raw files corresponding to individual independent experiments can be identified by the following filename prefixes: Exp. 1, “20120817_02-05_Q1_”; Exp. 2, “20130125_02-05_Q1_”; Exp. 3, “20130918_22-27_Q3_”; and Exp. 5, “20140516_14-19_F_”. Proteins detected in IgG controls were subtracted as nonspecific background, and proteins identified in at least two independent biological replicates were retained as candidate Ror2 interactors. The experimental design and individual samples are detailed in **Fig. 1c** and **Suppl. Figs. S1–S3**. The reproducibility of the approach was further supported by the recurrent identification of previously established Ror2-interacting proteins (**Suppl. Table S4**).

#### Co-immunoprecipitation

Endogenous Ror2 was pulled down from SN4741 cells grown to 90% confluency on 15 cm plates. Cells were harvested on ice, washed twice with ice-cold PBS (pH 7.4), and lysed in a 3mL/dish mild lysis buffer for 20 min (150 mM NaCl, 20 mM Hepes pH 7.4, 0.08 mM EDTA, 1% Triton X-100, freshly added 0.1 mM DTT, protease inhibitor cocktail [Roche], and phosphatase inhibitors [CalBiochem]). The lysates were centrifuged at 18.000g, +4°C for 20 min and the supernatant was subsequently transferred to a new tube. The lysates were pre-cleared using 30 uL of washed protein G-magnetic beads per sample, while rotating in +4°C for 45 min. Lysates were incubated with 5-10 µg antibody/sample for 3h at +4°C: rabbit Ror2 (HPA021868, Sigma Aldrich) and normal rabbit IgG (sc-2027, Santa Cruz). Protein-G magnetic beads (Invitrogen) were added to the lysate and incubated overnight at +4°C. Beads were washed six times with lysis buffer (without DTT). Protein complexes were eluted with 30 uL of 1× Laemmli buffer, denatured at 95 °C for 5 min, and loaded onto 8% polyacrylamide SDS-PAGE gels.

#### LC-MS/MS

The samples were freshly prepared and directly loaded onto 8% polyacrylamide SDS-PAGE gels, which were subsequently fixed using 100 % methanol and silver-stained^63^. Each gel lane was divided into 20 bands, and de-stained in 50 mM ammonium bicarbonate and 50% acetonitrile. Proteins were reduced in 10 mM DTT, alkylated in 55 mM iodoacetamide, and dehydrated in 100% acetonitrile, using a liquid-handling robot (MultiProbe II, Perkin Elmer). Gel pieces were dehydrated in 100% acetonitrile, followed by tryptic digestion overnight using 13 ng/µl trypsin at 37 °C. Extracted peptides from consecutive bands were pooled into two MS/MS runs per lane based on protein saturation; first MS/MS run comprised unsaturated proteins within the specific silver stained gel lane, and second MS/MS run comprised saturated proteins within the silver stained gel lane (**Fig. 1b**). Peptides were analyzed as described previously^63^. Briefly, nano-LC-MS/MS analysis was carried out using an Easy-nLC system (Thermo Scientific) directly coupled to an Orbitrap Q Exactive mass spectrometer (Thermo Scientific). Peptide separation was performed on a 10-cm fused SilicaTip column (New Objective, Inc.) packed in-house with 3-µm C18-AQ ReproSil-Pur (Dr. Maisch GmbH). Peptides were eluted using a linear gradient from 3% to 48% acetonitrile over 89 min at a flow rate of 300 nl/min. MS acquisition consisted of a full survey scan covering m/z 300–1650, acquired at a resolution of R = 70,000 at m/z 400, followed by up to ten data-dependent HCD MS/MS scans targeting the ten most abundant precursor ions with charge states ≥2, acquired at a resolution of R = 17,500.Tandem mass spectra were extracted using Raw2MGF (in-house software, Karolinska Institutet) and searched against the mouse SwissProt protein database (version 2012.03) using Mascot Daemon 2.3.02 (Matrix Science Ltd.). Search parameters included trypsin cleavage (allowing two missed sites), a peptide mass tolerance of 10 ppm, and a fragment ion tolerance of 0.05 Da. Fixed modifications included carbamidomethylation of cysteine, while variable modifications included methionine oxidation and asparagine/glutamine deamidation. Proteins were identified with a Mascot score threshold of 20, a significance threshold of 0.02, and MudPit scoring. Data were filtered to include only proteins meeting these criteria, with results exported for further analysis.

#### Subsequent data analysis

Specific Ror2 binding partners were identified by comparison to IgG controls (**Suppl. Table S1**). IP-specific hits were generated by subtracting proteins also found in the IgG controls (“-IgG”) (**Suppl. Table S2**). A total of 625 recurrent Ror2 binding partners were detected in at least two replicates (**Suppl. Table S3**). Protein IDs were converted to UniProt identifiers using PANTHER database (version 19.0)^105^ and analyzed via DAVID^106^ for gene ontology (GO) and KEGG pathway enrichment, with 58% – 93% of proteins annotated in DAVID. Only GO terms with P value < 0.05 were considered significant (**Suppl. Tables S5-6**).

### Ubiquitination assay

HEK293T cells were seeded into 6-well plate 600,000 cells/well and transfected with 100–200 ng plasmid per well using polyethylenimine (2 mg/ml, 2:1 µg DNA ratio). Medium was replaced 5 h post-transfection. 24 h post-transfection, cells were pretreated with epoxomicin (100 nM, Sigma-Aldrich) for 8 h and lysed in 600 µl buffer (6 M guanidine hydrochloride, 0.1 M NaH_2_PO4/Na_2_HPO_4_ pH 8.0, 10 mM imidazole). Lysate were sonicated, heated (95 ◦C, 6 min), and centrifuged (16.200 g, 20 min). 500 µl of cleared lysate was incubated overnight with 5 µl magnetic His-beads (GE-Healthcare). Beads were incubated with washing buffer (8 M urea, 0.1 M NaH_2_PO_4_/Na_2_HPO_4_ pH 6.3, 0.01 M Tris pH 8.0, 20 mM imidazole), re-suspended in 100 µl of 1× sample buffer, and denatured (95°C, 6 min) before loading into the 8% polyacrylamide SDS-page gels. Input lysates were prepared by ethanol precipitation of 100 µl lysate in 900 µl ice-cold ethanol overnight at -20 °C, followed by centrifugation (16.200 g, 20 min), washing with 90% ice-cold ethanol, drying, and resuspension of the pellet in 100 µl 1× sample buffer^60^. The assay was performed in two independent experiments with similar results.

### Dual luciferase reporter assay

HEK293T cells were seeded on 24-well plates and transfected with 50 ng Renilla-tkLuc, 200 ng Super8xTopFlash-Luc, 200 ng Ror2_ECD-Lrp6_ICD, 200 ng hSorCS2/pcDNA3.1./Ror2-FL-Flag. Cells were transfected at least 24 h post-seeding and treated in complete media at least 24 h post-transfection. Mock condition corresponds to cells treated with PBS-0.1% BSA for 16h, while Wnt treatment comprises incubation with recombinant Wnt3a (100 ng/ml, 16 h; R&D Systems, #1324-WN/CF) and recombinant Wnt5a (100 ng/ml, 16 h; R&D Systems, #645-WN/CF). Cells were lyzed 48 hours post transfection in 50 µl lysis buffer. Samples were analyzed with dual luciferase reporter assay kit (Promega) according to manufacturer’s instructions as described previously^63^. The samples were measured for 1 second per channel using a Victor plate-reader (PerkinElmer). LARII luciferase activity (Super8xTopFlash signal) was normalized to StopGLO luciferase (Renilla signal) to account for overall translational efficiency and further normalized to 1 (pcDNA3.1, mock) to assess fold change. The assay was performed in four (**Fig. 4g**) or three (**Fig. 8a**) independent biological experiments.

### Immunofluorescence and proximity ligation assay

Transfected HEK293T cells were fixed 24 - 72 hours post transfection in 4% PFA at RT for 20 min, and subsequently washed in 1x PBS-pH7.4. The samples were blocked in 1x PBS-pH7.4 with 0.1% BSA, 0.3% Triton-X-100 and 5% donkey serum (Jackson Immuno Research) at RT for 45 min, and incubated with primary antibodies in a wet chamber at 4 °C overnight (**Suppl. Table S7**). Coverslips were washed in 1x PBS-pH7.4, and incubated with secondary antibodies Alexa Fluor-488/-568/-647 (1:800, Thermo Fisher Scientific) and Hoechst (1:2000, Thermo Fisher Scientific) at RT for 1 h. Coverslips were washed in 1x PBS-Ph7.4 and then mounted with Mounting Fluorescence Medium (DAKO). For the PLA assay, we used commercially available kit Duolink *In Situ* Detection Reagents Orange (Sigma Aldrich) and followed the manufacturer’s protocol. We used anti-FlagM2 (1:1000, mouse, Sigma) and hSorCS2 (1:2000, rabbit, DAKO) antibodies for overnight incubation, and anti-mouse-PLUS and anti-rabbit-MINUS Duolink probes (Sigma Aldrich) prior ligation and amplification. Single transfections were incubated with both antibodies and both probes, serving as negative controls. All samples were imaged on confocal scanning microscope LSM-780 (Zeiss). The proximity ligation assay was performed in three independent biological experiments.

### Co-localization analysis of Ror2-SorCS2 complex in subcellular compartments

Co-localization of two channels was quantified using the EzColocalization plugin in Fiji software. Confocal images were split into individual channels and uploaded into EzColocalization as ‘Reporter 1’ and ‘Reporter 2’. Regions of interest (ROIs) of equal size were applied to all double-stained cells for cell identification. Co-localization was evaluated using the Threshold Overlap Score (TOS)^107^, employing Costes’ method for thresholding^108^. The TOS linear value, ranging from -1 (anti co-localization) to +1 (maximum co-localization), was used to quantify co-localization intensity^107^. True co-localization was defined as TOS > 0.1 and non-co-localization as TOS < 0.01. For quantitative analyses of overexpressed markers, 60 cells per experimental group were analyzed across three independent biological experiments (20 cells per group in each experiment). For endogenous markers, 16–60 cells per experimental group were analyzed across three independent biological experiments. Because the endogenous markers were detected using distinct antibodies with different staining characteristics and intensity distributions, direct statistical comparisons between the different markers were not considered appropriate (**Fig. 3l-m**).

### Whole-mount RNA *in situ* hybridization in zebrafish

#### Sample preparation

Zebrafish embryos were collected at specific stages (3.5 hpf, 50% epiboly, bud stage, 7-somite stage, 22-somite stage, and Prim-6)^101^. Embryos were anesthetized, de-chorionated, and fixated in 4% PFA-PBS at +4°C overnight.

#### Probes preparation and alkaline phosphatase WISH

Gene-specific mRNA sequences were obtained from NCBI database. Primers (amplifying products of 330-400 bp for cDNA and ∼1100 bp for gDNA) were designed using NCBI’s primer tool. OligoAnalyzer3.1 (Integrated DNA technologies) and BLAST analysis were used for quality control and probes specificity. T3 or T7 polymerase promotor sequences were added to reverse primers. Probe PCR fragments were amplified from cDNA with KOD HotStart polymerase (Novagen) following standard protocol using 250 ng of embryonic cDNA/gDNA and 5% DMSO. PCR products were purified using NucleoSpin PCR Clean-up (Macherey-Nagel) and transcribed with a DIG RNA labelling kit (Roche) using T3/T7 polymerase at 37°C for 2 h. Probes were purified with SigmaSpin Sequencing reaction CleanUp columns (Sigma Aldrich), and assessed for integrity via RNA gel electrophoresis (Flashgel System, Lonza). RNA concentration was measured on NanoDrop 2000 (Thermo Scientific). Gene-specific probes were validated with sequencing using the forward primer (Eurofins). For apWISH, embryos were processed as described previously^58^.

Embryos older than 22 hpf were bleached in 3% H2O2/0.5% KOH for 30 min. Probes (50 ng) were hybridized in 200 µL hybridization mix, and stained embryos were mounted in 75% glycerol and imaged on an Olympus IX71 microscope with a DP71 camera. We performed each staining across three independent biological experiments and assessed ≥10 embryos per group and experiment.

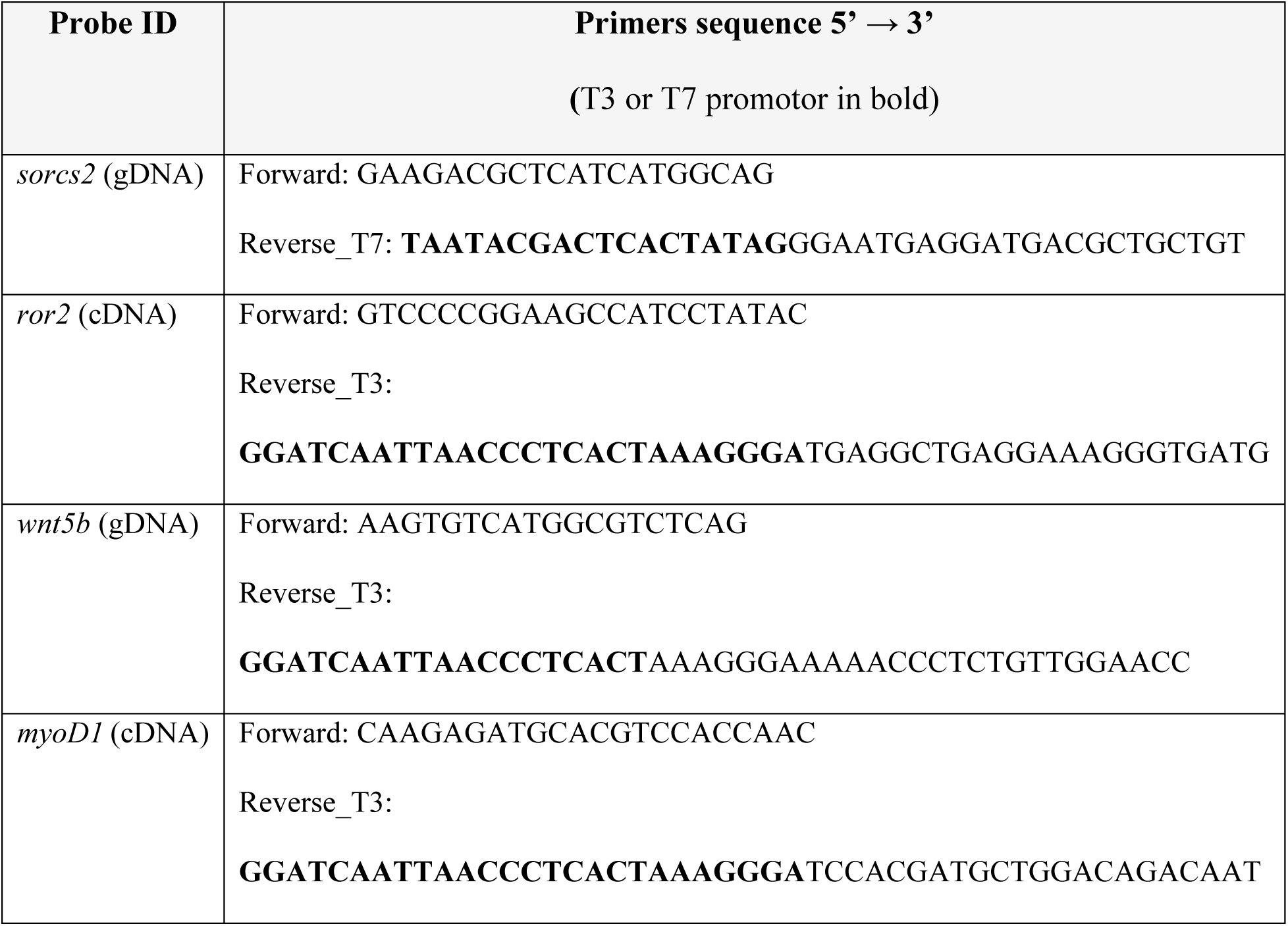

#### Fluorescent multiplex WISH

Whole-mount HCR v3.0^71^ (Molecular Instruments) was performed following the manufacturer’s protocol and as described previously^58^. Embryos were incubated overnight at 37°C with 5 pmol/μl of each probe. Amplifiers used were HCR-B1 for *wnt5b*-B1, HCR-B2 for *sorcs2*-B2, and HCR-B3 for *ror2*-B3 detection. To enhance the signal, two different *sorcs2*-B2 probes were combined. After washing, embryos were mounted laterally on coverslips using Prolong Glass Antifade Mountant (Thermo Fisher Scientific). Yolk sacks were removed for 22-somite and Prim-6 stage embryos. Imaging was performed with a Zeiss LSM-780 confocal laser-scanning microscope using a 20× objective with 0.6× zoom. Younger stages were captured as single planes and post-processed in Adobe Photoshop. Z-stacks were acquired at 4 µm intervals for the 22-somite stage and 5 µm intervals for the Prim-6 stage, with high-resolution snapshots exported from Imaris_3D-Viewer (Oxford Instruments) after mild post-processing. Brain morphogenesis was assessed with Fiji. We performed each staining across three independent biological experiments and assessed ≥10 embryos per group and experiment.

### Macro-anatomical morphometric analysis using brightfield microscopy

Live E12.5 and E14.5 mouse embryos were imaged in ice-cold 1× PBS (pH 7.4) using a Zeiss stereomicroscope in brightfield mode. Morphometric measurements were performed in Fiji using calibrated images and were applied consistently across embryos. At E12.5, measurements included dorsal brain length (from the anterior forebrain to the hindbrain-spinal cord interface), forebrain-midbrain thickness (from the anterior forebrain to dorsal midbrain), nasal-maxillary width (maximum width of nasal and maxillary regions), and trunk length (distance from the caudal end of the hindbrain to the base of the hindlimb). A total of 47 littermate embryos derived from 6 independent breeding females were quantified at E12.5. At E14.5, additional parameters reflecting advanced morphological development were quantified, including naso-maxillary length (anterior-posterior length of the nasal and maxillary regions), forelimb length (from the shoulder to the distal digit), and hindlimb length (from the hip to the distal digit). A total of 48 littermate embryos derived from 9 independent breeding females were quantified at E14.5.

### Light-sheet microscopy and Imaris-based segmentation of *Hb9:GFP/+* lines

We analyzed the E12.5 Hb9:GFP/+ (*WT* and *Sorcs2 KO*) samples and the corresponding light-sheet images described in^58^. Light-sheet images were post-processed in Zen Blue 3.7 software (Zeiss) and converted into “.ims” format using ImarisFile Converter 10.2.0 software (Oxford Instruments). the segmentation, isolated areas of interests and the measurement points are visualized in **Suppl. Video SV1**. The analysis comprised n = 10 littermates *Hb9:GFP/+; Sorcs2^+/+^* embryos derived from 3 independent breeding females and n = 9 littermate *Hb9:GFP/+; Sorcs2^−/−^*embryos derived from 2 independent breeding females.

#### Spinal cord segmentation

The images were segmented in Imaris v10.2.0. (Oxford Instruments) based on fluorescence and using Surfaces. 3D segmentation of the spinal cord was performed based on high-intensity Hb9:GFP fluorescence and histological landmarks, including head-trunk interface and hypoglossal nerve^109^. A region of interest (ROI) encompassing the spinal cord was defined manually in XYZT space, and segmentation was restricted to this ROI. Surfaces were generated from the Hb9:GFP channel using manual intensity thresholding (threshold value = 214.1), thereby selectively segmenting regions with high fluorescence intensity. Surface smoothing was enabled with a grain size of 50 µm. Segmented objects were filtered by size, retaining only surfaces exceeding 1.19 × 10⁸ voxels. The spinal cord length (applied from outer and inner side) and spinal cord angle were measured using measurement points.

#### Forelimb and hindlimb bud segmentation

Three-dimensional segmentation of forelimb buds and hindlimbs was performed in Imaris based on limb autofluorescence. For each sample, a region of interest (ROI) encompassing the respective limb was manually defined in XYZT space, and segmentation was restricted to this ROI. Surfaces were generated from the autofluorescence channel using manual intensity thresholding to selectively segment regions with high fluorescence intensity. For forelimb buds, a single manual threshold was applied (threshold value = 214.1), whereas hindlimb segmentation was performed using dual manual thresholds (threshold A = 336; threshold B = 1716) to account for stronger and more heterogeneous paw autofluorescence. Surface smoothing was enabled, with a grain size of 50 µm for forelimb buds and 100 µm for hindlimbs. Size-based filtering was applied to forelimb segmentations to exclude small objects, retaining only surfaces larger than 1.19 × 10⁸ voxels, while no size filtering was applied to hindlimb segmentations. Classification and shortest-distance calculations were enabled for downstream spatial analyses. We extracted quantitative measurements of segment areas and volumes from the Imaris Statistics module. The limb diameter (A-P, Pr-Di) were measured using measurement points.

### Fluorescent multiplex RNA *in situ* hybridization in VM sections

#### Sample preparation

Wild-type embryos were collected at E10.5 and E11.5 within 5 minutes after sacrificing the mother. Fresh-frozen (FF) embryos were embedded in Tissue-TEK O.C.T. (Sakura) on dry ice. For the consecutive IF staining, embryos were fixed in 4% PFA-PBS at +4°C for 2 h, washed in ice-cold PBS-pH7.4, immersed in 10%, 20%, and 30% saccharose solutions, and embedded in O.C.T. Midbrains were cryo-sectioned using Cryostar NX70 (Axlab) as 20 µm sections for FF and 16 µm for PFA-fixed samples. Coronal sections of midbrain were collected sequentially on SuperFrost Slides (Thermo Scientific), air-dried in - 20°C for 2h, and stored at -80°C. Tyrosine hydroxylase IF staining (PelFreeze) confirmed the mDa domain position.

#### RNA scope

We used 3-multiplex fluorescent RNA scope kit v1 (BioTechne), following the manufacturer’s protocol. Probes included *Mm-Sorcs2-C1* (#473411-C1), *Mm-Ror2-C3* (#430041-C3), *Mm-Lmx1a-C2* (#493131-C2), with AMP4-FL-B developing solution (BioTechne). Nuclei were counterstained with Hoechst for 10 min (1:2000, Thermo Fisher Scientific). Slides were mounted with 24 mm x 60 mm coverslips using fluorescence mounting medium (DAKO). For IF, slides were washed in PBS post-RNA scope, blocked with PBTA (0.1% BSA, 0.3% TritonX-100, 5% donkey serum in 1x PBS-pH7.4) at RT for 1 h, and incubated overnight at +4°C with Glast-Alexa-488 conjugated antibody in PBTA. Slides were washed in 1x PBS-pH7.4 and counterstained with Hoechst (1:2000, Thermo Fisher Scientific) and mounted as described above. Imaging was performed on a confocal scanning microscope LSM-780 (Zeiss). We performed each staining across three independent biological experiments and three biological replicates.

### Real-time quantitative PCR

Dissected mouse ventral midbrains were stored at −70 °C prior RNA extraction. Total RNA was extracted using TRIzol reagent (Invitrogen) and resuspended in RNase-free water (Invitrogen). RNA concentration was determined using a NanoDrop One spectrophotometer (Thermo Scientific). For cDNA synthesis, 1 µg of RNA was reverse-transcribed using the High-Capacity RNA-to-cDNA Kit (Applied Biosystems), and the resulting cDNA was diluted to a final concentration of 20 ng/µl with ultrapure water. TaqMan quantitative PCR was performed on a QuantStudio 7 Flex Real-Time PCR System (Applied Biosystems) using MicroAmp Optical 384-well plates sealed with MicroAmp Optical Adhesive Film (Applied Biosystems). Reactions were run where possible in technical triplicates in a 10 µl reaction volume using TaqMan Fast Advanced Master Mix (Applied Biosystems) with 60 ng of input cDNA per reaction. The following TaqMan Gene Expression Assays (Invitrogen) were used: *Ror2* (Mm00443470_m1), *Sorcs2* (Mm00473050_m1), *Wnt5a* (Mm00437347_m1), *Wnt1* (Mm01300555_g1), *Tcf4* (Mm00443210_m1), and *Gapdh* (Mm99999915_g1). Ct values were determined by automatic thresholding using QuantStudio Real-Time PCR Software (Applied Biosystems). Gene expression levels were normalized to *Gapdh* as a reference gene (generating ΔCt values). Relative gene expression was calculated using the 2^-ΔΔCt^ method^110^, normalized to the mean of all WT E11.5 samples. A total of n = 35 *WT* embryos (7 breeding females) and n = 31 *Sorcs2 KO* embryos (6 breeding females) were analyzed across 3 independent qPCR experiments.

### Immunofluorescence of mDA lineage

#### Sample preparation & cryosectioning

Embryos from *Ror2^+/-^; Sorcs2^+/-^*breeding were collected at E12.5 within 5 minutes of sacrificing the mother. Trunks were used for genotyping, while heads were fixed in 4% PFA-PBS at +4°C overnight, washed in PBS-pH7.4, and sequential immersed in 10%, 20%, and 30% saccharose (in 1x PBS-pH7.4) solutions. Tissue was embedded in Tissue-TEK O.C.T. (Sakura) on dry ice and cryo-sectioned as 14 µm coronal midbrain sections using a Cryostar NX70 (Axlab). Sections were collected sequentially on nine SuperFrost slides (Thermo Scientific), air-dried at RT for 1 h, and stored at -80°C.

#### Immunofluorescence

Slides were thawed for 20 min, washed in PBS-pH7.4, and subjected to antigen retrieval (DAKO) in preheated 1x solution at 95°C for 20 min using a steamer (Bosch). After washing, sections were blocked in PBTA solution (0.1% BSA, 0.3% TritonX-100, 5% donkey serum in 1x PBS-pH7.4) at RT for 1 h. Following primary antibodies were used: rabbit TH (1:500, PelFreeze), goat Nurr1 (1:80, R&D Systems), mouse Sox2 (1:300, R&D Systems), rabbit Glast (1:100, Nittobo), sheep mSorCS2 (1:100, R&D Systems), mouse EEA1 (1:100, Cell Signaling). The primary antibodies were diluted in PBTA with 5% donkey serum and incubated either for 24 hours at 4°C for cell fate tracing and co-localization studies or 48 hours at 4°C for the EEA1+ vesicle reconstructions (**Suppl. Table S7**). Slides were washed and incubated with Alexa-conjugated secondary antibodies (1:800) and Hoechst (1:2000) at RT for 1 h (Thermo Fisher Scientific). After washing, slides were mounted with fluorescence mounting medium (DAKO) and 24 × 60 mm coverslips. Imaging was performed on LSM-780 and LSM-800 confocal scanning microscopes (Zeiss).

### Cell fate quantification of mDA lineage

The samples were prepared and stained as described in the previous section. We analyzed four TH-positive VM sections per embryo. The mDa domain borders were identified histologically based on the most lateral Nurr1+ staining (white lines, **Fig. 7f**). The following populations were distinguished based on their immunoreactivity: mDa progenitors [Sox2+, Nurr1-, TH-], postmitotic mDa neuroblasts [Sox2-, Nurr1+, TH-], and postmitotic immature mDa neurons [Sox2-, Nurr1+, TH+]. Cell counting was performed using the Cell counter plugin in Fiji. A total of n = 35 littermate embryos originating from 10 breeding females were analyzed across independent experiments.

### Confocal 3D imaging and quantitative analysis of EEA1+ endosomes in the mDA lineage

#### Samples

Quantification included littermate embryos derived from three independent *Sorcs2^+/−^* breeding pairs, comprising n = 5 *Sorcs2^-/-^* embryos and n = 5 control embryos (*WT + Sorcs2^+/-^*).

#### Z-stack confocal scanning

The sections and immunofluorescent staining were prepared as described above. We acquired confocal images of EEA1+ endosomes and the mDA lineage using LSM-800 confocal microscope with a 40x air objective and 0.5x zoom (Zeiss). We collected z-stacks at 1 µm intervals spanning 13–23 single sections (from background to background) at a resolution of 3000 × 3000 pixels. We acquired tiled images with 10% overlap when necessary.

#### Postprocessing and segmentation

The images were first post-processed in licensed Zen Blue 3.7 software (Zeiss) using the “Fuse tiles” function with 10% overlap and best-fit alignment, which were subsequently converted to .ims file type using ImarisFile Converter 10.2.0 software (Oxford Instruments). EEA1+ vesicles were segmented with Imaris 10.2.0 (Oxford Instruments) via “spot segmentation”. We integrated Spots detection algorithm with a fixed parameter set applied across all images: EEA1 as the source channel (estimated XY diameter 0.5 µm; Z diameter 1.0 µm), background subtraction enabled, quality filter > 1.92, and region growing based on absolute intensity with a manual threshold of 6.88. After automated detection, we manually removed spots outside the mDA lineage and excluded false-positive vesicles originating from blood vessels (the excluded area was determined based on autofluorescence).

#### EEA1+ vesicle quantification

We extracted quantitative measurements from the Imaris Statistics module, including vesicle counts and volume-related parameters. For downstream analyses, we calculated total vesicle counts and combined vesicle volumes across all EEA1+ endosomes within defined mDA subclusters, proliferating progenitors and postmitotic precursors (see e.g. **Fig. 7f**) and per tissue section. We subsequently combined data from all 4 sections to overview the mDA lineage across the A-P axis. We statistically evaluated the single sections too, but the genotypes did not show any differences (data not shown).

### Allen Brain Atlas, E10.5 spatial expression map, and single-cell RNA sequencing datasets

RNA in situ hybridization images of *Tcf4* in E11.5 and E13.5 mouse embryos were accessed and downloaded from Allen Brain Atlas database: https://developingmouse.brain-map.org/. Single-cell RNA sequencing profiles of developing mouse ventral midbrain^75^ were accessed as snapshots from the Linnarsson Lab data platform (https://linnarssonlab.org/ventralmidbrain/), with the raw data available at NCBI-GEO database (GSE76381). The E10.5 imputed spatial expression map profiles were accessed through the developmental atlas at http://www.mousebrain.org; the underlying data are deposited under BioProject accession PRJNA637987. Imputed expression data were inferred from spatial mapping of clusters to the E10.5 embryo and were given only for genes expressed in those clusters as described in^55^.

### Graphical visualization

Images were mildly post-processed using Fiji or Adobe Photoshop 2025. Figures were arranged in Adobe Illustrator 2025. Schematics were created in Adobe Illustrator 2025 or with BioRender.com by the corresponding author Alena Salasova. Z-stacks of HCR v3.0 images, light-sheet microscopy, and 3D *in vivo* sorting were exported as high-resolution snapshots from the Imaris 10.2.0_3D viewer (Oxford Instruments).

### Statistics and reproducibility

Results are presented as mean ± standard error of the mean (S.E.M.). All graphs and statistical analyses were performed using GraphPad Prism 10 (Dotmatics). Prior to analysis, data were first assessed for outliers (ROUT, Q = 5 - 10%) and Gaussian and lognormal distributions using Shapiro-Wilk test and Kolmogorov-Smirnov test. A P-value < 0.05 was considered statistically significant, with specific P-values, statistical tests, sample sizes, and numbers of independent experiments detailed in the corresponding figures and figure legends.

Experiments were performed following methodological optimization, using at least three independent biological experiments and replicates together with technical replicates and appropriate positive and negative controls, as specified in the Legend and Methods sections. Sample sizes were based on established experimental practice and our previous experience with these experimental systems^58, 63, 111^ and were considered sufficient to assess reproducibility across independent biological replicates. No statistical methods were used to predetermine sample sizes. For embryonic analyses, individual embryos represented biological replicates and, where applicable, were obtained from different breeding females and independent matings to account for biological and developmental variability. Embryos were analyzed irrespective of sex. Experiments were independently repeated as specified in the corresponding figure legends and, where applicable, by different researchers and/or in different laboratories. Key findings were validated using complementary experimental approaches and experimental models. Experiments affected by technical failure or inadequate sample quality were repeated using additional biological and technical replicates.

## Data Availability

Source data are provided with this paper. The data generated in this study required to interpret and verify the findings are provided in the Supplementary Information, Supplementary Datasets, and Source Data File. The mass spectrometry raw data are available via the ProteomeXchange Consortium under accession PXD085067 and can be accessed through the PRIDE repository [https://www.ebi.ac.uk/pride/archive/projects/PXD085067]. Reused single-cell RNA-sequencing data^75^ presented in this study are available in the NCBI Gene Expression Omnibus database under accession code GSE76381 [https://www.ncbi.nlm.nih.gov/geo/query/acc.cgi?acc=GSE76381]. Reused E10.5 imputed spatial gene-expression data^55^ were accessed through the developmental atlas at http://www.mousebrain.org, with the underlying data deposited under BioProject accession PRJNA637987 [https://www.ncbi.nlm.nih.gov/bioproject/PRJNA637987/]. RNA in situ hybridization images of *Tcf4* in E11.5 and E13.5 mouse embryos were obtained from the Allen Developing Mouse Brain Atlas [https://developingmouse.brain-map.org/]. Unique reagents, raw data generated in this study, and additional information are available upon request from the corresponding authors, Alena Salasova and Anders Nykjær, subject, where applicable, to institutional requirements and a material transfer agreement.

## Acknowledgements

We acknowledge imaging core facilities at Biomedicum (BIC), Karolinska Institutet (Sweden), Faculty of Health, Aarhus University (Denmark), and CEITEC-CELLIM (Czech Republic, supported by MEYS CR, LM2023050 Czech-BioImaging) for providing access to the equipment and technical support. We thank Dorothea Rutishauser and Carina Palmberg for performing the LC-MS/MS experiments, and Massimiliano Gaetani for technical assistance at the Proteomics Facility led by Prof. Roman Zubarev (Karolinska Institutet). We thank core facility INFINiGENE for explorative experiments not included in this study. We thank Maria Undengaard Vendelbo for her outstanding care of our mouse lines, and Shanzheng Yang, Patrik Ernfors, Alessandra Nanni, Benedicte Vestergaard, Petra Kompanikova, and all EA and AN lab members for technical support and discussions.

## Author Contributions Statement

Study Design & Conceptualization: AS, AN, EA. Methodology: AS, LK, CY, KKS, ACN, OB, MVG, JL, SN, ERA. Investigation: AS, LK, KKS, SN, CY, OB. Validation: AS, LK, CY, SN, ACN. Formal Analysis: AS, LK, CY, EMT. Visualization: AS, LK, CY, OB. Resources: AN, EA, AS, VB, CO, ERA, JK. Writing – Original and Final Draft: AS. Writing – Review & Editing: AS, EA, AN. Supervision: AS, EA, VB, AN, ERA. Project Administration: AS, EA, AN. Funding Acquisition: AN, EA, AS, VB.

