## Supplementary Figures and Figure Legend for "SorCS2 controls Wnt signaling homeostasis during neurodevelopment by regulating Ror2 activity and trafficking"

### Supplementary Figure Legend

**Suppl. Fig. S1: Validation and GO analysis of IP-MS/MS results.** (a) Mouse mDa progenitor cell line SN4741 served as a suitable mDa progenitor model, expressing Ror2 but lacking Tyrosine Hydroxylase, a marker specific to mDa neurons. (b) Pulldown of Ror2 using older lots of the Sigma antibody followed by IB showed that Ror2 interacts with SorCS2, Ptk7, Lrp4 and CAMKII. SorCS2 two-chain (2ch) isoform showed stronger binding to Ror2 than the single-chain (1ch). (c) Schematic of SorCS2 maturation after protein synthesis. The first cleavage of SorCS2 single-chain by Furin likely takes place in the Trans-Golgi Network (TGN) and results in the propeptide removal. From there, SorCS2 single-chain is transported in sorting vesicles to the plasma membrane to be available for cell signaling. In parallel, SorCS2 single-chain can be further cleaved into the two-chain isoform that might exhibit distinct signaling and functional features. This cleavage takes place either within the sorting vesicles or at the plasma membrane. Created with BioRender. (d) Validation of SorCS2 antibody and CRISPR-Cas9-generated *Sorcs2* knockout (KO) cell lines, alongside ventral midbrain tissue from *Sorcs2*<sup>+/-</sup> E14.5 littermates. (e) Functional annotation of 625 recurrent Ror2 binding partners identified by IP-MS/MS using DAVID for gene ontology (GO) terms related to “Protein Modification Type.” We matched 92.6% of proteins annotated in DAVID for the analysis. Only GO terms with $P < 0.05$ are shown (f) KEGG pathway analysis of Ror2 binding partners, focusing on disease-associated pathways, with 58.7% of proteins annotated in DAVID. Only GO terms with $P < 0.05$ are shown. (e-f) Detailed P-values to be found in **Suppl. Tables S5-S6**. All experiments were performed in independent biological experiments and replicates, $n \geq 3$.

**Suppl. Fig. S2: Imputed spatial expression map of E10.5 embryos.** The images were resourced as spatial gene expression map from *mousebrain.org*<sup>1</sup>. Spatial expressions of *Wnt1* (a), *Wnt5a* (b), *Sorcs2* (c), and *Ror2* (d) are shown as imputed data derived from cluster-to-embryo mapping at E10.5 to allow direct comparison across genes and their expression overlaps. *Wnt1* and *Wnt5a* were also directly measured by HybISS. Imputed *Sorcs2* panel (c) shows high expression midbrain floorplate, spinal cord, and otocyst (a key role in inner ear development).

**Suppl. Fig. S3: Validation of SorCS2-dependent specificity in the regulation and localization of Ror2.** (a-b) Co-expression of Ror2-Flag with SorCS2-FL in HEK293T cells, followed by immunofluorescence and confocal scanning. Experiments were performed in independent biological experiments, n > 5. (a) Upon overexpression, Ror2 predominantly localizes in filopodia, while SorCS2 co-expression reduces the filopodia formation and Ror2 localization at the plasma membrane. (b) Ror2-SorCS2 co-localized in the intracellular vesicles (yellow arrows) and aggregate-like structures (arrowheads). (c) Immunoblotting (IB) of Lrp4-GFP pulldown. Overexpressed Lrp4 interacted with Ror2 in HEK293T cells but did not reduce Ror2 protein levels. Three independent biological experiments. (d) Overexpressed hSorCS1c in HEK293T cells did not reduce Ror2 protein levels, contrasting the Ror2 co-expression with SorCS2. Two independent biological experiments. (e-f) Immunofluorescence of transfected HEK293T cells followed by confocal scanning. Two independent biological experiments. (e) SorCS1c did not co-localize with Ror2. (f) SorCS2 co-localized with Dvl2-GFP to minimal extent, excluding the possibility of overloading the translational machinery with overexpression. The scale bar = 10 µm; blue channel = nuclear Hoechst staining.

**Suppl. Fig. S4: SorCS2 downregulates total protein levels of several Wnt/PCP-associated Ror2 binding partners.** (a-c, e) Overexpression of constructs in HEK293 followed by immunoblotting. Two independent biological experiments. SorCS2 co-expression reduces total protein levels of Ror1-V5 **(a)**, Ptk7-myc **(b)**, Vangl2-HA **(c)**, and Wnt5a-V5 **(e)**. **(e)** Overexpression of constructs in HEK293 followed by immunofluorescence. Co-expression of SorCS2 with Wnt5a-V5 changed localization of intracellular Wnt5a to aggregate-like structures, similarly to Ror2. **(e)** Co-immunoprecipitation shows that SorCS2 single-chain also binds Wnt5a and downregulates its protein levels, similarly to Ror2. **(d-e)** Three independent biological experiments.

**Suppl. Fig. S5: The extracellular cysteine-rich domain of Ror2 contributed to SorCS2 intracellular co-localization.** **(a-i)** Immunofluorescence followed by confocal scanning of overexpressed Ror2 Full-length (Ror2-Flag) and Ror2 truncated variants in HEK293T cells, in absence **(b-e)** or presence **(f-i)** of SorCS2-FL **(a)**. Only the deletion of the ΔCRD domain failed to form aggregate-like structures and exhibited minimal to no co-localization with SorCS2-FL **(g)**, unlike Ror2-FL **(f)**, Ror2-Δ745 (tyrosine protein kinase deletion) **(h)**, or Ror2-Δ469 (the entire intracellular sequence deletion) **(i)**. Scale bar = 10 μm, blue channel = nuclear Hoechst staining. Experiments were performed in five independent biological experiments and replicates.

**Suppl. Fig. S6: Ror2-SorCS2 complex co-localizes with overexpressed endosomal and lysosomal markers.** All panels show overexpression in HEK293T cells. **(a)** Initial IB quantification of HEK293T cells co-expressing Ror2-Flag and SorCS2-FL treated with chloroquine or dynasore for 24 hours (see **Fig. 3a-b**). The data points correspond to four independent biological experiments, $n = 4$. Values are normalized to GAPDH but not transformed into comparable fold change. Mixed-effects model ($P_{\text{Treatment}} = 0.0001$, $P_{\text{Expression}} = 0.0166$) with multiple comparison Fisher's LSD test. **(b)** Immunoblot analysis of ubiquitination assay in HEK293 cells co-expressing His-tagged ubiquitin, Ror2-Flag, and SorCS2-FL, followed by His-specific pulldown. No ubiquitination enrichment (seen as a protein smear in the IBs lines) was observed in any of the immunoblots including the IB:Flag. the experiment was performed in two independent biological experiments. **(c-g)** Representative confocal images showing the co-localization of Ror2-Flag and SorCS2-FL together with overexpressed constructs encoding fluorescently labeled sorting markers. **(c)** Ror2-SorCS2 complex co-localized with the early endosome marker Rab5a-GFP, with the early and recycling endosome marker Rab4-GFP **(d)**, and with the recycling endosome marker Rab11-GFP **(e)**. **(f)** Ror2-SorCS2 complex displayed minimal co-localization with the late endosome marker Rab7-dsRed. **(g)** Ror2-SorCS2 complex showed strong co-localization with the lysosomal marker Lamp1a-mCherry. Scale bar = 10 μm / 20 μm. Nuclear Hoechst staining in blue. **(c-g)** Experiments were performed in independent biological experiments, n = 6. Source data are provided as a Source Data file.

**Suppl. Fig. S7: Co-localization analysis of the Ror-SorCS2 complex with markers labeling endogenous endosomal and lysosomal vesicles.** Representative confocal images showing the co-localization of overexpressed Ror2-Flag and SorCS2-FL in HEK293T with a panel of endogenous markers labelling endosomal and lysosomal vesicles: early endosome markers EEA1 **(a)** and Rab5a **(b)**, late endosome markers Rab7 **(d)** and Rab9a **(e)**, and late endosome/lysosome marker Lamp2a **(f)**. Ror2-Flag was expressed alone (*left panels*) or co-expressed with SorCS2-FL (*right panels*). Scale bar = 10 μm / 5 μm (cropped images). Nuclear Hoechst staining in blue. All experiments were performed in independent biological experiments, n = 4.

**Suppl. Fig. S8: Co-localization analysis of the Ror-SorCS2 complex with markers labeling endogenous subcellular compartments.** Representative immunofluorescence images showing the co-localization of overexpressed Ror2-Flag and SorCS2-FL in HEK293T with a panel of endogenous markers labelling subcellular compartments: GM-130 **(a)**, Vti1a **(b)**, TGN-46 **(c)**,

Golgin-97 **(d)**, and Mannosidase-2 **(e)**. Ror2-Flag was expressed alone (*left panels*) or co-expressed with SorCS2-FL (*right panels*). Scale bar = 10 μm / 5 μm (cropped images). Nuclear Hoechst staining in blue. All experiments were performed in independent biological experiments, n = 4.

**Suppl. Fig. S9: Co-localization analysis of the Ror-SorCS2 complex with endogenous autophagy markers.** Representative immunofluorescence images showing the co-localization of overexpressed Ror2-Flag and SorCS2-FL in HEK293T with a panel of endogenous autophagy markers: ATG9A **(a)**, ATG16L1 **(b)**, Beclin-1 **(c)**, ATG5 **(d)**, ATG4B **(e)**, and LC3B **(f)**. Ror2-Flag was expressed alone (*left panels*) or co-expressed with SorCS2-FL (*right panels*). Scale bar = 10 μm / 5 μm (cropped images). Nuclear Hoechst staining in blue. All experiments were performed in independent biological experiments, n = 4.

**Suppl. Fig. S10: Control *ror2-sorcs2* microinjections in zebrafish embryos**

**(a)** Evaluation of Ror2 overexpression model at Prim-6 stage. Compared to control (Ctr) and *ror2* sMO1 condition, the injection of Ror2 mRNA did not promote CE defects in zebrafish. Gain of function approach in zebrafish is therefore not a suitable model to assess the regulatory role of SorCS2 on overexpressed Ror2, as established in *Xenopus l.* and human cell cultures. *Sorcs2* overexpression caused some worsening of CE defects. The data are compiled from 10 independent microinjection experiments. **(b)** Prim-6 stage zebrafish embryos injected with control morpholino (ctr MO), *sorcs2*-specific splice-site morpholino (*sorcs2* sMO), *ror2* sMO2, and a combination of *sorcs2* and *ror2* sMOs. Co-injection of *ror2* with *sorcs2* sMOs exacerbated *ror2*-dependent body axis and head defects. The data are compiled from 2 independent microinjection experiments. **(c)** Morpholino microinjections in zebrafish embryos with mixed genetic background (Casper x AB). *Sorcs2* knockdown alone did not cause CE defects at the Prim-6 stage, but the co-injection with

*ror2* sMO2 significantly exacerbated the *ror2*-dependent phenotype. The data are compiled from 5 independent microinjection experiments. **(d)** Quantification of *ror2* sMO2 injection in *WT* and *sorcs2*<sup>-/-</sup> embryos showing aggravated CE defects in mutants. The data are compiled from 2 independent microinjection experiments. **(e)** Injection of *wnt5b* translation-blocking MO (tMO) into *sorcs2*<sup>-/-</sup> embryos did not exacerbate the *wnt5b*-dependent CE defects. The data are compiled from 2 independent microinjection experiments. **(f)** Assessment of *sorcs2* sMO toxicity in mutant *sorcs2*<sup>-/-</sup> embryos at the Prim-6 stage. Injection of ctr MO into *sorcs2*<sup>-/-</sup> embryos slightly increased CE defects compared to *WT*, but the phenotype was not worsened by *sorcs2* sMO, suggesting minimal toxicity and off-target effects. The data originates from one microinjection experiment. Source data are provided as a Source Data file.

**Suppl. Fig. S11: Expression profiling of *ror2*, *sorcs2*, and *wnt5b* during zebrafish embryogenesis.** **(a)** Alkaline phosphatase WISH labeling *sorcs2* transcripts in uninjected *WT* zebrafish embryos during early embryogenesis. *Sorcs2* transcripts were detected as early as the blastula stage (3.5 hpf), with signal intensity gradually increasing throughout the embryo over the first 24 hours. At the Prim-6 stage, *sorcs2* expression was enriched increased in the lens, telencephalon, diencephalon, midbrain, and hindbrain. **(b-e)** Fluorescent multi-probe WISH using HCR3.0 labeling *ror2*, *sorcs2*, and *wnt5b* transcripts in uninjected *WT* zebrafish embryos during convergent extension and somitogenesis. *Sorcs2* was consistently co-expressed with *ror2* across all analyzed stages. **(b)** At the 50% epiboly stage, *sorcs2*, *ror2*, and *wnt5b* were co-expressed throughout the embryo. Scale bars: 100 μm / 20 μm in the cropped images. **(c)** At the bud stage, *sorcs2* was co-expressed with *ror2* throughout the embryo, while *wnt5b* transcripts were exclusively localized to the tail, suggesting the existence of an alternative signaling axis. Scale bars: 200 μm / 20 μm in the cropped images. **(d)** At the 7-somite stage, *sorcs2* was co-expressed with *ror2* throughout the embryo, including the head and segmented somites. At this stage, *wnt5b* transcripts remained absent in the head. Scale bars: 200 μm / 20 μm in the cropped images. **(e)** At the 22-somite stage, *sorcs2*, *ror2*, and *wnt5b* were co-expressed in the brain, particularly in the forebrain and midbrain. Scale bars: 100 μm / 20 μm (cropped images). All experiments were performed in three independent biological experiments, analyzing n = 10 embryos per group.

**Suppl. Fig. S12: Expression profiling and function of *ror2*, *sorcs2*, and *wnt5b* during zebrafish brain morphogenesis.** **(a)** At the Prim-6 stage, *sorcs2*, *ror2*, and *wnt5b* were co-expressed in the brain, particularly at the telencephalon-diencephalon boundary (TDB) and the midbrain-hindbrain boundary (MHB). T = telencephalon, D = diencephalon, M = midbrain, H = hindbrain, segmented into rhombomeres (r1-r7). TDB – blue, star, MHB – green, cross. **(b-c)** Comparison of whole-mount RNA *in situ* hybridization methods in *WT* embryonic brain at the Prim-6 stage: alkaline phosphatase WISH **(b)** and fluorescent multi-probe WISH (HCR3.0) visualized via Imaris-based segmentation **(c)**. Both approaches showed consistent *sorcs2*, *ror2*, and *wnt5b* expression, notably at the TDB and MHB (arrows). **(d-f)** Alkaline phosphatase WISH labeling *wnt5b* transcripts at the Prim-6 stage. In the brain, *wnt5b* is prominently expressed in the floorplate, TDB, and MHB. **(e)** *Wnt5b* expression increases in the tail of embryos injected with *ror2* sMO alone or co-injected with *sorcs2* sMO. In contrast, while single *ror2* or *sorcs2* sMO injections do not affect *wnt5b* expression in the brain, co-injection nearly abolishes *wnt5b* expression within brain boundaries (arrows). **(f)** Representative *sorcs2*<sup>-/-</sup> embryos injected with control MO or *ror2* sMO1. *ror2* knockdown in *sorcs2*<sup>-/-</sup> embryos led to loss of *wnt5b* expression at one or both brain boundaries (arrows). **(g)** Representative images of multiplex fluorescent WISH (HCR v3.0) detecting *fgf8* and *wnt5b* transcripts for 3D analysis of MHB morphology. Prim-6 stage zebrafish embryos were injected with control morpholino (ctr MO), *sorcs2*-specific splice-site morpholino (*sorcs2* sMO), *ror2* sMO1, and a combination of *sorcs2* and *ror2* sMOs. Confocal Z-stacks. Scale bars: 80 μm (lateral views), 50 μm (coronal views). The images were quantified in (Fig. 5m-p). All experiments were performed in three independent experiments, assessing $\geq 10$ embryos per group.

**Suppl. Fig. S13: Macro-anatomical measurements of *Ror2*<sup>+/-</sup>; *Sorcs2*<sup>+/-</sup> double heterozygote breeding embryos at E12.5.** (a-c) Body measurements of adult *Sorcs2*<sup>+/+</sup> and *Sorcs2*<sup>-/-</sup> mice at 9-10 weeks of age. (b) *Sorcs2*<sup>-/-</sup> mice exhibit a shorter anterior-posterior body axis compared to *Sorcs2*<sup>+/+</sup>, with no significant impact on limb length. Measurements include both sexes (males: WT = 4, KO = 5, females: WT = 4, KO = 3). Two-tailed multiple unpaired t-tests with Welch's correction. (c) Weight measurements of adult *Sorcs2*<sup>+/+</sup> and *Sorcs2*<sup>-/-</sup> littermates at 9-10 weeks old. *Sorcs2*<sup>-/-</sup> mice exhibit reduced weight, irrespective of gender. Two-tailed multiple unpaired t-tests with Welch correction. (d) Representative bright-field images of live littermate embryos that were used in (Fig. 6a-f) for macro-anatomy measurement of double-heterozygote breeding embryos at E12.5. The measured samples comprised of 47 quantified embryos originating from 6 independent breeding females. Source data are provided as a Source Data file.

**Suppl. Fig. S14: Macro-anatomical measurements of *Ror2*<sup>+/-</sup>; *Sorcs2*<sup>+/-</sup> double heterozygote breeding embryos at E14.5.** (a) Representative bright-field images of live embryos for macro-anatomy measurements of E14.5 embryos from *Ror2*<sup>+/-</sup>; *Sorcs2*<sup>+/-</sup> double-heterozygote breeding. (b) A representative brightfield photo of a live E12.5 embryo with highlighted features that were measured. The samples comprised of 48 quantified embryos (littermates) originating from 9 breeding females. (c-e) Quantification of anterior-posterior (A-P) proportions of E14.5 littermates. (c) Body axis length measured along the dorsal side of the embryo. Ordinary one-way ANOVA (P = 0.0012) with Fisher's LSD test. (d) Trunk length. Ordinary one-way ANOVA (P = 0.0029) with

Fisher's LSD test. **(e)** Brain length measured along the dorsal side of the brain. Ordinary one-way ANOVA ($P = 0.1408$) with Fisher's LSD test. **(f)** Forebrain-midbrain thickness measured as a straight line between forehead and dorsal midbrain. Ordinary one-way ANOVA ($P = 0.3039$) with Fisher's LSD test. **(g-h)** Quantification of craniofacial features in respect to length **(g)** and width **(h)** of naso-maxillary area. Ordinary one-way ANOVA ($P_{\text{length}} = 0.0011$, $P_{\text{width}} = 0.0018$) with Fisher's LSD test. **(i-j)** Limbs measurements. Ordinary one-way ANOVA ($P_{\text{forelimb}} = 0.0001$, $P_{\text{hindlimb}} = 0.00009$) with Fisher's LSD test. **(k)** Quantification of eye diameter. Ordinary one-way ANOVA ($P = 0.3770$) with Fisher's LSD test. Data are presented as mean ± S.E.M. Source data are provided as a Source Data file.

**Suppl. Fig. S15: Expression profiling using accessible single-cell RNA sequencing of mouse ventral midbrain. (a-g)** Extracted expression profiles from published single-cell RNA sequencing data of developing mouse VM (*La Manno and Gyllborg et al*)<sup>2</sup>. The grey shading indicates the statistically significant enrichment of expression in specific cell types. *Note*: expression profiles cannot be directly extracted from the explicit tissue age. **(a)** Cell type cluster classification from embryonic to postnatal VM development. **(b)** *Ror2* is expressed across all clusters, with higher levels observed in progenitor and neuroblast subtypes. **(c)** *Sorcs2* expression is enriched in radial glia and mediolateral neuroblasts. **(d)** *Slc1a3* (encoding Glast/EAAT1 protein) expression is specific for radial glia subtypes and ependymal cells. **(e)** *Wnt5a* expression is enriched in radial glia, mDa neuroblasts, and mDa neurons. **(f)** *Ror1* transcripts were not detected using this method. **(g)** *Tcf4* expression is enriched in neuronal progenitors, radial glia subtypes, and dopaminergic neuroblasts, and additional neuroblast subtypes. **(h)** Allen Brain atlas in situ hybridization staining shows predominal expression of *Tcf4* in the ventricular zone of midbrain both at E11.5 and E13.5. This area mostly contains neural progenitors including mDa radial glia.

**Suppl. Fig. S16: Expression profiles of *Ror2* and *Sorcs2* in *WT* ventral midbrain using fluorescent multiplex RNA-ISH.** Fluorescent multiplex RNA-ISH was performed on coronal sections of embryonic ventral midbrain (VM) with Hoechst nuclear staining (blue). **(a)** *Ror2* and *Sorcs2* were co-expressed in *Lmx1a*<sup>+</sup> mDa progenitors at E10.5. Scale bars: 100 μm /20 μm (cropped). **(b-c)** RNA-ISH combined with immunofluorescence labeling for the radial glia marker Glast at E10.5 **(b)** and E11.5 **(c)**. At both stages, *Ror2* and *Sorcs2* were co-expressed by newborn Glast<sup>+</sup> mDa progenitors, with their cell count gradually increased at E11.5. Scale bars: 50 μm /10 μm (cropped). Experiments were performed in three independent biological experiments.

**Suppl. Fig. S17: Subcellular fractionation of SN4741 cell lines.** Protein analysis of Wnt signaling components in control (*Ctr*) and *Sorcs2* KO monoclonal mDa progenitor lines (SN4741) using subcellular fractionations and immunoblotting. The band intensity was normalized to GAPDH and *Ctr* samples within each fraction. Homog. = Homogenate/input lysate. PM = Plasma Membrane. *cl3/6* = clone 3/6. Two-tailed multiple unpaired t-tests. **(a)** SorCS2 immunoblotting confirming purity of the cytosol fraction and validation of the *Sorcs2* KO. **(b)** GAPDH immunoblotting was used as a loading and fractionation control. **(c)** Total Dvl3 protein levels (see **Fig. 8d-e**). **(d-e)** Total Lrp6 protein levels. **(f-g)** Detection and quantification of phosphorylated Lrp6 at Ser1490. All graphs represent mean ± S.E.M. Five independent experiments with 1-2 biological replicate/genotype. Source data are provided as a Source Data file.

**Suppl. Fig. S18: Analysis of Wnt signaling components in mDa progenitor CRISPR/Cas9 lines treated with Wnt3a and Wnt5a.** **(a)** Representative immunoblots of monoclonal control (*Ctr*), *Ror2* KO, *Sorcs2* KO, and *Ror2*; *Sorcs2* dKO mDa progenitor lines generated via CRISPR/Cas9 in SN4741 cells. Cells were treated with mock (PBS + 0.1% BSA), recombinant human Wnt5a (100 ng/ml), or Wnt3a (100 ng/ml) for 2 hours (acute stimulation) or 16 hours

(chronic stimulation). **(b-g, i)** Quantifications of the immunoblots. Two independent CRISPR clones per genotype were pooled from 3-5 independent experiments and statistically tested using two-way ANOVA with Fisher's LSD test **(b-g)**. Acute and chronic Wnt3a stimulation amplified canonical Wnt/β-catenin signaling consistently in all lines based on Lrp6 phosphorylation **(b)**, Dvl3 phosphorylation **(c-d, f)**, and dephosphorylated (active) β-catenin **(e, g)**. Acute **(c-d)** and chronic **(e)** Wnt5a stimulation amplified noncanonical Wnt signaling consistently in all lines based on Dvl3 phosphorylation. **(d)** As observed for the 16h treatment **(Fig. 9n)**, *Sorcs2 KO* and *Ror2*; *Sorcs2 dKO* lines exhibited increased Dvl3 phosphorylation in mock, indicating enhanced basal Wnt signaling in the absence of SorCS2. **(h)** Assessing Wnt5a-dependent electrophoretic mobility of phosphorylated Ror2 using Sigma antibody. **(i)** Increased Ror2 phosphorylation in *Sorcs2 KO* was evident already during acute Wnt5a stimulation. Four independent experiments. All graphs represent mean ± S.E.M. Source data are provided as a Source Data file.

**Suppl. Fig. S19: Original uncropped immunoblots presented in the Suppl. Fig. S1**

**Suppl. Fig. S20: Original uncropped immunoblots presented within the Suppl. Fig. S3**

**Suppl. Fig. S21: Original uncropped immunoblots presented within the Suppl. Fig. S4**

**Suppl. Fig. S22: Original uncropped immunoblots presented within the Suppl. Fig. S6**

**Suppl. Fig. S23: Original uncropped immunoblots presented within the Suppl. Fig. S17**

**Suppl. Fig. S24: Original uncropped immunoblots presented within the Suppl. Fig. S18**

**Suppl. Video SV1: Light-sheet microscopy of E12.5 *Hb9:GFP/+*; *Sorcs2*<sup>+/+</sup> embryo and 3D**
**reconstructions and quantification of selected areas of interest.** Transgenic *Hb9:GFP*
highlights all motor neurons, enabling the segmentation and measurement of the spinal cord using
Imaris software. Forelimb and hindlimb buds were segmented based on their natural
autofluorescence and measured accordingly. Related to **Fig. 6i-l.**

**Suppl. Video SV2: Confocal imaging and 3D reconstructions of EEA1+ sorting vesicles in**
**tissue sections and mDa lineage.** Related to **Fig. 9c-e.**

**Suppl. Table S1: Comprehensive list of all proteins detected across all IP-MS/MS**
**experiments.** Related to **Fig. 1.**

**Suppl. Table S2: Filtered list of Ror2-specific proteins identified in IP-MS/MS experiments,**
**with or without Wnt5a treatment.** Related to **Fig. 1.**

**Suppl. Table S3: Summary of candidate Ror2 binding partners consistently detected across**
**IP-MS/MS datasets.** Related to **Fig. 1.**

**Suppl. Table S4: Annotated list of known Ror2 binding partners and Wnt signaling**
**components identified across all IP-MS/MS datasets.** Related to **Fig. 1.**

**Suppl. Table S5: Detailed results of DAVID GO analysis for “Biological Process”.** Related
**to Fig. 1.**

**Suppl. Table S6: Detailed results of DAVID GO analysis for “KEGG Pathway”.** Related to
**Fig. 1.**

**Suppl. Table S7: Detailed list of antibodies used in this study.**

282   **References:**

1. La Manno G, *et al.* Molecular architecture of the developing mouse brain. *Nature* **596**, 92-96
(2021).

2. La Manno G, *et al.* Molecular Diversity of Midbrain Development in Mouse, Human, and Stem
Cells. *Cell* **167**, 566-580.e519 (2016).
